# NetRAX: Accurate and Fast Maximum Likelihood Phylogenetic Network Inference^⋆^

**DOI:** 10.1101/2021.08.30.458194

**Authors:** Sarah Lutteropp, Céline Scornavacca, Alexey M. Kozlov, Benoit Morel, Alexandros Stamatakis

## Abstract

Phylogenetic networks are used to represent non-treelike evolutionary scenarios. Current, actively developed approaches for phylogenetic network inference jointly account for non-treelike evolution and incomplete lineage sorting (ILS). Unfortunately, this induces a very high computational complexity. Hence, current tools can only analyze small data sets.

We present NetRAX, a tool for maximum likelihood inference of phylogenetic networks in the absence of incomplete lineage sorting. Our tool leverages state-of-the-art methods for efficiently computing the phylogenetic likelihood function on trees, and extends them to phylogenetic networks via the notion of “displayed trees”. NetRAX can infer maximum likelihood phylogenetic networks from partitioned multiple sequence alignments and returns the inferred networks in Extended Newick format.

On simulated data, our results show a very low relative difference in BIC score and a near-zero unrooted softwired cluster distance to the true, simulated networks. With NetRAX, a network inference on a partitioned alignment with 8, 000 sites, 30 taxa, and 3 reticulations completes within a few minutes on a standard laptop.

Our implementation is available under the GNU General Public License v3.0 at https://github.com/lutteropp/NetRAX.

## 1 Introduction

It is not always possible to describe evolution via a phylogenetic tree. Events such as horizontal gene transfer, hybridization, or recombination induce non tree-like evolutionary relationships among taxa. In such cases, a phylogenetic network better describes evolutionary relationships. A phylogenetic network differs from a phylogenetic tree by having nodes with two parents, the so-called *reticulations*, in addition to regular tree nodes.

Initially, phylogenetic network inference methods based on maximum likelihood (ML) did not account for incomplete lineage sorting (ILS). For instance, the NEPAL tool [19] does not account for ILS in its network likelihood model. This tool starts from a phylogenetic tree and adds reticulations while keeping the underlying backbone-tree topology fixed. The PhyloDAG tool [20] does also not account for ILS. It implements an efficient expectation-maximization algorithm to infer ML networks using the mixture model of [27]. Very recently, a new ML network inference tool disregarding ILS, PhyLiNC, has been proposed [1]. The tool is available as part of the PhyloNetworks [26] package, although it is deactivated by default and the authors emphasize that it is not ready to be used yet [2]. Unfortunately, NEPAL and PhyloDAG also appear to not be in a state for routinely reconstructing phylogenetic networks from genomic data, as we elaborate in Section 3.4.

In the past years, the focus shifted towards developing methods for ML network inference that also account for ILS. While models accounting for ILS are expected to yield more accurate networks as they incorporate additional mechanisms to explain non-treelike evolution, they face substantial computational challenges. For example, the ILS-aware ML method implemented in PhyloNET [29] can only be applied to very small data sets with 10 taxa and up to 4 reticulations [25]. Consequently, faster-to-compute pseudolikelihood models accounting for ILS, such as implemented in the SNaQ tool [25], were developed. These pseudolikelihood models first compute ILS-aware likelihoods of 4-taxon subnetworks (quartets) on gene trees, and subsequently compute a pseudolikelihood for the entire network from these quartet likelihoods. More recent versions of PhyloNET also deploy pseudolikelihoods [29], but still face scalability challenges [4]. Apart from ML based network inference tools, there also exist tools that rely on maximum parsimony, e.g., [18], which often perform poorly [12], or Bayesian inference, e.g., [30], which face substantial scalability challenges.

In this paper we present NetRAX, a tool for ML inference of phylogenetic networks that does not account for ILS. By leveraging state-of-the-art methods for efficiently computing the phylogenetic likelihood function on trees and extending them to networks, our tool permits to run –on a standard laptop and within minutes– phylogenetic analyses on partitioned alignments for evolutionary scenarios with several reticulations. Based on the performance on simulated data and on an empirical data set NetRAX allows to routinely perform phylogenetic inference in the presence of non-treelike evolution. The remainder of this paper is structured as follows. We introduce the NetRAX likelihood model in Section 2.1 and outline its implementation in Section 2.2. We describe the optimization of branch lengths and other non-topology model parameters in Sections 2.3 and 2.4. In Section 2.5, we outline the available topology-rearrangement moves and in Section 2.6 we present the search algorithm to find the best-scoring network. To compare network topologies, we implement normalized versions of common network distance measures, which we describe in Section 3.1. In Section 3.2, we describe our synthetic data simulation. We detail our experimental setup in Section 3.3. The experimental results on synthetic data are discussed in Section 3.4 and we assess NetRAX results on empirical data in Section 4. We conclude with a discussion of future work in Section 5.

## 2 Materials and Methods

In the following, we provide a general abstract outline of NetRAX. Additional technical details are provided in the Supplementary Material.

### 2.1 Phylogenetic Network Likelihood Model

A binary phylogenetic network *N* is a single-source, directed, acyclic graph. We call its source node the *root* node of *N*. Apart from the root, there are three types of nodes: (i) internal tree nodes with 1 incoming edge and 2 outgoing edges, (ii) reticulation nodes with 2 incoming edges and 1 outgoing edge, and (iii) leaf nodes with one incoming edge and no outgoing edges. Each leaf is associated with a distinct taxon. Each edge *e* in a phylogenetic network has a branch length and a probability *P* (*e*). The incoming edges of a reticulation node (called *reticulation edges*) are assigned inheritance probabilities that must sum to one. The probability of observing a non-reticulation edge is always equal to one. Figure 1 shows a simple example network. Nowadays, typical phylogenetic analyses are conducted on multiple sequence alignments (MSA) 𝒜 that are subdivided into multiple partitions *A*_1_, …, *A*_*p*_. A partition consists of a set of sites that are likely to have evolved together (e.g., sites of a single gene), following the same evolutionary process. With NetRAX, our goal is to infer a phylogenetic network *N* from a partitioned MSA 𝒜 that maximizes a network likelihood *L*(*N*|𝒜).

**Fig. 1.**
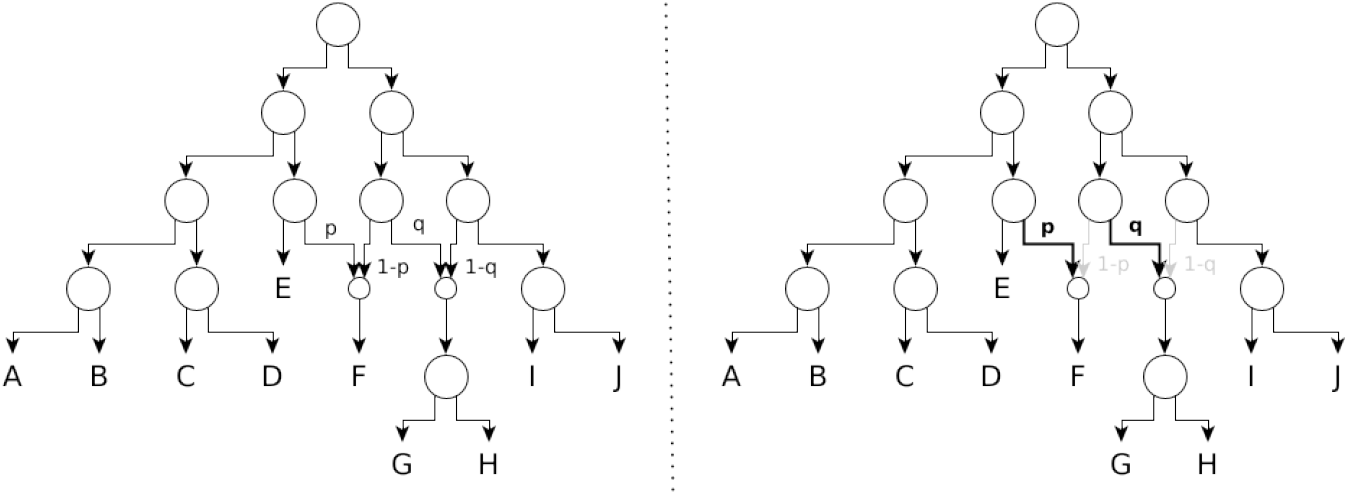
Left: A phylogenetic network with two reticulation nodes. Right: A displayed tree of the phylogenetic network on the left. The probability of displaying the highlighted tree is the product *p* ∗ *q* over the respective reticulation probabilities.

In literature, there exists a plethora of different phylogenetic network definitions. Here, we assume that neither ILS nor recombination occurs among MSA sites. Given these assumptions, we can model reticulate evolution in a network via its set of induced *displayed trees* [16].

We obtain a displayed tree from a network by choosing one parent per reticulation node (disabling the incoming edges belonging to non-chosen parents). Figure 1 shows a displayed tree in a phylogenetic network.

Let *N* = (*V, E*) be a phylogenetic network with a set of displayed trees 𝒯 (*N*). To compute the probability of a displayed tree *T* in 𝒯 (*N*), we proceed as follows. Let *E*_*r*_ be the set of reticulation edges that need to be taken in order to generate *T*. The probability of *T* in *N* is then:

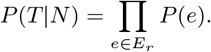

To define *L*(*N*|𝒜), let 𝒜 be a partitioned MSA with partitions *A*_1_, …, *A*_*p*_, that is, every MSA site is assigned to exactly one partition *A*_*i*_ (note that, in case of SNP data sets, each partition contains only one site). Let *ϑ* = (*ϑ*_1_, …, *ϑ*_*p*_) be the parameter vector, storing the per-partition network branch lengths and likelihood model parameters. We consider each partition as being independent. Thus, we define the likelihood of a phylogenetic network given *ϑ* as the product over the per-partition likelihoods:

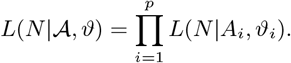

To avoid numerical underflow, we take the logarithm:

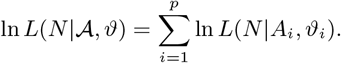

We assess and implement two versions for computing the log likelihood (lnL) on partitioned networks. They both aggregate over the lnL of the trees displayed by the network [16]:

1. **Weighted Average Version (**`LhModel.AVERAGE`):

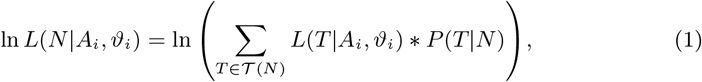
2. **Best Tree Version (**`LhModel.BEST`):

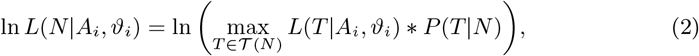

where *L*(*T*|*A*_*i*_, *ϑ*_*i*_) is the standard likelihood function for a tree *T*, given an alignment *A*_*i*_ and parameter vector *ϑ*_*i*_.

In the weighted average version, the likelihood of a network for a given partition is the weighted average over the displayed tree likelihoods. We use the sum here, because the probability of event *A* or *B* to occur is the sum over the probability of observing *A* and the probability of observing *B*. The weighted average can thus be interpreted as the expected value, if we treat each displayed tree as a statistical event.

Because we need to compute the logarithm of a sum over very small numbers and we need to exponentiate the tree lnLs, we use arbitrary-precision arithmetics (using MPFR C++ library [13]) to compute *L*(*N*|𝒜_*i*_) from the per-partition displayed tree likelihoods. As in the RAxML-NG [17] tool for ML phylogenetic tree inference, we use the libpll [9] and pll-modules [6] to compute displayed tree lnLs via the standard Felsenstein pruning algorithm [7].

Currently, NetRAX supports two branch length models. Under the linked branches model, we share the same set of branch lengths among *all* partitions. Under the unlinked branches model, each partition has its own independent set of branch lengths. The branch length model choice has an effect on the type of reticulations we can recover. Figure 2 shows an example network with a reticulation that can not be recovered under the unlinked branches model. Thus, by default, we use linked branch lengths in NetRAX.

**Fig. 2.**
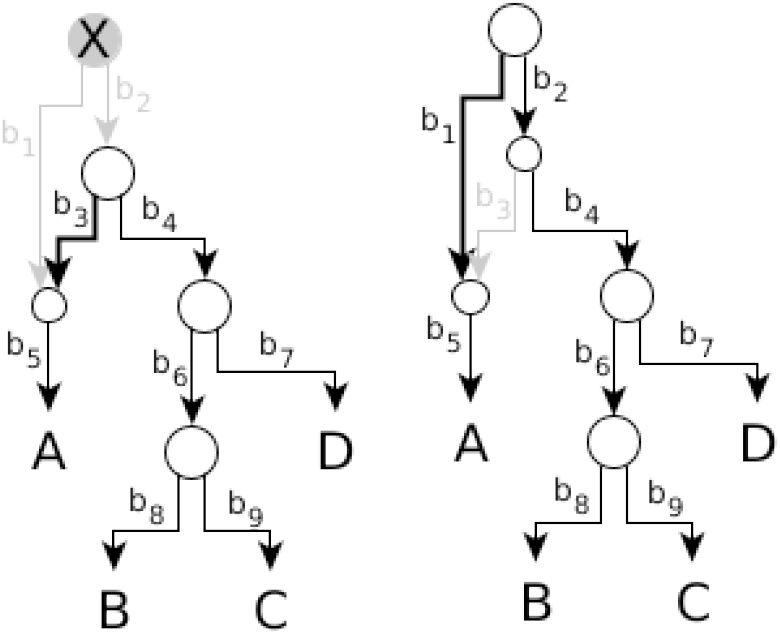
Two displayed trees in a phylogenetic network. Both displayed trees induce the same topology after collapsing single-child nodes. They only differ in some branch lengths. For example, in the left tree, the branch length between the root node and leaf A is *b*_3_ + *b*_5_, and in the right tree it is *b*_1_ + *b*_5_.

### 2.2 Computing the Likelihood of a Phylogenetic Network

In order to compute the lnL of a network *N* using the formulas from the previous section, we first need to compute the per-partition likelihoods *L*(*T*|*A*_*i*_, *ϑ*_*i*_)_*i*∈{1,…,*p*}_ and the probabilities *P* (*T*|*N*) of all trees *T* ∈ 𝒯 (*N*) displayed by *N*.

We use the libpll library to compute ln *L*(*T*|𝒜_*i*_, *ϑ*_*i*_). To compute the per-partition lnLs of a phylogenetic tree, libpll uses an internal per-node data structure, called *conditional likelihood vector* (CLV) [8] that was introduced by Joe Felsenstein [7]. A CLV for a node *v* stores the per-site likelihoods for the subtree rooted at *v* (see Figure 3). The libpll library computes the per-node CLVs via a post-order traversal of the tree using the Felsenstein pruning algorithm [7]. It computes the CLV of a given node based on the CLVs of its respective children. The libpll library also provides incremental likelihood computations: it only updates those CLVs affected by a topological rearrangement move or branch length change and re-uses unaffected CLVs that are still valid.

**Fig. 3.**
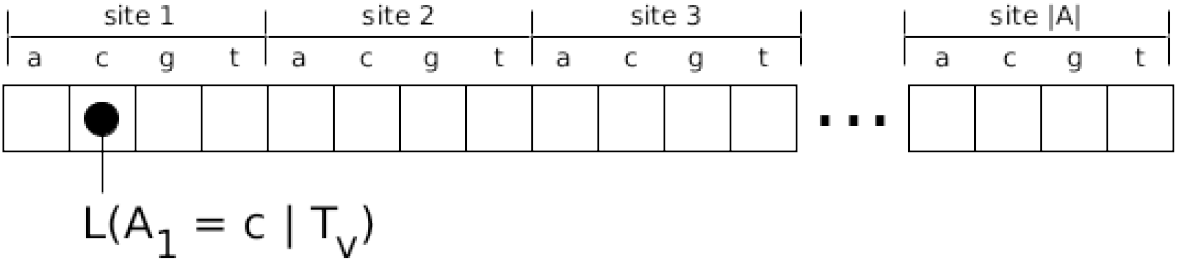
A conditional likelihood vector (CLV) for a subtree *T*_*v*_ rooted at a node *v* in a phylogenetic tree *T* and a MSA partition *A*.

In NetRAX, we do not store each displayed tree topology separately, but use a network data structure that implicitly induces each tree. This allows us to avoid redundant CLV computations: When two displayed trees share an identical subtree, there is no need to compute the CLVs for nodes in this subtree more than once (see below).

We parallelize per tree lnL computations over the MSA sites by using MPI. For this, we reuse the load balancing solutions provided by RAxML-NG.

#### Sharing CLVs among Displayed Trees

Naïvely, by explicitly iterating over each displayed tree in a phylogenetic network *N* with *n* nodes and *r* reticulations, one would require *n* ∗ 2^*r*^ CLVs to calculate the lnL of the network: one CLV per node and displayed tree. To improve efficiency, for each node *v* in *N*, we store as many CLVs as there are different displayed subtree topologies rooted at *v*. By sharing CLVs among identical subtrees in multiple displayed trees, we reduce the total number of CLVs required to compute the lnL of this network.To implement this CLV sharing optimization, we update the CLVs via a bottom up traversal (reversed topological sort) of the nodes in the phylogenetic network. For each node *v* we visit, we update the CLVs for each of the distinct displayed subtree topologies present at *v*.

Due to the peculiarities of networks, multiple, predominantly technical, modifications to the network traversal and displayed tree data structure are required that are described in the supplement.

In the following, we describe how we optimize the network topology *N* and its associated parameter vector *ϑ* to maximize ln *L*(*N*|𝒜, *ϑ*).

### 2.3 Branch Length Optimization

Our goal is to optimize a branch length *b* in a network *N*, with respect to the lnL of the network. Overall, we aim at finding the branch length assignments 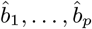 that maximize ln 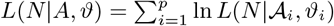.

As in standard ML implementations for tree inference, we optimize *b* via the Newton-Raphson method. For this, we need the first and second derivatives of the network lnL with respect to *b*. We derive formulas for efficiently computing (ln *L*(*N*|*A, ϑ*))′ and (ln *L*(*N*|*A, ϑ*))″ from ln *L*(*T*|*A*_*i*_, *ϑ*_*i*_), (ln *L*(*T*|*A*_*i*_, *ϑ*_*i*_))′, and (ln *L*(*T*|*A*_*i*_, *ϑ*_*i*_))″ in the supplementary text. Note that, when optimizing a branch length *b*, we neither need to recompute the per-partition lnL nor its derivatives for displayed trees not containing *b*.

In the following, we describe the efficient computation of the per-partition lnL and the per-partition lnL derivatives for all displayed trees *T* ∈ 𝒯 (*N*).

#### Branch-Length Optimization in Phylogenetic Trees

In order to avoid costly CLV updates when optimizing a branch (*u, v*) in a tree, all modern ML tree inference tools re-root the tree at the node *u* before optimizing the branch. After re-rooting the tree, the edge directions in subtrees rooted at nodes that lie on the path between the old and the new root (both ends included) change. We thus need to recompute the CLVs for the nodes residing on this path. When evaluating different values for the branch length (*u, v*) in the re-rooted tree, we can reuse the CLVs stored at *u* and *v* without having to recompute them. Note that, the phylogenetic likelihood is the same regardless of the root placement under the commonly used time-reversible models of evolution.

#### Re-rooting Displayed Trees in a Phylogenetic Network

In NetRAX, we deploy an analogous strategy for efficient branch length optimization as used for phylogenetic trees. For this, before optimizing a branch (*u, v*), we need to re-root *all* displayed trees (where the branch is present) at the source node *u*.

Recall that we do not explicitly store each displayed tree topology in order to avoid redundant CLV updates during the network lnL computation. Instead, we perform a custom bottom-up traversal of the nodes in the phylogenetic network, updating all unique subtree CLVs shared among subsets of displayed trees. This complicates the re-rooting operation for the displayed trees.

The difficulty here is that with the original network root, the edge directions (and thus, the parent-child relationships needed for computing CLVs) are identical for all displayed trees. But when re-rooting the displayed trees, the parent-child relationships depend on the specific displayed tree we are currently considering (see Figure 4). These differing edge directions affect the order in which we need to process the nodes when recomputing the CLVs. For networks, we need to recompute the CLVs (which are shared among subsets of displayed trees) that lie on *any* path between the old root and the new root (both ends included). We devised the following approach for resolving the edge direction problem: We process the paths between the network root and the new root successively. Before processing the next path, we invalidate the shared CLVs at all nodes on the current path, except for the new root node. We detect, in advance, at which nodes we need to restore the old shared CLVs (to the values they had when using the original root), save, and restore them accordingly. After we finished optimizing a branch, we recompute the CLVs with regard to the original network root. This means that, when successively optimizing *k* branches, we need to re-root the displayed trees in the network as follows: new_root_1 → old_root → new_root_2 → old_root → … new_root_k → old_root. We do this because, in networks, unlike phylogenetic trees, the root placement *does* influence the lnL value.

**Fig. 4.**
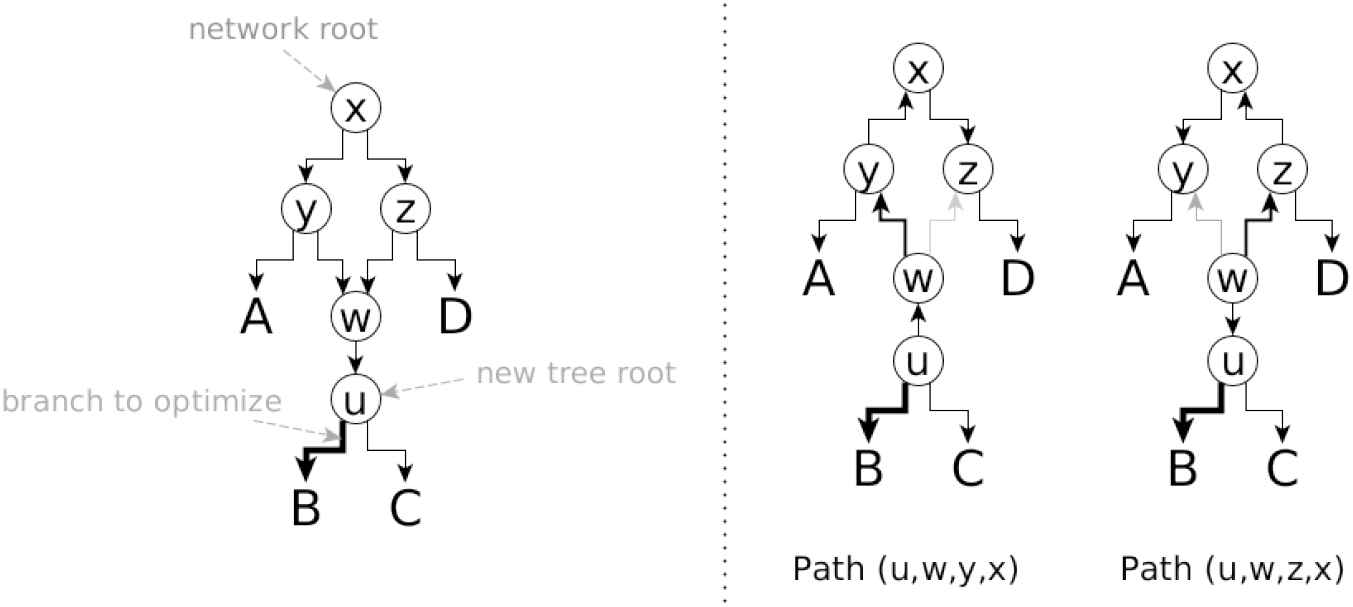
We re-root the displayed trees at node *u* before optimizing branch (*u, v*). In the first re-rooted displayed tree, the node y is a parent of node x and we need to recompute the CLVs on the path (u,w,y,x). However, in the second re-rooted displayed tree, the node y is a child of node x and we need to recompute the CLVs on the path (u,w,z,x).

### 2.4 Optimizing Non-Topology Parameters

Apart from optimizing branch lengths, we also need to optimize the evolutionary model parameters (by default, we use the GTR+*Γ* [28] model, although NetRAX supports all models that are supported by RAxML-NG) and reticulation probabilities. Recall that our goal is to optimize the overall network lnL. This is, we aim to find optimal parameters for the parameter vector *ϑ*, to maximize 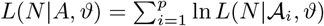.

#### Likelihood Model Parameter Optimization

We reuse routines from RAxML-NG for optimizing these parameters optimizations. As these methods do not rely on an explicit tree topology, we can directly reuse them for networks.

#### Optimizing Reticulation Probabilities

For optimizing the first-parent probability *p* of a reticulation, we also reuse Brent’s single-parameter optimization method as implemented in RAxML-NG. Since the first and second parent probability of a reticulation sum to 1.0, the probability for taking the second parent follows from the first parent probability.

The Brent optimization method requires us to recompute the network lnL when *p* changes. Fortunately, per-partition lnLs of the displayed trees do *not* depend on *p*. Changing *p* only affects the probabilities *P* (*T*|*N, ϑ*)_*T*∈𝒯(*N*)_ of displaying the trees. We can thus re-use the existing (already computed) per-partition lnLs for displayed trees when recomputing the network lnL during the optimization of *p*.

### 2.5 Supported Topology-Rearrangement Moves

NetRAX implements the following rooted network topology rearrangement moves proposed by Gambette *et al*. [10], as well as reversal (undo) operations for them: rNNI move, rSPR move, arc insertion move, and arc removal move. When undoing a move, we restore the original topology and branch lengths. Doing or undoing a move also invalidates some CLVs. Evidently, there exists a trade-off between recomputing the invalidated CLVs versus storing them. In our current implementation, we simply recompute the CLVs to reduce code complexity and because this strategy has proved to be efficient in RAxML-NG.

#### Comparing Networks of different Complexity

NetRAX supports vertical topology-rearrangement moves that increase (arc insertion move) or decrease (arc removal move) the number of reticulations in a network. Because model complexity changes when adding or removing reticulations from a network, we cannot compare networks of different complexity directly via their respective lnL. For this, NetRAX implements AIC, AICc, and BIC scoring. By default, NetRAX uses the BIC score to compare different networks, since [22] showed that using BIC performs best in network searches.

The BIC score of a network *N* with *r* reticulations on a partitioned MSA 𝒜 and parameter vector *ϑ* is defined as follows: BIC(*N*|𝒜,*ϑ*) = −2 ∗ ln *L*(*N*|𝒜,*ϑ*) + # free_parameters ln(sample_size). The free parameters ∗ are the substitution model parameters, the reticulation first-parent probabilities, and the branch lengths. The sample size is the product of the number of taxa and the number of MSA sites.

### 2.6 Network Search

NetRAX uses a greedy hill climbing approach to search for network topologies. It deploys an outer search loop to iterate over different move types (see Section 2.5) and an inner search loop to search for the best-scoring network using a specific move type. We provide an overview in Figure 5 and detailed pseudocode in the supplement.

**Fig. 5.**
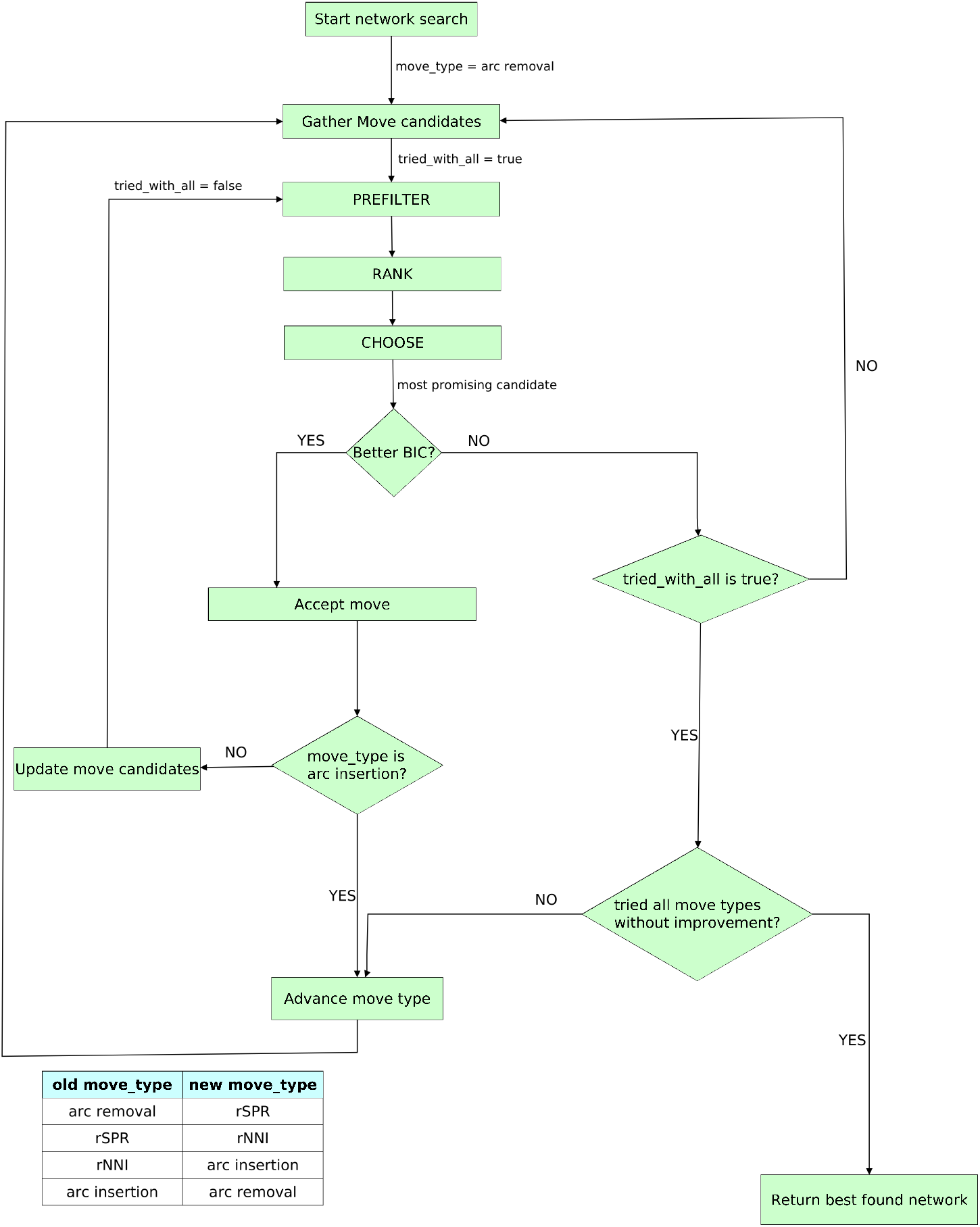
Overview of the NetRAX network search algorithm for a single start network.

In the outer search loop, we search in waves, by repeatedly iterating over move types in the following order: arc removal, rSPR, rNNI, arc insertion. For each move type, we invoke an inner search loop (Section 2.6). The outer search loop terminates when no move type improves the BIC score.

#### Starting Networks

NetRAX can initiate a network search from a given set of start networks provided in Extended Newick format. For example, it can be launched on a user-specified number of strictly bifurcating random and maximum parsimony trees or a best-known ML tree that can be generated, for instance, with RAxML-NG [17] in a separate step.

#### Inner Search Loop

As already mentioned, the inner search loop searches for the best-scoring network using a single move type only.

#### Assembling the set of Move Candidates

We can reach multiple alternative network topologies by applying a single move of a given type to the current network. We call such a move a *move candidate*.

We build the set of move candidates for a specific move type by iterating over all nodes in the network. For each node, we add the move candidates for the current node to the set. When assessing possible move candidates for rSPR or arc insertion moves, NetRAX uses a default search radius of 5. This is, for each node in the network, we only consider and evaluate move candidates within radius 5 around the current node. Due to their smaller neighborhood size, we do not restrict the search radius for rNNI or arc removal moves.

#### Filtering Move Candidates

In order to determine the most promising move candidate(s) and accelerate move evaluation, we apply a three stage pre-filtering process inspired by analogous heuristics used in RAxML-NG: We filter the move candidates using the PREFILTER, RANK, and CHOOSE stages. These stages differ in the number of branches we optimize before scoring each candidate:

– PREFILTER – Do not optimize branch lengths. (Exception: For arc insertion moves, we do need to optimize the length of the newly introduced branch, because we do not have a reasonable initial guess for it.)
– RANK – Optimize branches directly affected by the move.
– CHOOSE – Optimize all branches in the network.

We use the Elbow Method (see supplement) to determine the number of promising move candidates to keep after each filtering stage. The most promising move candidate is the one with the lowest (=best) BIC score after the CHOOSE stage.

#### Accepting a Move and updating the set of Move Candidates

If the most promising move candidate obtained by the CHOOSE stage yields a better-scoring network, we accept the move and apply it to the current network.

When accepting a move, we optimize all branch lengths, reticulation probabilities, and remaining model parameters in this new best network.

If the inner search loop executes arc insertion moves, it terminates after accepting a move and immediately returns to the outer search loop. This is done in order to reduce the time spent optimizing a network with a too high reticulation count. For all other move types, after accepting a move, we continue searching for score-improving moves of the same type until we cannot find a better-scoring network by considering other promising candidate moves from the PREFILTER phase. First, we remove previous promising moves that have become inapplicable after accepting the current move, and add new move candidates to the set, that are seeded at nodes directly affected by the accepted move. When we do not find a better-scoring network by searching this modified set of candidate moves, we again consider the complete set of move candidates. If these do also not yield a better-scoring network, the inner search loop terminates.

## 3 Simulation Study

### 3.1 Topology-based evaluation metrics

For comparing network topologies, we implement normalized versions of the following metrics [14]: (i) softwired cluster distance, (ii) hardwired cluster distance, (iii) displayed trees distance, (iv) tripartition distance, (v) nested labels distance, and (vi) path multiplicity distance. Distance measures (i), (ii), and (iii) have both unrooted and rooted versions.

Due to space constraints, we only discuss and report unrooted softwired cluster distances (SCD) here, and discuss the remaining measures in the supplement. Since the SCD is based on the topologies of displayed trees, it alleviates effects caused by network identifiability issues, where different networks can induce the same set of displayed trees [21]. Further, it resembles the Robinson-Foulds distance [24] on trees, yielding it easy to interpret. We choose the unrooted version, as NetRAX does not explicitly infer a network root.

Every edge in a tree induces a *bipartition*, since its removal splits the set of taxa into two subsets. Bipartitions induced by edges leading to leaf nodes are *trivial*, as they are present in any tree on the given set of taxa.

*(Normalized) Unrooted Softwired Cluster Distance* For a given network *N*, let 𝒯(*N*) be the displayed trees of *N* and let *B*(*N*) be the set of all nontrivial bipartitions of the unrooted trees in 𝒯(*N*). The unrooted softwired cluster distance between two networks *N*_1_ and *N*_2_ is:

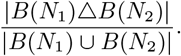

### 3.2 Simulation of Phylogenetic Networks and Sequences

Our simulator is included in the NetRAX GitHub repository. We simulate networks under the birth-hybridization process of [30] with the following parameter set *λ* ∼ U(0,20) + 5, *ν* = *λ*\* 0.003, *τ*_0_ ∼exp(20) + 0.1, where *λ* is the speciation rate, *ν* the hybridisation rate, and *τ*_0_ is the overall time the process is run. The parameters have been chosen to obtain a reasonable amount of reticulations with respect to the taxon range we intend to study. Unless stated otherwise, we set reticulation probabilities to 0.5 for all reticulations in the network.

We repeat our simulations until we obtain a network with the desired number of taxa and number of reticulations. Because our simulator generates ultrametric networks, we discard networks that contain unrecoverable reticulations, that is, networks for which at least two of its displayed trees have the same topology. Note that branch lengths are linked in our simulations.

Subsequently, we simulate sequences for each displayed tree of the simulated network, and concatenate them into a partitioned MSA (using one partition per displayed tree), which is the input of NetRAX. We simulate the sequences using Seq-Gen-1.3.4 [23] with the following parameters: `-mHKY -t3.0 -f0.3,0.2,0.2,0.3`. Although NetRAX supports the GTR model (remember that HKY85 is nested within GTR) we simulated under HKY85 as some of the competing tools we assessed only support HKY85.

The partition length for each displayed tree is proportional to the probability of displaying the tree. By default, we simulate 2^*r*^ ∗ 1000 MSA sites in total, where *r* is the number of reticulations in the network. We do not draw the number of MSA sites from some distribution to keep the data sets more comparable and the results easier to interpret.

### 3.3 Experimental Setup

We conducted extensive experiments on simulated data. For each experiment, we report (i) the number of reticulations in the inferred network, (ii) the relative BIC-score difference between the true and the inferred network, (iii) the unrooted SCD between the true and inferred network, and (iv) the total inference time. In the supplementary text, we further provide additional plots and tables for relative AIC/cAIC/lnL differences, as well as further topology-based evaluation metrics (rooted SCD, rooted/unrooted hardwired cluster distance, rooted/unrooted displayed trees distance, tripartition distance, nested labels distance, path multiplicity distance).

We evaluated NetRAX with `LhType.AVERAGE` and `LhType.BEST`, under the linked branch lengths model on phylogenetic networks and MSAs simulated as described in Section 3.2, under different settings:

*A1: Multiple Starting Trees* We simulated 50 networks each for (i) 10 taxa and 1 reticulation, (ii) 20 taxa and 2 reticulations, (iii) 30 taxa and 3 reticulations.

*A2:* 40 *Taxa* We simulated 50 networks each for (i) 40 taxa and 1 reticulation, (ii) 40 taxa and 2 reticulations, (iii) 40 taxa and 3 reticulations, (iv) 40 taxa and 4 reticulations.

*B: Reticulation Probability* We simulated 50 data sets with 20 taxa and 1 reticulation each. Here, we varied the first-parent probability of all reticulations to be in {0.1, 0.2, 0.3, 0.4, 0.5}.

*C: Unpartitioned Data* We simulated 50 data sets with 20 taxa and 1 reticulation. In addition to normal inference, we started a second inference where we merged all simulated partitions into a single partition before running the inference.

*D: Scrambled Partitions* We simulated a single data set with 30 taxa and 3 reticulations. Before executing NetRAX with a RAxML-NG ML starting tree, we randomly scrambled the partitions such that *p* ∈ {0%, 10%, 20%, 30%, 40%, 50%, 60%, 70%, 80%, 90%, 100%} of the sites from each partition were randomly reassigned to other partitions. Our model assumes that all sites belonging to a partition evolved together. Hence, we are violating this assumption to assess the stability of NetRAX under model violations.

*E: Different Alignment Size* We simulated a single network with 30 taxa and 3 reticulations. For this network we then simulated {50, 100, 500, 1000, 5000, 10000, 50000, 100000} sites per partition.

*F: Parallel Scalability* We simulated 10 networks with 20 taxa and 3 reticulations each. We simulated 10, 000 MSA sites per displayed tree, resulting in a MSA with 80, 000 sites. We started the NetRAX inferences from RAxML-NG ML starting trees, using {1, 2, 4, 8, 16, 32, 64} MPI processes. We also simulated 1 network with 20 taxa and 3 reticulations and a 800, 000 -site MSA. We started the NetRAX inferences from RAxML-NG ML starting trees, using {16, 32, 48, 64, 80, 96, 112, 128} MPI processes.

*G: Comparison with other tools* We simulated a partitioned 10-taxa 1 reticulation data set with reticulation probability 0.5 and 2000 MSA sites. We inferred a ML tree, as well as two partition trees (one for each partition) with RAxML-NG. We also generated a set of 14 unique tree topologies out of 10 random and 10 maximum parsimony RAxML-NG trees, which we used for calling NetRAX with a set of multiple starting trees.

For each simulated data set, we initiated the NetRAX inference from a RAxML-NG ML tree. In addition, for the data sets in A1, we also started another NetRAX inference using 3 random and 3 maximum parsimony starting trees. We inferred networks using NetRAX, PhyLiNC, PhyloDAG, SNaQ, PlyloNET MPL (maximum pseudo-likelihood), PhyloNET MP (maximum parsimony), and PhyloNet ML. We ran PhyLiNC, SNaQ, and PhyloNET using both 1 and 2 as the maximum number of reticulations. In addition, we also compared the number of inferred reticulations with NetRAX, PhyloNET MP, and PhyloDAG (as these were the fastest tools and performed well on the smaller dataset) on a 20 taxon 2 reticulations data set with reticulation probabilities 0.5, and 4000 MSA sites. We also attempted an inference with NEPAL. However, the tool segfaulted and its authors unfortunately have lost its source code [19]. Note that PhyLiNC is officially not ready for use yet [2], but already available on GitHub.

PhyLiNC, and PhyloDAG operate on unpartitioned data, whereas NetRAX requires a partitioned MSA. SNaQ operates on a set of quartets that can be inferred from gene trees, and a given starting topology (we used the RAxML-NG best ML tree). Both PhyloNet MPL and PhyloNet ML operate on a set of gene trees. We inferred the gene trees via RAxML-NG, one gene tree per MSA partition. Note that SNaQ and PhyloNET account for ILS, while the other tools ignore ILS in their likelihood model.

*Hardware used for the Experiments* We ran experiments A1, A2 and B on a single inhouse cluster compute node with Intel CPUs (E5-2630v3 with 20 MB cache, running at 2.40GHz). The node has 2 CPUs with 8 physical cores each and 64 GB RAM. We used the same cluster for experiment F, using up to 4 compute nodes in the first, and up to 8 compute nodes in the second experiment. We ran experiments C, D, G on a Lenovo Thinkpad T460p laptop. The laptop has 16 GB RAM and contains a single Intel CPU (i7-6700HQ with 6 MB cache, running at 2.60 GHz) with 4 physical cores. We ran experiment E on a machine with an Intel Xeon Gold 6148 (Skylake-SP), 40 cores, 754 GB RAM.

### 3.4 Results and Discussion

We only discuss representative results here, and refer to the supplementary text for comprehensive results (including percentiles and standard deviation) for all experiments. Here, we report the unrooted SCD to assess topological distances to the true simulated network. To quantify the NetRAX search algorithm quality, we compare the BIC scores of the true and the inferred networks. As we optimize for BIC, a worse inferred BIC indicates that the search algorithm got stuck in a local optimum. A better inferred BIC can be encountered because of the finite number of simulated MSA sites (ML is consistent on MSAs with infinite sites).

#### Multiple Starting Trees, 30 taxa, 3 reticulations

In Table 1 and Figure 6, we observe that running NetRAX inferences from multiple starting trees yields more accurate networks than running a NetRAX inference from a single ML tree. This is because a single NetRAX inference run can become stuck in local optima. However, initiating multiple independent NetRAX searches evidently results in a higher accumulated runtime.

**Table 1.**
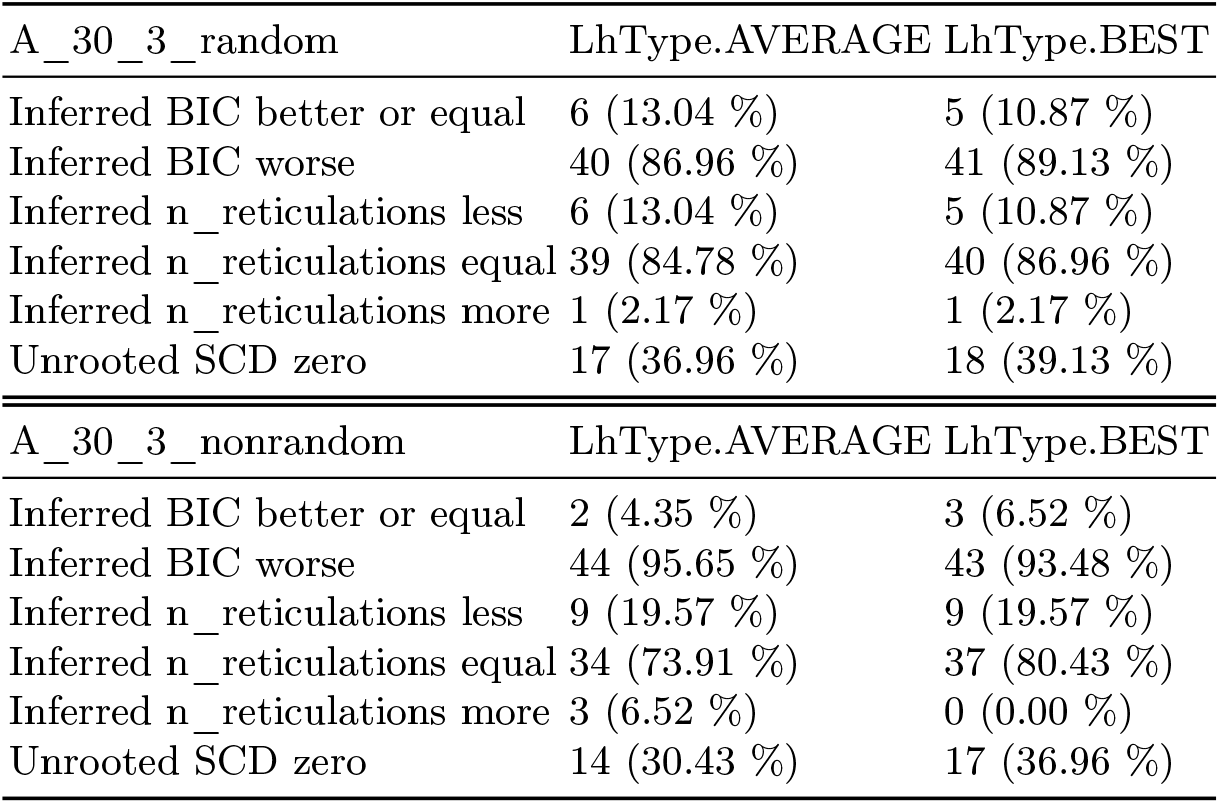
Summary statistics 30 taxa and 3 reticulations. Top: Starting from 3 maximum parsimony and 3 random trees. Bottom: Starting from the RAxML-NG ML tree.

| A_30_3_random | LhType.AVERAGE | LhType.BEST |
| --- | --- | --- |
| Inferred BIC better or equal | 6 (13.04 %) | 5 (10.87 %) |
| Inferred BIC worse | 40 (86.96 %) | 41 (89.13 %) |
| Inferred n_reticulations less | 6 (13.04 %) | 5 (10.87 %) |
| Inferred n_reticulations equal | 39 (84.78 %) | 40 (86.96 %) |
| Inferred n_reticulations more | 1 (2.17 %) | 1 (2.17 %) |
| Unrooted SCD zero | 17 (36.96 %) | 18 (39.13 %) |
| A_30_3_nonrandom | LhType.AVERAGE | LhType.BEST |
| Inferred BIC better or equal | 2 (4.35 %) | 3 (6.52 %) |
| Inferred BIC worse | 44 (95.65 %) | 43 (93.48 %) |
| Inferred n_reticulations less | 9 (19.57 %) | 9 (19.57 %) |
| Inferred n_reticulations equal | 34 (73.91 %) | 37 (80.43 %) |
| Inferred n_reticulations more | 3 (6.52 %) | 0 (0.00 %) |
| Unrooted SCD zero | 14 (30.43 %) | 17 (36.96 %) |

**Fig. 6.**
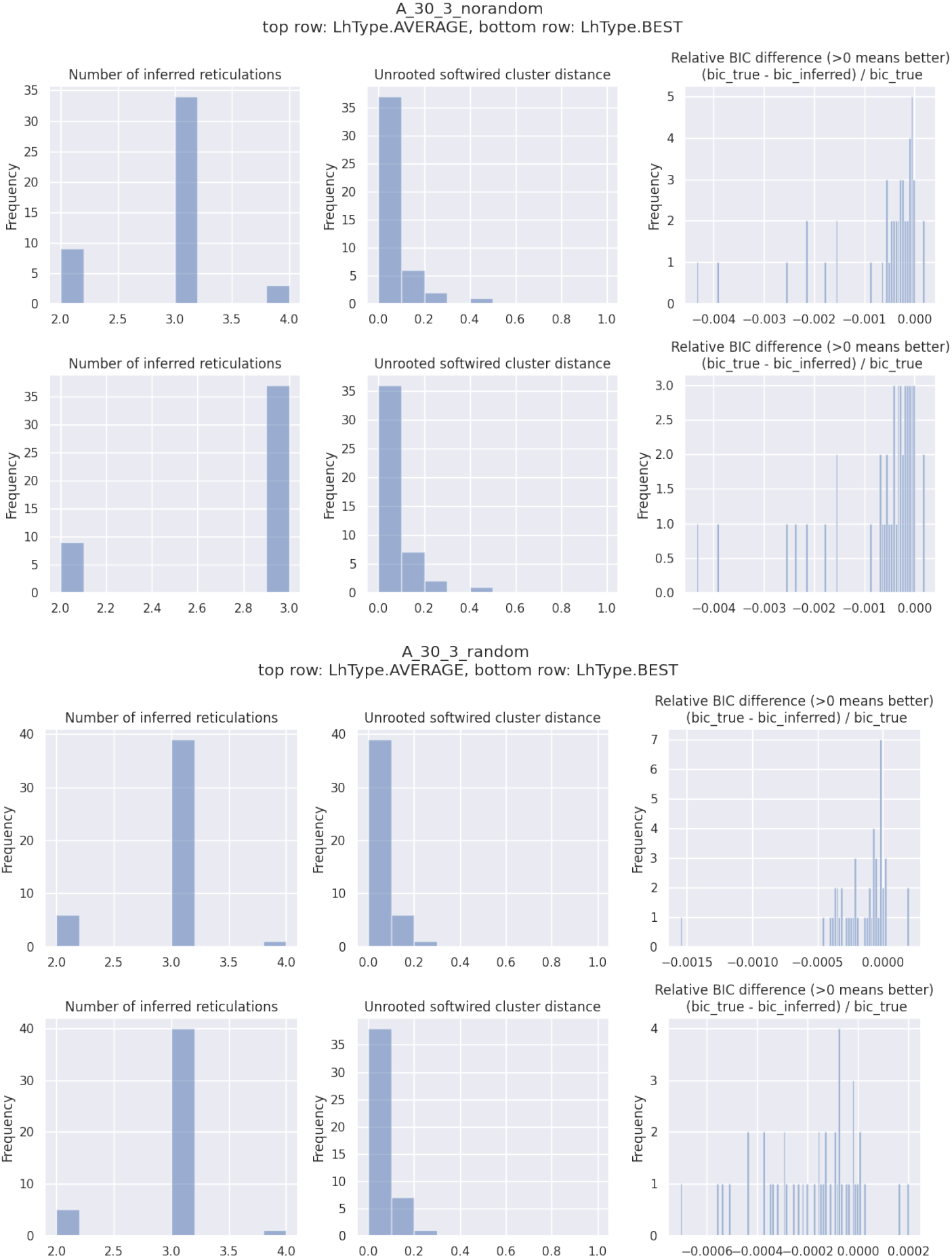
Number of inferred reticulations, unrooted SCD, and relative BIC difference for 50 simulated data sets with 30 taxa and 3 reticulations each. Top: starting from a RAxML-NG ML tree. Bottom: starting 3 random and 3 maximum parsimony trees.

#### Start from ML tree, 40 taxa, 4 reticulations

In Table 2, we see that NetRAX achieves slightly better inference results with `LhType.AVERAGE` than `LhType.BEST`. However, these differences are small, as we can see in Figure 7.

**Table 2.**
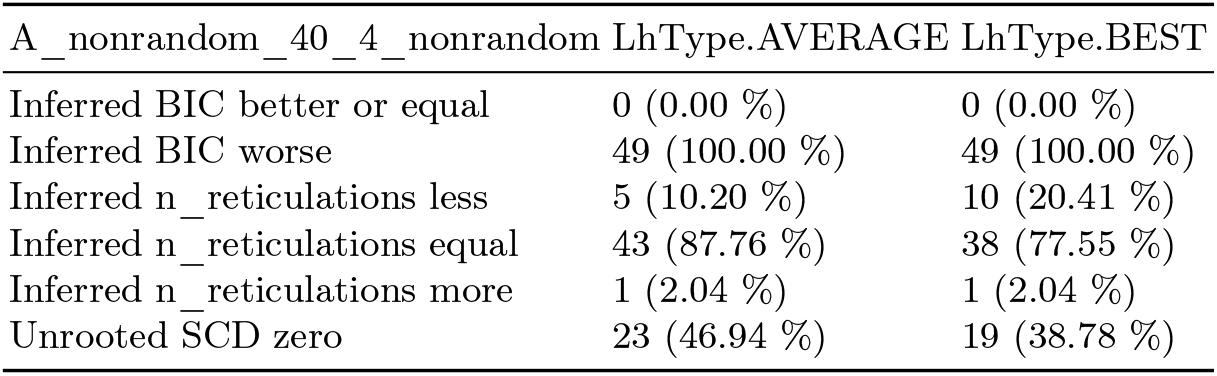
Summary statistics for 40 taxa, 4 reticulations, starting from the RAxML-NG ML tree.

**Fig. 7.**
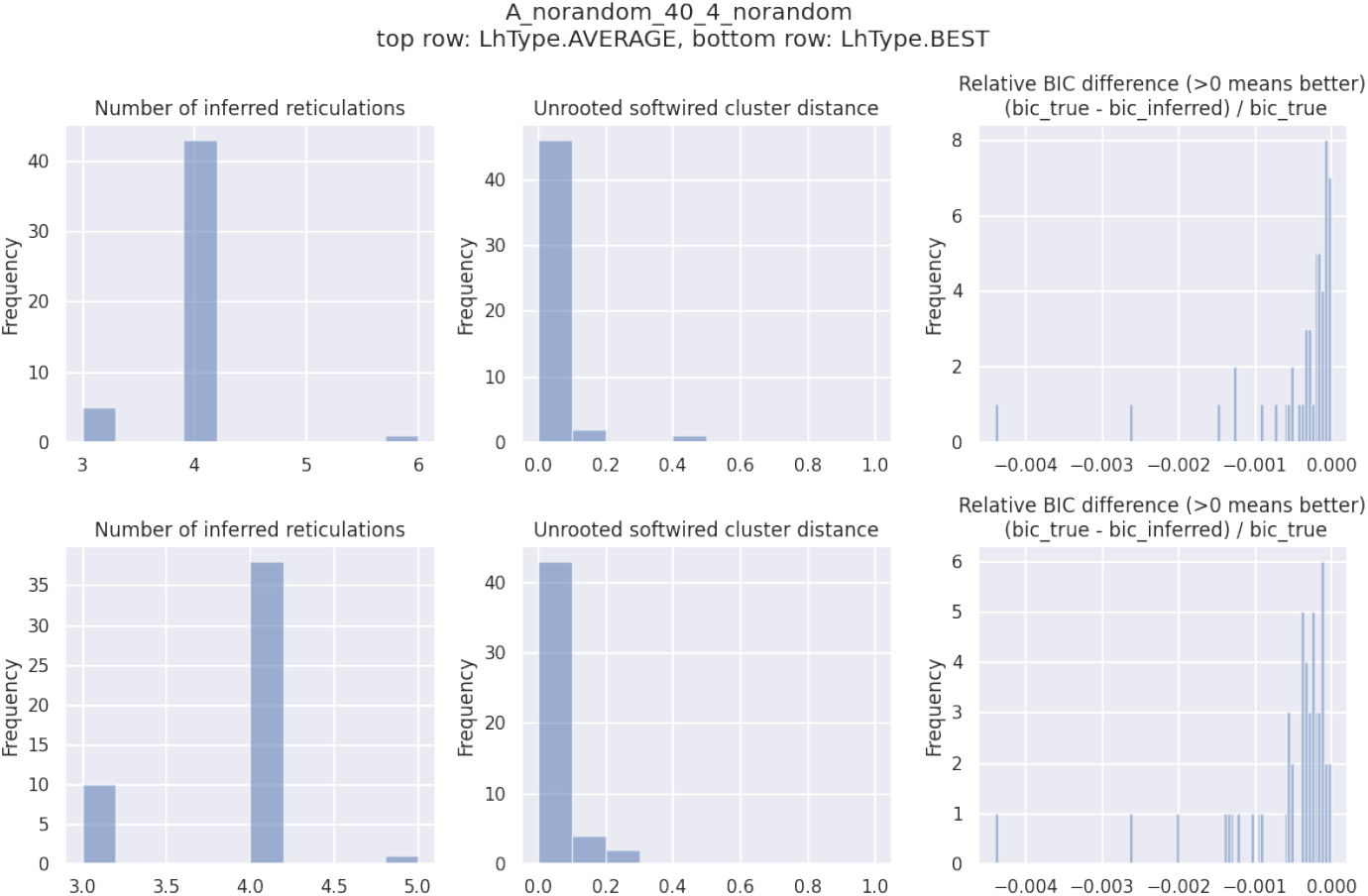
Number of inferred reticulations, unrooted SCD, and relative BIC difference for 50 simulated data sets with 40 taxa and 4 reticulations each, starting from the RAxML-NG ML tree.

#### Reticulation Probability

Under our simulation setting, the number of MSA sites to simulate per displayed tree is proportional to the probability of displaying the tree. Changing the reticulation probability has no adverse effect on NetRAX inference quality, as long as there are sufficient MSA patterns (a *pattern* is a unique site in the MSA) per displayed tree, see Table 3. Note that, with reticulation probability 0.1, we only simulated 200 MSA sites for the less likely displayed tree, with some of the sites consisting of all-equal characters.

**Table 3.**
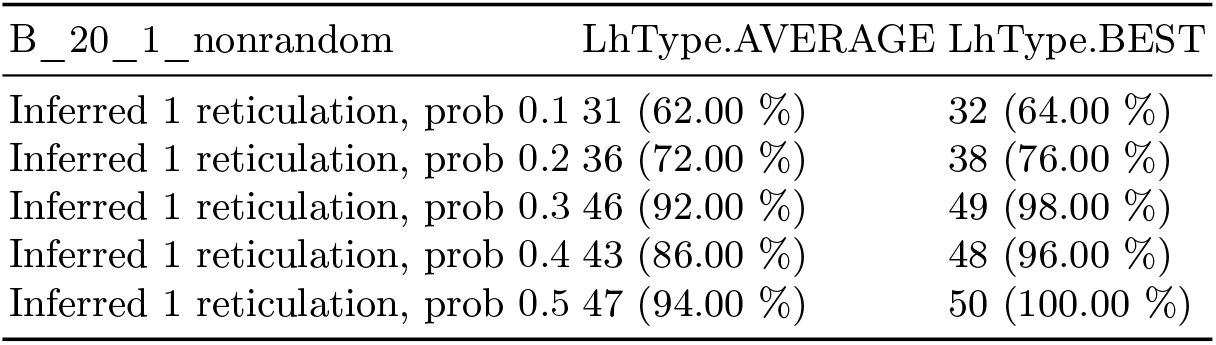
Summary statistics for 50 simulated networks with 20 taxa, 1 reticulation, 2000 simulated MSA sites, starting from the RAxML-NG ML tree, with varying reticulation probability. We report the number of NetRAX inference runs that correctly inferred a single reticulation.

| B_20_1_nonrandom | LhType.AVERAGE | LhType.BEST |
| --- | --- | --- |
| Inferred 1 reticulation, prob 0.1 | 31 (62.00 %) | 32 (64.00 %) |
| Inferred 1 reticulation, prob 0.2 | 36 (72.00 %) | 38 (76.00 %) |
| Inferred 1 reticulation, prob 0.3 | 46 (92.00 %) | 49 (98.00 %) |
| Inferred 1 reticulation, prob 0.4 | 43 (86.00 %) | 48 (96.00 %) |
| Inferred 1 reticulation, prob 0.5 | 47 (94.00 %) | 50 (100.00 %) |

#### Unpartitioned Data

In all NetRAX inference runs with an unpartitioned MSA, NetRAX inferred a bifurcating tree, under both `LhModel.AVERAGE` and `LhModel.BEST`. Thus, NetRAX is not able to infer reticulations on unpartitioned data.

#### Scrambled Partitions

In Table 4, we observe that NetRAX is still able to infer a ‘good’ network if at most 20% of all MSA sites are perturbed among partitions. We observe no substantial difference in result quality between `LhType.BEST` and `LhType.AVERAGE`.

**Table 4.** Results for 30 taxa, 3 reticulations, with scrambled partitions, starting from the RAxML-NG ML tree. Top: `LhType.AVERAGE`, Bottom: `LhType.BEST`. We use **+** for better-or-equal BIC, and - for worse BIC.

| scrambling factor | 0% | 10% | 20% | 30% | 40% | 50% | 60% | 70% | 80% | 90% | 100% |
| --- | --- | --- | --- | --- | --- | --- | --- | --- | --- | --- | --- |
| Inferred BIC | - | - | - | + | + | + | + | + | + | + | + |
| Inferred n_reticulations | **3** | **3** | 5 | **3** | **3** | **3** | **3** | 1 | 0 | 0 | 1 |
| Unrooted SCD | **0** | **0** | 0.09 | 0.17 | 0.21 | 0.21 | 0.21 | 0.35 | 0.38 | 0.38 | 0.36 |
| Runtime RAxML (seconds) | 120 | 125 | 123 | 123 | 127 | 124 | 125 | 133 | 125 | 128 | 126 |
| Runtime NetRAX (seconds) | 380 | 314 | 2274 | 313 | 230 | 314 | 448 | 72 | 10 | 8 | 55 |

| scrambling factor | 0% | 10% | 20% | 30% | 40% | 50% | 60% | 70% | 80% | 90% | 100% |
| --- | --- | --- | --- | --- | --- | --- | --- | --- | --- | --- | --- |
| Inferred BIC | - | - | - | + | + | + | + | + | + | + | + |
| Inferred n_reticulations | **3** | **3** | 5 | **3** | **3** | **3** | 2 | 1 | 0 | 0 | 1 |
| Unrooted SCD | **0** | **0** | 0.09 | 0.17 | 0.21 | 0.23 | 0.21 | 0.36 | 0.38 | 0.38 | 0.37 |
| Runtime RAxML (seconds) | 120 | 125 | 123 | 123 | 127 | 124 | 125 | 133 | 125 | 128 | 126 |
| Runtime NetRAX (seconds) | 277 | 219 | 2062 | 216 | 165 | 156 | 134 | 48 | 6 | 5 | 21 |

#### Varying Alignment Size

In Table 5, we see that NetRAX under `LhType.BEST` is faster for smaller MSAs, but that under `LhType.AVERAGE` it is faster for larger MSAs. The quality of the inferred network is similar among both likelihood types. Based on our empirical observations this surprising speed difference is because (i) a single network lnL computation under `LhType.BEST` is faster than under `LhType.AVERAGE`. (ii) for larger MSAs, we infer the best network with less overall moves, when using `LhType.AVERAGE`. Note that we used data sets with very few, equally-sized partitions here and only one partition per displayed tree.

**Table 5.** Results for 30 taxa, 3 reticulations, different MSA size, starting from the RAxML-NG ML tree. Top: `LhType.AVERAGE`, Bottom: `LhType.BEST`. We use **+** for better-or-equal BIC, and - for worse BIC.

| msa patterns | 397 | 794 | 3882 | 7634 | 36264 | 69919 | 316379 | 597921 |
| --- | --- | --- | --- | --- | --- | --- | --- | --- |
| Inferred BIC | + | - | - | - | - | - | - | - |
| Inferred n_reticulations | 2 | 1 | **3** | **3** | **3** | **3** | **3** | **3** |
| Unrooted SCD | 0.08 | 0.19 | 0.03 | 0 | 0 | 0 | 0.03 | 0.03 |
| Runtime RAxML (seconds) | 5 | 9 | 25 | 34 | 139 | 340 | 2081 | 4340 |
| Runtime NetRAX (seconds) | 101 | 37 | 304 | 379 | 643 | 880 | 4544 | 10452 |

| msa patterns | 397 | 794 | 3882 | 7634 | 36264 | 69919 | 316379 | 597921 |
| --- | --- | --- | --- | --- | --- | --- | --- | --- |
| Inferred BIC | - | - | - | - | - | - | - | - |
| Inferred n_reticulations | 1 | 1 | **3** | **3** | **3** | **3** | **3** | **3** |
| Unrooted SCD | 0.21 | 0.21 | 0.03 | **0** | **0** | **0** | 0.03 | 0.03 |
| Runtime RAxML (seconds) | 5 | 9 | 25 | 34 | 139 | 340 | 2081 | 4340 |
| Runtime NetRAX (seconds) | 14 | 10 | 86 | 151 | 444 | 1165 | 5083 | 11650 |

#### Parallel Scalability

Figure 8 shows the NetRAX inference time using different numbers of MPI processes. The inference under `LhType.BEST` required 17 moves in total: 4 rSPR moves, 6 rNNI moves, 2 arc removal moves, 5 arc insertion moves. The inference under `LhType.AVERAGE` required 12 moves in total: 4 rSPR moves, 3 rNNI moves, 1 arc removal move, 4 arc insertion moves. We recommend using at least 2000 MSA patterns per MPI process.

**Fig. 8.**
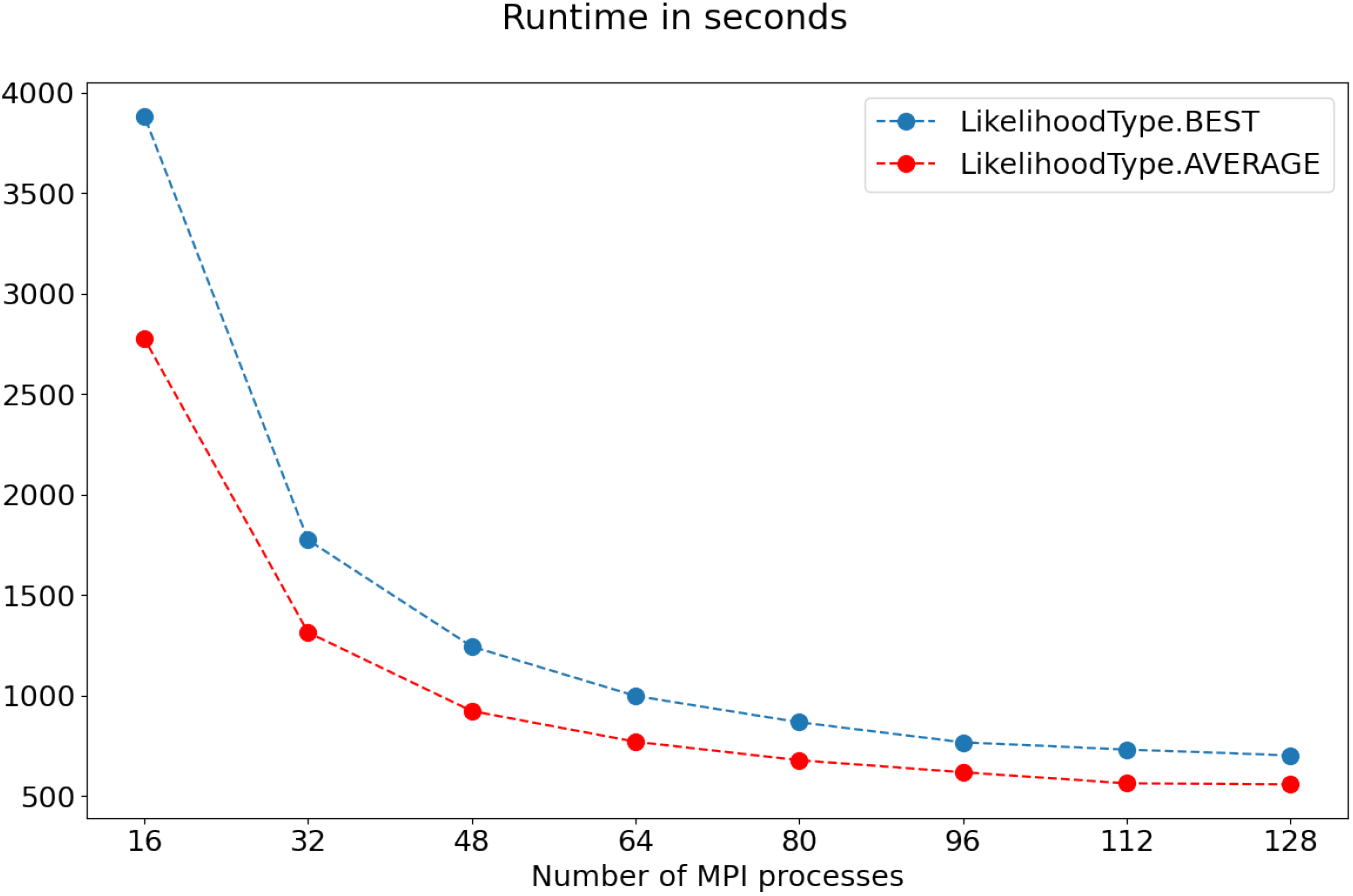
NetRAX inference time in seconds for a single simulated data set with 20 taxa and 3 reticulations, with 189, 212 MSA patterns in total, starting from a RAxML-NG ML tree.

#### Comparison with other tools

For the simulated 10 taxa 1 reticulation data set with 2, 000 MSA sites, we report the total runtime as well as unrooted SCD for all tools in Table 6. Both NetRAX starting from the RAxML-NG ML tree and SNaQ showed perfect inference accuracy, with the unrooted SCD being zero. Only PhyloNET MP was faster than NetRAX, but the tool does not optimize branch lengths. On the simulated 20 taxa 2 reticulations data set, PhyloDAG inferred a network with 14 reticulations. PhyloNET MP with maximum number of reticulations set to 2 only inferred a single reticulation (with unrooted SCD 0.28). NetRAX correctly inferred 2 reticulations under all settings (with a better BIC than the true network but unrooted SCD between 0.19 and 0.22).

**Table 6.** Runtime (in seconds) and unrooted SCD to the true network for inferences using NetRAX, PhyLiNC, PhyloDAG, SNaQ, PhyloNET MPL, and PhyloNet ML. The term `single` refers to starting NetRAX from the best RAxML-NG ML tree. The term `multi` refers to starting NetRAX from 11 unique tree topologies out of 10 random and 10 RAxML-NG maximum parsimony trees. Under all configurations, the network inferred by NetRAX had a better BIC than the true network. ^*a*^: The tool printed error messages, but nonetheless returned a network. ^*b*^: PhyloDAG only returned the network as a picture. We had to manually write the Extended Newick for it. ^*c*^: The inferred network has 2 reticulations, but one reticulation has 0/1 probability. We had to manually prune the network to remove this trivial reticulation.

| Tool | Inference runtime | avg runtime per run | Inferred reticulations | Unrooted SCD |
| --- | --- | --- | --- | --- |
| NetRAX single `LhType.AVERAGE` | 3 | 3 | 1 | 0 |
| NetRAX multi `LhType.AVERAGE` | 40 | 4 | 1 | 0.1 |
| NetRAX single `LhType.BEST` | 2 | 2 | 1 | 0 |
| NetRAX multi `LhType.BEST` | 28 | 3 | 1 | 0.1 |
| PhyLiNC max_reticulations 1 | 45919 | 4592 | 1 | 0.36 |
| PhyLiNC max_reticulations 2<sup>a</sup> | 38365 | 3837 | 2 | 0.56 |
| PhyloDAG<sup>b</sup> | 145 | 145 | 1 | 0.58 |
| SNaQ max_reticulations 1 | 4899 | 490 | 1 | 0 |
| SNaQ max_reticulations 2<sup>c</sup> | 7489 | 749 | 1 | 0 |
| PhyloNET MPL max_reticulations 1 | 158 | 16 | 1 | 0.3 |
| PhyloNET MPL max_reticulations 2 | 223 | 22 | 2 | 0.1 |
| PhyloNET MP max_reticulations 1 | 8 | 2 | 1 | 0 |
| PhyloNET MP max_reticulations 2 | 8 | 2 | 2 | 0.25 |
| PhyloNET ML max_reticulations 1 | 387 | 78 | 1 | 0.27 |
| PhyloNET ML max_reticulations 2 | 19799 | 3960 | 2 | 0.18 |

## 4 Empirical Data

We executed a NetRAX inference on an empirical snake genomes data set [5,3]. We downloaded the individual gene alignments from https://datadryad.org/stash/dataset/doi:10.5061/dryad.4qs50 and merged them into a partitioned MSA, treating each per gene MSA as one partition. The merged data set comprises 23 species, with one individual per species. There are 6, 737 distinct MSA site patterns in the merged MSA, and 304 partitions.

We inferred a ML tree for the complete data set with RAxML-NG using its default GTR+*Γ* substitution model. We then used the ML tree inferred by RAxML-NG as starting network for NetRAX under the `LhType.BEST` and `LhType.AVERAGE` model, with linked branch lengths. We additionally started NetRAX inferences using 14 unique tree topologies, taken from 10 parsimony and 10 random start trees.

We ran the NetRAX inferences on a machine equipped with an Intel Xeon Gold 6148 (Skylake-SP), 40 cores, 754 GB RAM. We compared our NetRAX results with the 1-reticulation network inferred by SNaQ from the Burbrink and Gebara paper.In all cases, NetRAX inferred a better BIC score than the published network (which can be expected, since the reported BIC is based on the network likelihood definition used by NetRAX).

Figure 9 shows the 1-reticulation snakes network inferred by Burbrink and Gehara using the SNaQ tool [3]. NetRAX `LhType.AVERAGE` single, `LhType.AVERAGE` multi, and `LhType.BEST` single inferred the same 1-reticulation network (see Figure 10). NetRAX `LhType.BEST` multi inferred a network with 2 reticulations (see Figure 11). Table 7 presents detailed results on topological distances, inferred BIC scores, and NetRAX inference runtime.

**Fig. 9.**
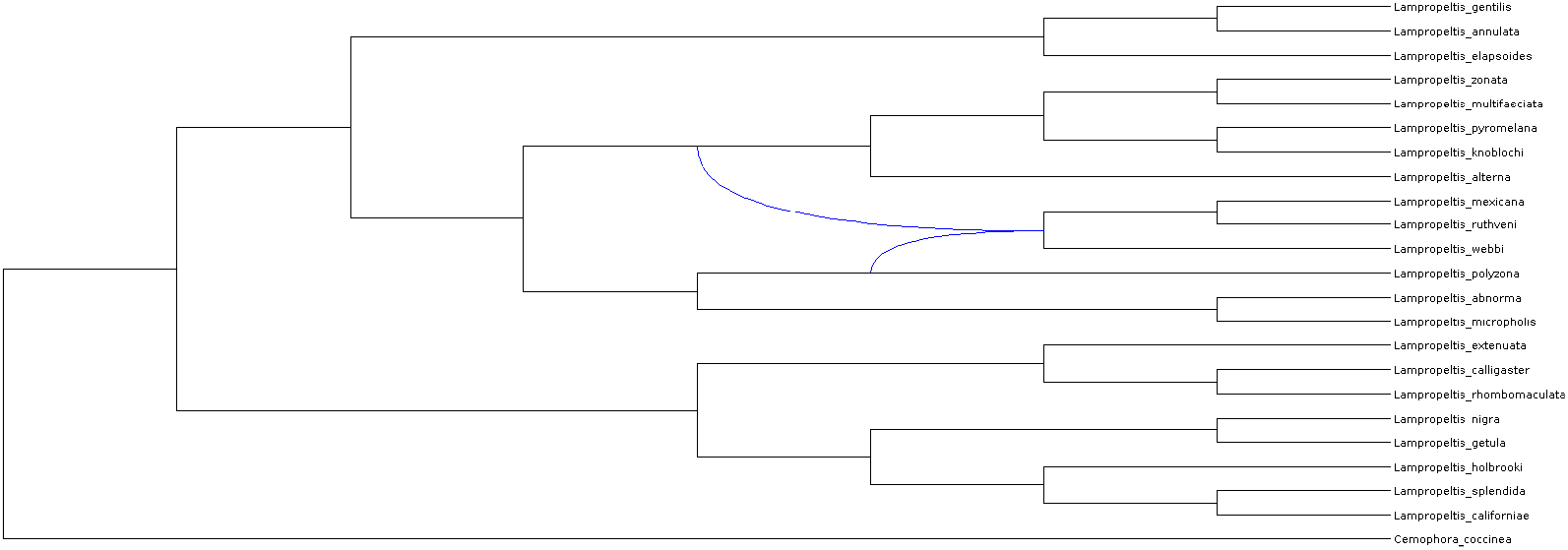
The snakes network as inferred by SNaQ. The network has 1 reticulation. Under `LhModel.BEST`, it has BIC 1499723.786. Under `LhModel.AVERAGE`, it has BIC 1499417.194.

**Fig. 10.**
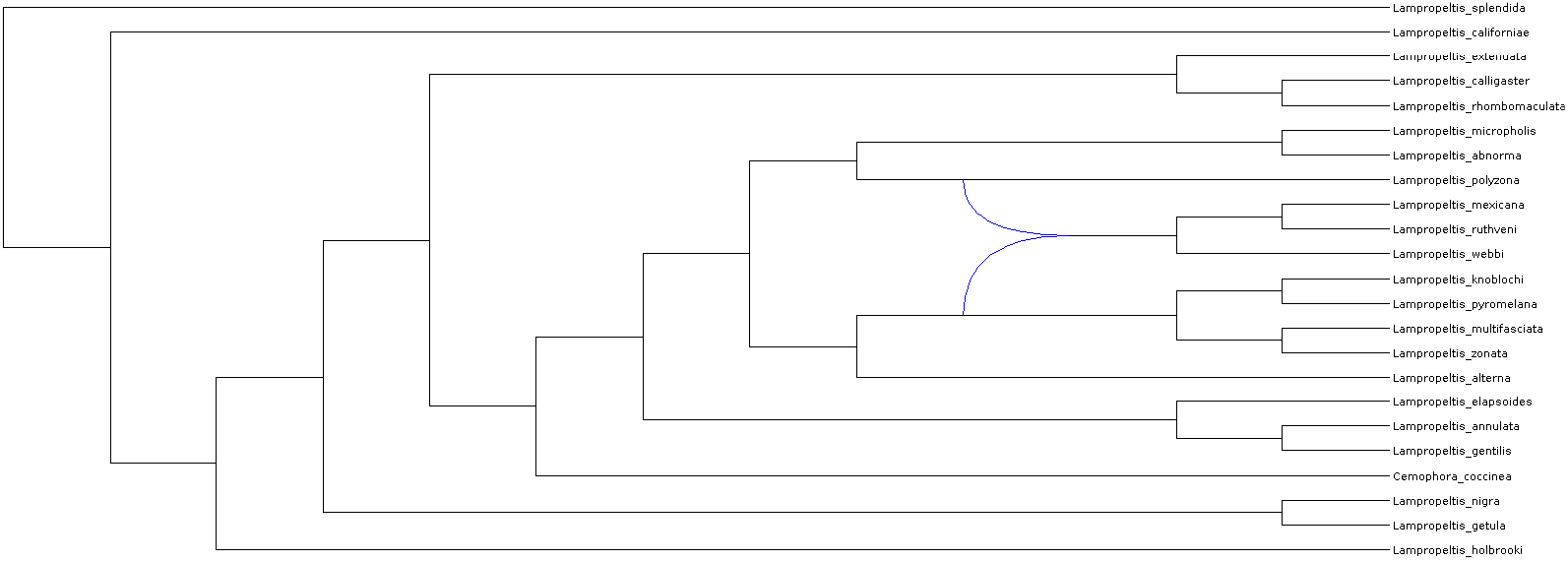
The inferred 1-reticulation network by NetRAX `LhType.AVERAGE` single, `LhType.AVERAGE` multi, and `LhType.BEST` single.

**Fig. 11.**
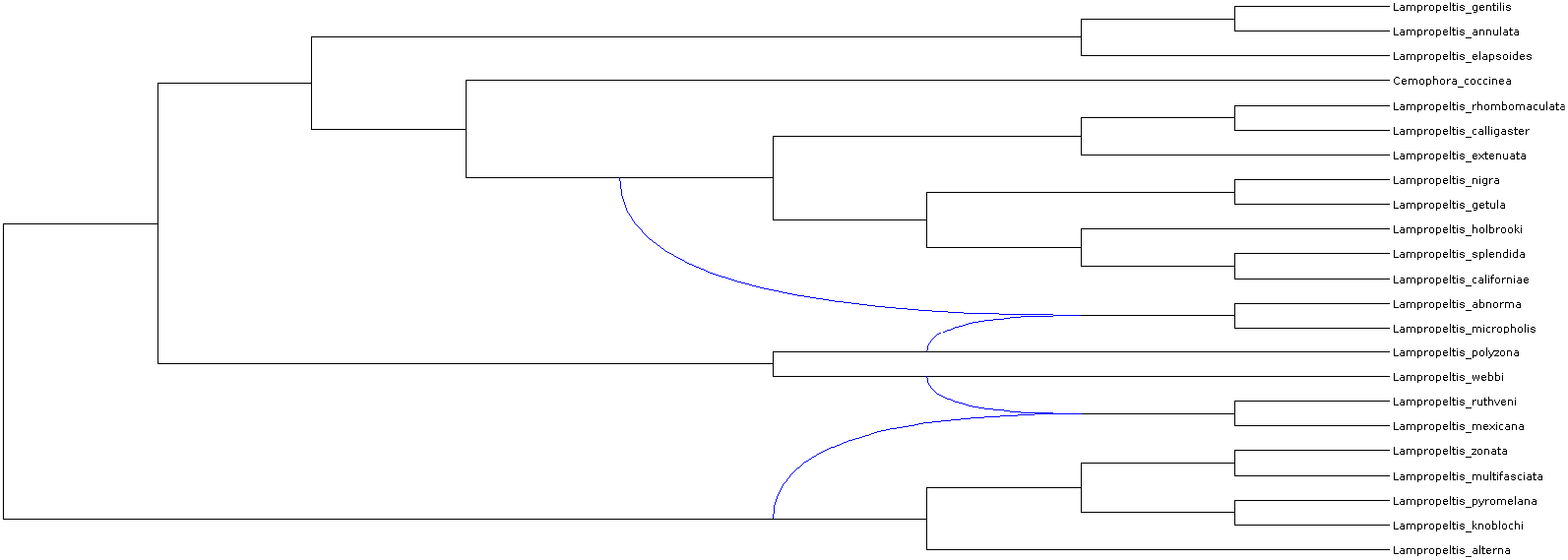
The inferred 2-reticulation network by NetRAX `LhType.BEST` multi.

**Table 7.**
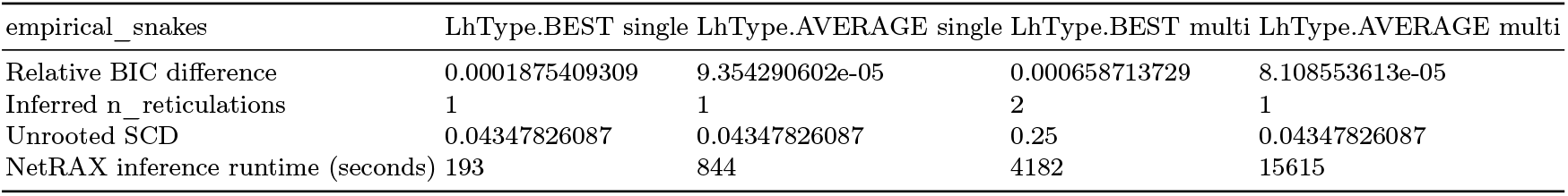
NetRAX results for the empirical snakes data set, for `LhType.BEST` and `LhType.AVERAGE`, compared to the 1-reticulation network inferred by SNaQ. The term `single` refers to starting from the RAxMLNG best ML tree. The term `multi` refers to starting from 14 unique tree topologies out of 10 random and 10 RAxML-NG maximum parsimony trees.

The inferred 1-reticulation network is very similar to the one inferred by SnaQ, and it is in agreement with some of the analyses presented in [3], where they observed “the occasional exclusion of *L. alterna* from the major sister clade”. The inferred 2-reticulation network contains two differences: first, *L. webbi* is not part of the hybrid clade but rather one of its donor; second, a second reticulation is inferred. Note that this network is not a level-1 network, so SnaQ is not able to consider it in its search algorithm. Additional analyses are required to propose a robust phylogeny for this clade since the relative BIC difference under the networks inferred by NetRAX is less than 0.05 percent, and that the current NetRAX search algorithm is prone to getting stuck in local optima. Improving the search algorithm remains future work.

We also initiated a NetRAX inference on an empirical wheat-genomes data set [11], for which Glémin *et al*. estimate that at least 6 reticulations are present. We downloaded the individual gene alignments from https://bioweb.supagro.inra.fr/WheatRelativeHistory/index.php?menu=download and merged them into a partitioned MSA. We treated each gene MSA as one partition.

The merged data set comprises of 47 individuals from 17 species. There are 1, 387, 815 MSA patterns in the merged MSA, and 8738 partitions. We also created a data set with 17 sequences representing the 17 species, by applying a majority-consensus and randomly resolving ties to obtain representative per-species sequences.

We inferred a ML tree for both, the complete, and the subsampled data set with RAxML-NG using its default GTR+GAMMA substitution model. We then used the ML trees inferred by RAxML-NG as starting networks for NetRAX under the `LhType.BEST` model, with linked branch lengths. At the time of the submission and after two months, both inferences are still running. In Figure 12, we present the current best-scoring networks inferred so far using the Dendroscope [15] tool.

**Fig. 12.**
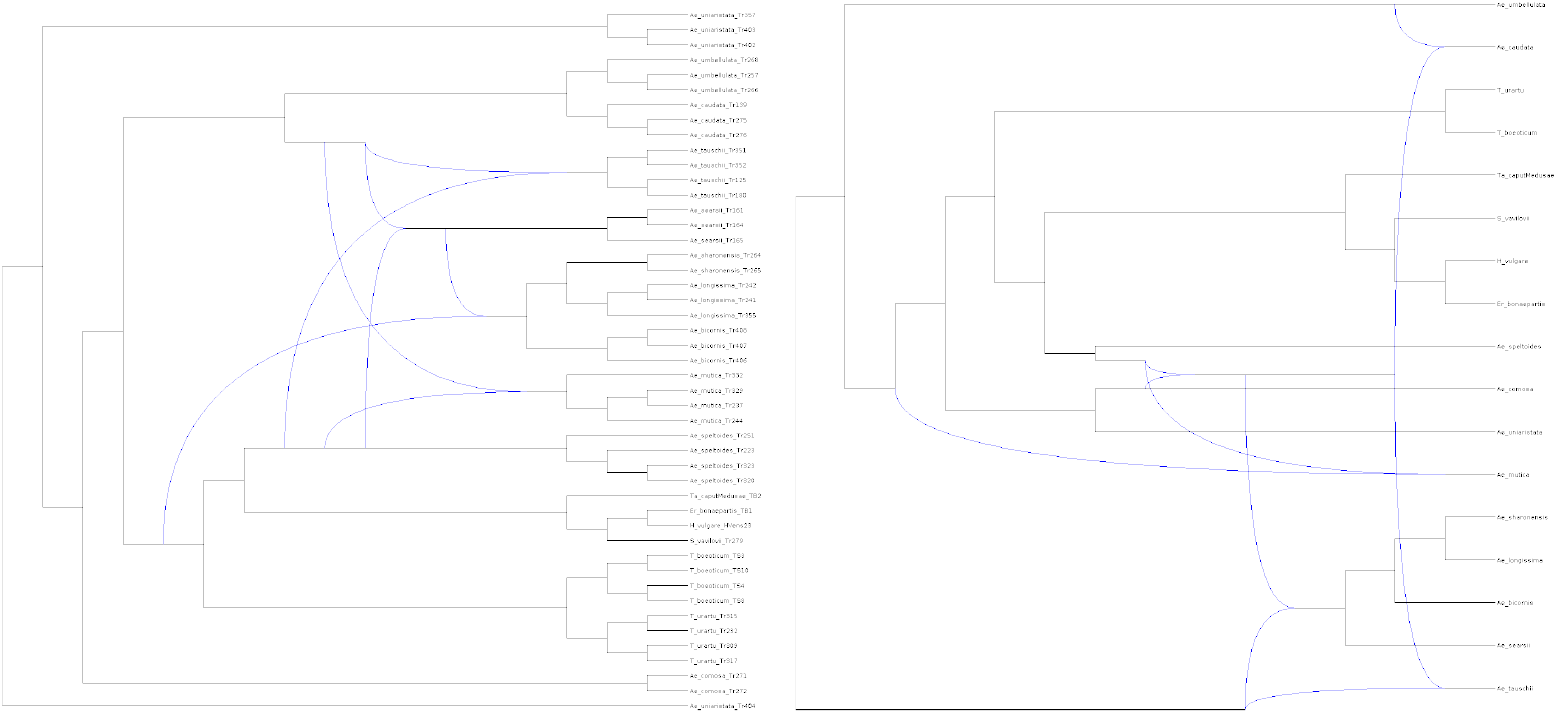
Left: The currently best-scoring network for the full wheat genome data set, with 4 reticulations. Right: The currently best-scoring network for the subsampled wheat genome data set, with 5 reticulations.

It is out of the scope of the paper to present a new evolutionary scenario for this very difficult data set based on ongoing –and thus uncompleted– work. The point we want to make here is that NetRAX scales up and can analyse complex genomic data.

## 5 Conclusion and Future Work

We have presented NetRAX, which, to the best of our knowledge is the only efficient and scalable ML tool for phylogenetic network inference in the absence of ILS that yields accurate resulting networks.

We demonstrated that NetRAX can infer ML networks with up to 40 taxa and up to 4 reticulations on data sets with thousands of MSA sites in less than a day. Our experimental results on simulated data sets show that NetRAX infers high-quality ML networks with very low unrooted SCDs and very low relative BIC differences compared to the true, simulated network. We also demonstrate that NetRAX yields biologically plausible results on a well-studied empirical dataset.

Starting the network inference from multiple starting trees tends to yield more accurate results. For large data sets, we nonetheless recommend using NetRAX with a single ML tree inferred, for instance, via RAxML-NG to keep inference times within acceptable limits. Our experiments show that NetRAX can infer highly accurate ML networks, even with a single ML starting tree.

Future work includes further improving the scalability of NetRAX on larger ML networks and MSAs, and extending its model. For example, we intend to develop a scaled branches model, where all partitions share the same set of branch lengths, but branch lengths are scaled proportionally per partition via a single per-partition scaling parameter.

## References

1. Allen-Savietta, C.: Estimating Phylogenetic Networks from Concatenated Sequence Alignments. The University of Wisconsin-Madison (2020)

2. Ané, C.: Phylonetworks users google group discussion. https://groups.google.com/g/phylonetworks-users/c/KCu45cDRy_Q/m/RLpaZJajBAAJ (2021), website. Accessed August 14th, 2021

3. Burbrink, F.T., Gehara, M.: The biogeography of deep time phylogenetic reticulation. Systematic biology 67(5), 743–755 (2018)

4. Cao, Z., Liu, X., Ogilvie, H.A., Yan, Z., Nakhleh, L.: Practical aspects of phylogenetic network analysis using phylonet. BioRxiv p. 746362 (2019)

5. Chen, X., Lemmon, A.R., Lemmon, E.M., Pyron, R.A., Burbrink, F.T.: Using phylogenomics to understand the link between biogeographic origins and regional diversification in ratsnakes. Molecular phylogenetics and evolution 111, 206–218 (2017)

6. Darriba, D.: pll-modules. https://github.com/ddarriba/pll-modules (2016), website. Accessed July 28, 2021

7. Felsenstein, J.: Evolutionary trees from dna sequences: a maximum likelihood approach. Journal of molecular evolution 17(6), 368–376 (1981)

8. Flouri, T.: Computing the likelihood of a tree. https://github.com/xflouris/libpll/wiki/Computing-the-likelihood-of-a-tree (2015), xwebsite. Accessed July 28, 2021

9. Flouri, T.: libpll-2. https://github.com/xflouris/libpll-2.git (2015), website. Accessed July 28, 2021

10. Gambette, P., Van Iersel, L., Jones, M., Lafond, M., Pardi, F., Scornavacca, C.: Rearrangement moves on rooted phylogenetic networks. PLoS computational biology 13(8), e1005611 (2017)

11. Glémin, S., Scornavacca, C., Dainat, J., Burgarella, C., Viader, V., Ardisson, M., Sarah, G., Santoni, S., David, J., Ranwez, V.: Pervasive hybridizations in the history of wheat relatives. Science advances 5(5), eaav9188 (2019)

12. Hejase, H.A., Liu, K.J.: A scalability study of phylogenetic network inference methods using empirical datasets and simulations involving a single reticulation. BMC bioinformatics 17(1), 1–12 (2016)

13. Holoborodko, P.: Mpfr c++. http://www.holoborodko.com/pavel/mpfr/ (2010), website. Accessed July 28, 2021

14. Huson, D.H., Rupp, R., Scornavacca, C.: Phylogenetic networks: concepts, algorithms and applications. Cambridge University Press (2010)

15. Huson, D.H., Scornavacca, C.: Dendroscope 3: an interactive tool for rooted phylogenetic trees and networks. Systematic biology 61(6), 1061–1067 (2012)

16. Jin, G., Nakhleh, L., Snir, S., Tuller, T.: Maximum likelihood of phylogenetic networks. Bioinformatics 22(21), 2604–2611 (2006)

17. Kozlov, A.M., Darriba, D., Flouri, T., Morel, B., Stamatakis, A.: Raxml-ng: a fast, scalable and user-friendly tool for maximum likelihood phylogenetic inference. Bioinformatics 35(21), 4453–4455 (2019)

18. Nakhleh, L., Jin, G., Zhao, F., Mellor-Crummey, J.: Reconstructing phylogenetic networks using maximum parsimony. In: 2005 IEEE Computational Systems Bioinformatics Conference (CSB’05). pp. 93–102. IEEE (2005)

19. NEPAL: http://old-bioinfo.cs.rice.edu/nepal/ (2006), website. Accessed July 28, 2021

20. Nguyen, Q., Roos, T.: Likelihood-based inference of phylogenetic networks from sequence data by phylodag. In: International Conference on Algorithms for Computational Biology. pp. 126–140. Springer (2015)

21. Pardi, F., Scornavacca, C.: Reconstructible phylogenetic networks: do not distinguish the indistinguishable. PLoS computational biology 11(4), e1004135 (2015)

22. Park, H.J., Nakhleh, L.: Inference of reticulate evolutionary histories by maximum likelihood: the performance of information criteria. In: BMC bioinformatics. vol. 13, pp. 1–10. BioMed Central (2012)

23. Rambaut, A., Grass, N.C.: Seq-gen: an application for the monte carlo simulation of dna sequence evolution along phylogenetic trees. Bioinformatics 13(3), 235–238 (1997)

24. Robinson, D.F., Foulds, L.R.: Comparison of phylogenetic trees. Mathematical biosciences 53(1-2), 131–147 (1981)

25. Solís-Lemus, C., Ané, C.: Inferring phylogenetic networks with maximum pseudolikelihood under incomplete lineage sorting. PLoS genetics 12(3), e1005896 (2016)

26. Solís-Lemus, C., Bastide, P., Ané, C.: Phylonetworks: a package for phylogenetic networks. Molecular biology and evolution 34(12), 3292–3298 (2017)

27. Strimmer, K., Moulton, V.: Likelihood analysis of phylogenetic networks using directed graphical models. Molecular biology and evolution 17(6), 875–881 (2000)

28. Tavaré, S., et al.: Some probabilistic and statistical problems in the analysis of dna sequences. Lectures on mathematics in the life sciences 17(2), 57–86 (1986)

29. Wen, D., Yu, Y., Zhu, J., Nakhleh, L.: Inferring phylogenetic networks using phylonet. Systematic biology 67(4), 735–740 (2018)

30. Zhang, C., Ogilvie, H.A., Drummond, A.J., Stadler, T.: Bayesian inference of species networks from multilocus sequence data. Molecular biology and evolution 35(2), 504–517 (2018)

