## Supplementary Text for "NetRAX: Accurate and Fast Maximum Likelihood Phylogenetic Network Inference^⋆^"

**Abstract.** In this supplementary text, we derive the first and second derivatives for the phylogenetic network likelihood function, explain technical details on how to make libpll/pll-modules work with networks, define normalized variants of common topological distance measures on networks, and provide detailed experimental results on simulated and empirical data.

### 1 Phylogenetic Network Loglikelihood Derivatives

#### 1.1 Derivatives of Tree loglikelihood and Tree Likelihood

The derivatives of the tree loglikelihood for tree $T$ and alignment partition $A_i$ are:

$$(\ln L(T|A_i,\vartheta_i))' = \sum_{s\in A_i} \ln L(T|s,\vartheta_i)' \quad = \sum_{s\in A_i} \frac{L'(T|s,\vartheta_i)}{L(T|s,\vartheta_i)}. \tag{1}$$

$$\begin{aligned}(\ln L(T|A_i,\vartheta_i))'' &= \left(\sum_{s\in A_i} \frac{L'(T|s,\vartheta_i)}{L(T|s,\vartheta_i)}\right)' \\ &= \sum_{s\in A_i} \left(\frac{L'(T|s,\vartheta_i)}{L(T|s,\vartheta_i)}\right)' \\ &= \sum_{s\in A_i} \frac{L''(T|s,\vartheta_i) * L(T|s,\vartheta_i) - (L'(T|s,\vartheta_i))^2}{(L(T|s,\vartheta_i))^2}.\end{aligned} \tag{2}$$

We compute the tree partition likelihood derivatives out of $L(T|A_i,\vartheta_i)$, $(\ln L(T|A_i,\vartheta_i))'$ and $(\ln L(T|A_i,\vartheta_i))''$ as follows:

$$\begin{aligned}L'(T|A_i,\vartheta_i) &= \left(\prod_{s\in A_i} L(T|s,\vartheta_i)\right)' \\ &= \left(\prod_{s\in A_i} L(T|s,\vartheta_i)\right) * \left(\sum_{s\in A_i} \frac{L'(T|s,\vartheta_i)}{L(T|s,\vartheta_i)}\right) \\ &= L(T|A_i,\vartheta_i) * (\ln L(T|A_i,\vartheta_i))'.\end{aligned} \tag{3}$$

$$\begin{aligned}L''(T|A_i,\vartheta_i) &= \big(L(T|A_i,\vartheta_i) * (\ln L(T|A_i,\vartheta_i))'\big)' \\ &= L'(T|A_i,\vartheta_i) * (\ln L(T|A_i,\vartheta_i))' + L(T|A_i,\vartheta_i) * (\ln L(T|A_i,\vartheta_i))''.\end{aligned} \tag{4}$$

##### 1.2 Network Loglikelihood Derivatives

The first and second derivatives of the phylogenetic network loglikelihood (with respect to a changed branch length) are:

$$(\ln L(N|\mathcal{A},\vartheta))' = \sum_{i=1}^{p} (\ln L(N|A_i,\vartheta_i))'. \tag{5}$$

$$(\ln L(N|\mathcal{A},\vartheta))'' = \sum_{i=1}^{p} (\ln L(N|A_i,\vartheta_i))''. \tag{6}$$

**Weighted Average Version**

$$\begin{aligned}(\ln L(N|A_i,\vartheta_i))' &= \left(\ln\left(\sum_{T\in\mathcal{T}(N)} L(T|A_i,\vartheta_i) * P(T|N)\right)\right)' \\ &= \frac{\left(\sum_{T\in\mathcal{T}(N)} L(T|A_i,\vartheta_i) * P(T|N)\right)'}{\sum_{T\in\mathcal{T}(N)} L(T|A_i,\vartheta_i) * P(T|N)} \\ &= \frac{\sum_{T\in\mathcal{T}(N)} L'(T|A_i,\vartheta_i) * P(T|N)}{\sum_{T\in\mathcal{T}(N)} L(T|A_i,\vartheta_i) * P(T|N)}.\end{aligned} \tag{7}$$

Let $u := \sum_{T\in\mathcal{T}(N)} L'(T|A_i,\vartheta_i) * P(T|N)$ and $v := \left(\sum_{T\in\mathcal{T}(N)} L(T|A_i,\vartheta_i) * P(T|N)\right)$.

$$\begin{aligned}u' &= \left(\sum_{T\in\mathcal{T}(N)} L'(T|A_i,\vartheta_i) * P(T|N)\right)' \\ &= \sum_{T\in\mathcal{T}(N)} L''(T|A_i,\vartheta_i) * P(T|N).\end{aligned} \tag{8}$$

$$\begin{aligned}v' &= \left(\sum_{T\in\mathcal{T}(N)} L(T|A_i,\vartheta_i) * P(T|N)\right)' \\ &= \sum_{T\in\mathcal{T}(N)} L'(T|A_i,\vartheta_i) * P(T|N).\end{aligned} \tag{9}$$

Using equations 8 and 9, we obtain:

$$\begin{aligned}(\ln L(N|A_i,\vartheta_i))'' &= \left(\frac{\sum_{T\in\mathcal{T}(N)} L'(T|A_i,\vartheta_i) * P(T|N)}{\sum_{T\in\mathcal{T}(N)} L(T|A_i,\vartheta_i) * P(T|N)}\right)' \\ &= \left(\frac{u}{v}\right)' \\ &= \frac{u' * v - u * v'}{v^2} \\ &= \frac{\left(\sum_{T\in\mathcal{T}(N)} L''(T|A_i,\vartheta_i) * P(T|N)\right) * \left(\sum_{T\in\mathcal{T}(N)} L(T|A_i,\vartheta_i) * P(T|N)\right) - \left(\sum_{T\in\mathcal{T}(N)} L'(T|A_i,\vartheta_i) * P(T|N)\right)^2}{\left(\sum_{T\in\mathcal{T}(N)} L(T|A_i,\vartheta_i) * P(T|N)\right)^2}.\end{aligned} \tag{10}$$

**Best Tree Version** We know that

$$(\max f(x), g(x))' = \begin{cases} f'(x), & \text{if } f(x) > g(x) \\ g'(x), & \text{if } f(x) < g(x) \\ f'(x), & \text{if } f(x) = g(x) \text{ and } f'(x) = g'(x) \\ \text{undefined}, & \text{otherwise.} \end{cases} \quad (11)$$

We assume that for empirical use it holds that

$$(L(T_1|A_i, \vartheta_i) * P(T_1|N) = L(T_2|A_i, \vartheta_i) * P(T_2|N)) \iff T_1 = T_2. \quad (12)$$

Let $\hat{T}$ be a displayed tree for which $L(T|A_i, \vartheta_i) * P(T|N)$ is maximal among all $T \in \mathcal{T}(N)$. With the above assumption, we obtain

$$\begin{aligned} (\ln L(N|A_i, \vartheta_i))' &= \left(\ln\left(\max_{T\in\mathcal{T}(N)} L(T|A_i, \vartheta_i) * P(T|N)\right)\right)' \\ &= \frac{\left(\max_{T\in\mathcal{T}(N)} L(T|A_i, \vartheta_i) * P(T|N)\right)'}{\max_{T\in\mathcal{T}(N)} L(T|A_i, \vartheta_i) * P(T|N)} \\ &= \frac{L'(\hat{T}|A_i, \vartheta_i) * P(\hat{T}|N)}{L(\hat{T}|A_i, \vartheta_i) * P(\hat{T}|N)} = \frac{L'(\hat{T}|A_i, \vartheta_i)}{L(\hat{T}|A_i, \vartheta_i)} \\ &= \left(\ln L(\hat{T}|A_i, \vartheta_i)\right)'. \end{aligned} \quad (13)$$

$$(\ln L(N|A_i, \vartheta_i))'' = \left(\ln L(\hat{T}|A_i, \vartheta_i)\right)''. \quad (14)$$

#### 2 Wrapping and Adapting libpll/pll-modules/RAxML-NG to work with Networks

A detailed explanation of CLVs and tree likelihood computation is available here: https://github.com/xflouris/libpll/wiki/Computing-the-likelihood-of-a-tree

Since libpll operates on a strictly bifurcating phylogenetic tree, it requires each inner node to have exactly two children. For each reticulation parent taken in order to display a tree in a network, we mark the reticulation edge coming from the alternative reticulation parent as inactive. This temporarily removes a child from the non-chosen reticulation parent.

We apply some adaptations to deploy libpll on networks: We need to handle single-child nodes (see Section 2.2), and we need to skip dead (unused) nodes (see Section 2.3).

##### 2.1 Fake Treeinfo

All operations in libpll, pll-modules, and RAxML-NG require phylogenetic trees. In order to make some of them work on networks as well, we implemented substantial changes in forked versions of their repositories. Most functions provided by pll-modules require a huge pllmod_treeinfo_t data structure. This structure stores, among other useful piece of information (such as pll_partition_t information), a pll_utree_t tree. For our use with networks, we created a fake treeinfo data structure with an empty tree, but overrode some of its parameters (like number of nodes in the "tree"). We also added function pointers to both pll_treeinfo_t and the TreeInfo class from RAxML-NG. We need these function pointers to call our own network likelihood function within the branch-length and model optimization procedures.

##### 2.2 Handling Single-child Nodes

Setting the reticulation node parents (activating one parent, deactivating the other parent) in order to display a particular tree results in a tree-like structure which is not strictly bifurcating. After choosing the active reticulation parents, each reticulation only has one active parent and one active child. However, libpll requires a strictly bifurcating tree for its likelihood computations. In order to handle these paths and to avoid complicated code that temporarily merges nodes and branches, we use the following trick: we introduce a new, "fake" node. We set all conditional per-site likelihoods in the CLV for the "fake" node to 1.0. We can interpret the "fake" node as an extra taxon that has only gap characters in its MSA sequence. Instead of skipping or merging a single-child node during likelihood computation, we temporarily add the "fake" node as its second child, with branch length zero.

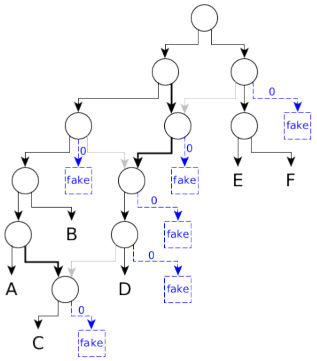


**Fig. 1.** A displayed tree which contains nodes that have only one active child. For likelihood computation using libpll, we add a temporary fake node with branch length zero as the second active child of all single-child nodes.

##### 2.3 Skipping Dead Nodes

Depending on which reticulation parents we select, it can happen that a large part of the resulting displayed tree consists of dead nodes (see Figure 2). A *dead* node is a non-leaf node that either has no active children, or is the only active child of a dead node, or a node with no active parent and only a single active child. We pre-detect such nodes in order to reduce the total number of required CLV update operations. We also handle the case where we need to assign the root node for a given displayed tree to another node, like in the situation presented in Figure 2.

#### 3 A note on Branch-Length Optimization

When optimizing a branch length, we do not need to recompute the per-partition lnL and its derivatives for trees where the branch is inactive. A branch is *active* in a displayed tree, if neither its source nor its target is a dead node, and it is not connecting an inactive parent to a reticulation (see Figure 3).

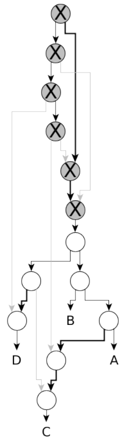


**Fig. 2.** Dead and therefore skipped nodes in a given displayed tree. In this figure, we mark dead nodes by the letter X. Note that the original root node belongs to the set of dead nodes. For this displayed tree, we use the highest non-dead node as its temporary root node.

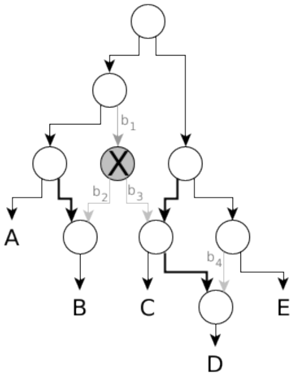


**Fig. 3.** A displayed tree in a phylogenetic network. The branches $b_1$, $b_2$, $b_3$, and $b_4$ are inactive in this displayed tree. When optimizing the length of any of the inactive branches, we do not need to recompute the likelihood of this displayed tree.

#### 4 More Distance Measures

We define normalized versions of further topological network distance measures from Huson and Scornavacca [1]. We call a property (cluster, tripartition, nested label, ...) of a network *trivial* if it occurs in all networks defined on the same set of taxa. We diverge from the distance definitions from Huson and Scornavacca by explicitly discarding the trivial properties of a network:

*(Normalized) X Distance* For a given network $N$ and a property $X$, let $X(N)$ be the set of nontrivial $X$ of $N$. The $X$ distance between two networks $N_1$ and $N_2$ is:

$$\frac{|X(N_1) \triangle X(N_2)|}{|X(N_1) \cup X(N_2)|}.$$

#### 5 The Elbow Method

Figure!4 gives an example of the Elbow method.

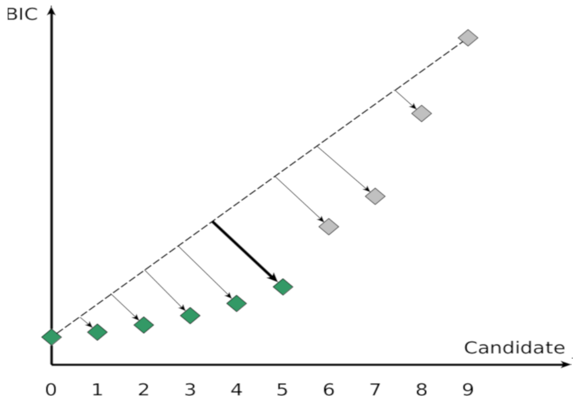


**Fig. 4.** In the Elbow Method [3,2], we sort the list of pre-scored move candidates by increasing BIC score. We need to find the point with the largest distance to the line from the first to the last candidate; this point corresponds to our chosen cutoff value. We keep all candidates with BIC score smaller-or-equal than this cutoff score value. In this example, we keep candidates 0 to 5.

#### 6 Detailed Experimental Results on Simulated Data

In this section, we provide detailed plots and tables for our experiments simulated data. We provide plots on overall simulated dataset statistics (number of taxa, number of reticulations, number of MSA patterns, number of near-zero branches), list percentiles for various topological distance measures and lnL/AIC/BIC/AICc scores, and plot distributions on normalized network distances. We also judge the overall inference quality using:

- Good result: BIC better-or-equal or unrooted softwired distance zero
- Passable result: BIC worse, unrooted softwired distance $>0$, but correct number of reticulations
- Bad result: BIC worse, unrooted softwired distance $>0$, wrong number of reticulations

##### 6.1 A1: Multiple starting trees, 10 taxa, 1 reticulation

In this experiment, we simulated 50 networks each for 10 taxa and 1 reticulation. In addition to the NetRAX inference starting from the RAxML-NG best ML tree, we also started another NetRAX inference using 3 random and 3 maximum parsimony starting trees.

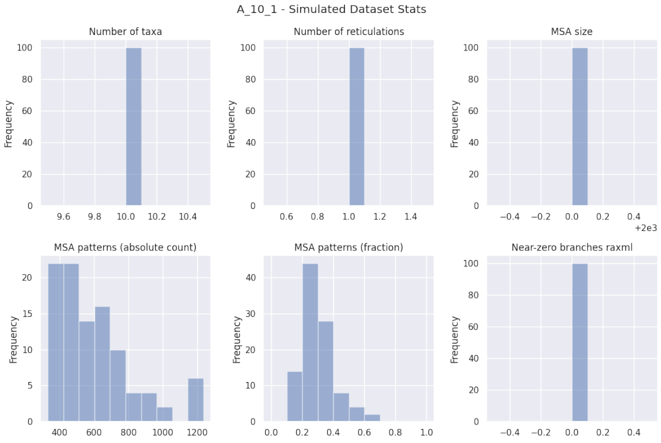


**Fig. 5.** Simulated dataset statistics for experiment A1, 10 taxa, 1 reticulation.

| A_10_1_norandom | LhType.AVERAGE | LhType.BEST |
|---|---|---|
| Inferred BIC better or equal | 24 (48.00 %) | 25 (50.00 %) |
| Inferred AIC better or equal | 20 (40.00 %) | 21 (42.00 %) |
| Inferred AICc better or equal | 20 (40.00 %) | 21 (42.00 %) |
| Inferred BIC worse | 26 (52.00 %) | 25 (50.00 %) |
| Inferred AIC worse | 30 (60.00 %) | 29 (58.00 %) |
| Inferred AICc worse | 30 (60.00 %) | 29 (58.00 %) |
| Inferred lnL better or equal | 19 (38.00 %) | 20 (40.00 %) |
| Inferred lnL worse | 31 (62.00 %) | 30 (60.00 %) |
| Inferred n_reticulations less | 9 (18.00 %) | 12 (24.00 %) |
| Inferred n_reticulations equal | 41 (82.00 %) | 38 (76.00 %) |
| Inferred n_reticulations more | 0 (0.00 %) | 0 (0.00 %) |
| Unrooted softwired distance zero | 21 (42.00 %) | 18 (36.00 %) |
| Good result | 35 (70.00 %) | 32 (64.00 %) |
| Passable result | 12 (24.00 %) | 12 (24.00 %) |
| Bad result | 3 (6.00 %) | 6 (12.00 %) |

| A_10_1_random | LhType.AVERAGE | LhType.BEST |
|---|---|---|
| Inferred BIC better or equal | 27 (54.00 %) | 28 (56.00 %) |
| Inferred AIC better or equal | 23 (46.00 %) | 24 (48.00 %) |
| Inferred AICc better or equal | 23 (46.00 %) | 24 (48.00 %) |
| Inferred BIC worse | 23 (46.00 %) | 22 (44.00 %) |
| Inferred AIC worse | 27 (54.00 %) | 26 (52.00 %) |
| Inferred AICc worse | 27 (54.00 %) | 26 (52.00 %) |
| Inferred lnL better or equal | 22 (44.00 %) | 23 (46.00 %) |
| Inferred lnL worse | 28 (56.00 %) | 27 (54.00 %) |
| Inferred n_reticulations less | 7 (14.00 %) | 7 (14.00 %) |
| Inferred n_reticulations equal | 43 (86.00 %) | 43 (86.00 %) |
| Inferred n_reticulations more | 0 (0.00 %) | 0 (0.00 %) |
| Unrooted softwired distance zero | 28 (56.00 %) | 27 (54.00 %) |
| Good result | 42 (84.00 %) | 43 (86.00 %) |
| Passable result | 7 (14.00 %) | 6 (12.00 %) |
| Bad result | 1 (2.00 %) | 1 (2.00 %) |

**Table 1.** Summary statistics for experiment A1, 10 taxa, 1 reticulation. Left: starting from RAxML-NG best tree, right: starting from 3 maximum parsimony and 3 random trees.

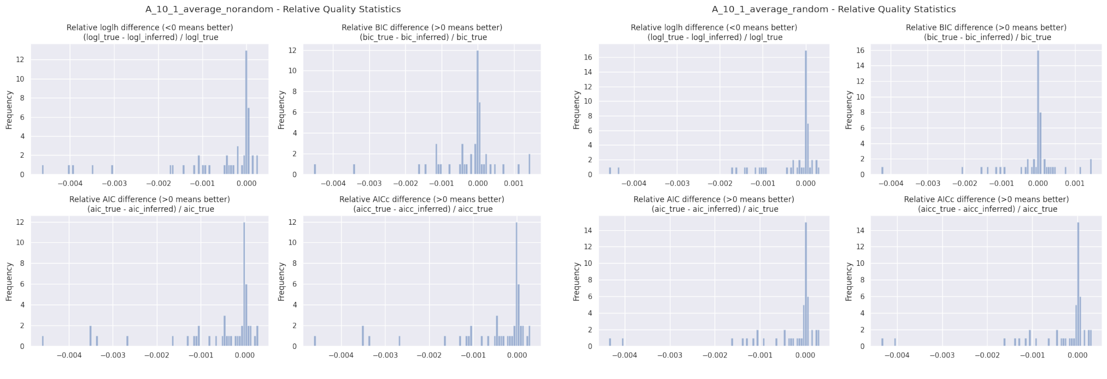


**Fig. 6.** Relative BIC, AIC, AICc, and loglikelihood differences for experiment A1, 10 taxa, 1 reticulation, `LhModel.AVERAGE`. Left: starting from RAxML-NG best tree, right: starting from 3 maximum parsimony and 3 random trees.

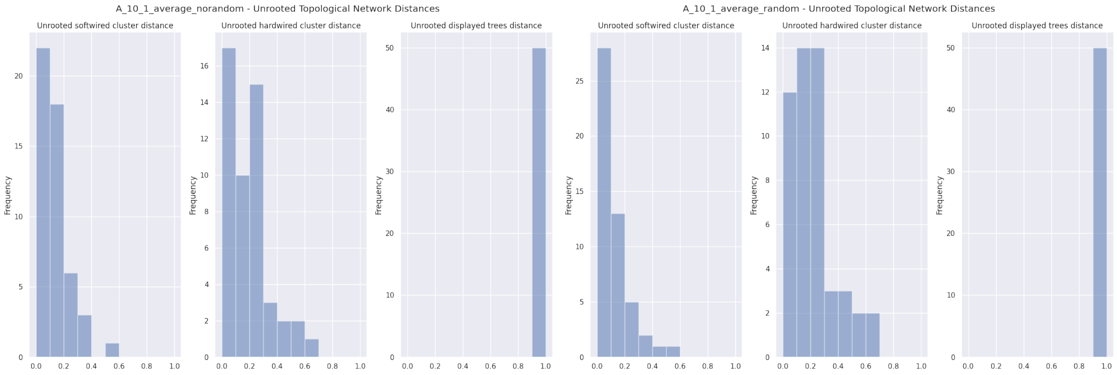


**Fig. 7.** Unrooted relative distances for experiment A1, 10 taxa, 1 reticulation, `LhModel.AVERAGE`. Left: starting from RAxML-NG best tree, right: starting from 3 maximum parsimony and 3 random trees.

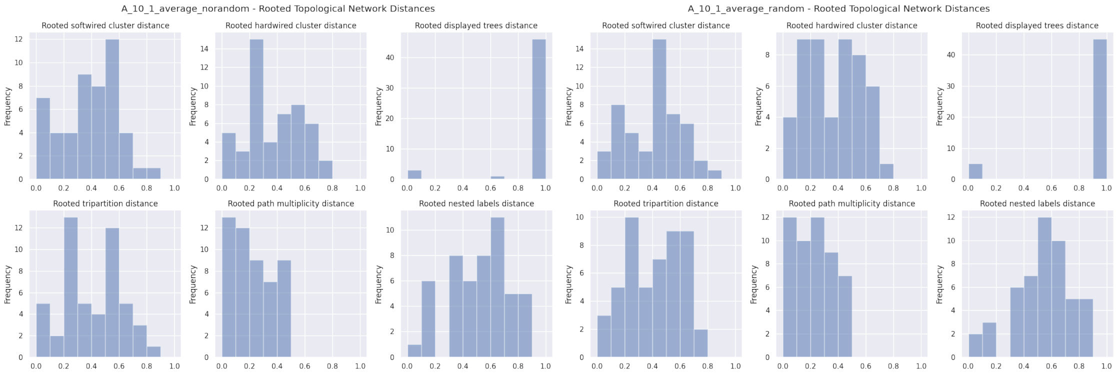


**Fig. 8.** Rooted relative distances for experiment A1, 10 taxa, 1 reticulation, `LhModel.AVERAGE`. Left: starting from RAxML-NG best tree, right: starting from 3 maximum parsimony and 3 random trees.

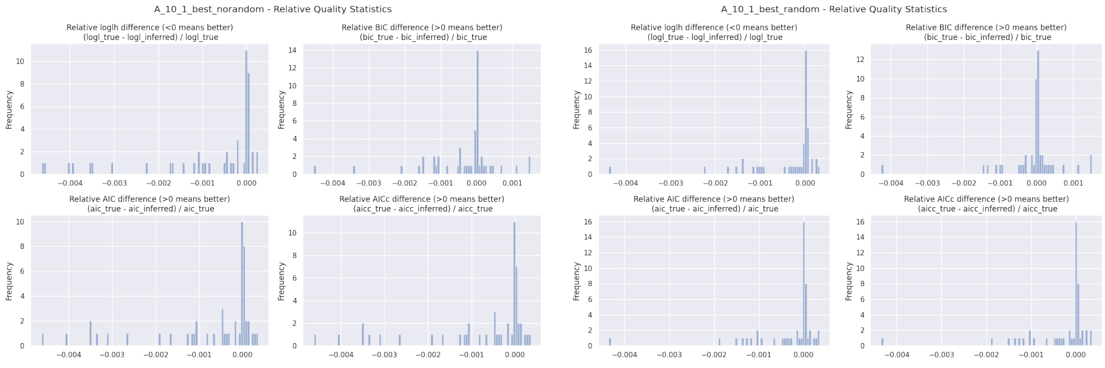


**Fig. 9.** Relative BIC, AIC, AICc, and loglikelihood differences for experiment A1, 10 taxa, 1 reticulation, `LhModel.BEST`. Left: starting from RAxML-NG best tree, right: starting from 3 maximum parsimony and 3 random trees.

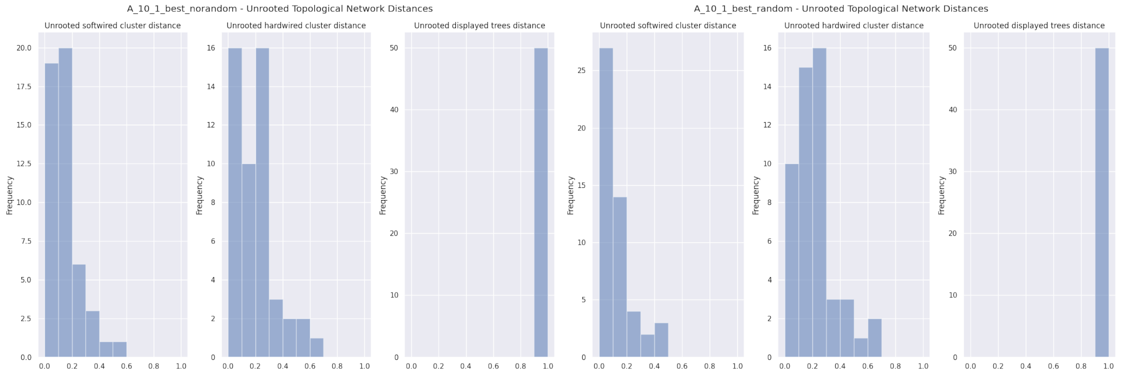


**Fig. 10.** Unrooted relative distances for experiment A1, 10 taxa, 1 reticulation, `LhModel.BEST`. Left: starting from RAxML-NG best tree, right: starting from 3 maximum parsimony and 3 random trees.

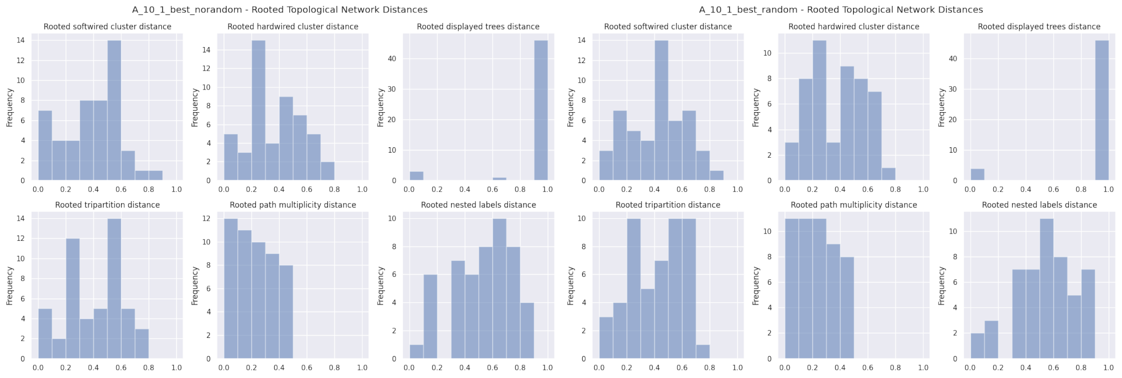


**Fig. 11.** Rooted relative distances for experiment A1, 10 taxa, 1 reticulation, `LhModel.BEST`. Left: starting from RAxML-NG best tree, right: starting from 3 maximum parsimony and 3 random trees.

| | count | mean | std | min | 10% | 25% | 50% | 75% | 90% | max |
|---|---|---|---|---|---|---|---|---|---|---|
| n_reticulations_inferred | 50.0 | 0.820000 | 0.388088 | 0.000000 | 0.000000 | 1.000000 | 1.000000 | 1.000000 | 1.000000 | 1.000000 |
| bic_diff | 50.0 | -4.420361 | 18.146684 | -71.946240 | -23.289644 | -8.825367 | -0.345645 | 0.707488 | 8.160995 | 39.836400 |
| bic_diff_relative | 50.0 | -0.000280 | 0.000986 | -0.004531 | -0.001156 | -0.000426 | -0.000016 | 0.000034 | 0.000368 | 0.001452 |
| aic_diff | 50.0 | -10.110872 | 18.584628 | -71.946230 | -29.855933 | -13.985325 | -0.794100 | 0.274478 | 1.858384 | 8.222450 |
| aic_diff_relative | 50.0 | -0.000573 | 0.001103 | -0.004623 | -0.001758 | -0.000618 | -0.000039 | 0.000015 | 0.000068 | 0.000303 |
| aicc_diff | 50.0 | -10.105313 | 18.580253 | -71.946240 | -29.852844 | -13.962157 | -0.794105 | 0.274477 | 1.886186 | 8.253330 |
| aicc_diff_relative | 50.0 | -0.000573 | 0.001103 | -0.004623 | -0.001758 | -0.000617 | -0.000039 | 0.000015 | 0.000069 | 0.000304 |
| lnL_diff | 50.0 | 5.775436 | 9.964811 | -2.248006 | -0.461930 | -0.040968 | 0.605118 | 7.999741 | 17.026374 | 35.973118 |
| lnL_diff_relative | 50.0 | -0.000649 | 0.001189 | -0.004647 | -0.001865 | -0.000929 | -0.000064 | 0.000006 | 0.000042 | 0.000261 |
| unrooted_softwired_network_distance | 50.0 | 0.100429 | 0.113179 | 0.000000 | 0.000000 | 0.000000 | 0.100000 | 0.125000 | 0.230000 | 0.500000 |
| unrooted_hardwired_network_distance | 50.0 | 0.169636 | 0.164495 | 0.000000 | 0.000000 | 0.000000 | 0.125000 | 0.250000 | 0.340000 | 0.636364 |
| unrooted_displayed_trees_distance | 50.0 | 1.000000 | 0.000000 | 1.000000 | 1.000000 | 1.000000 | 1.000000 | 1.000000 | 1.000000 | 1.000000 |
| rooted_softwired_network_distance | 50.0 | 0.386185 | 0.215836 | 0.000000 | 0.075000 | 0.255682 | 0.408333 | 0.543706 | 0.626786 | 0.800000 |
| rooted_hardwired_network_distance | 50.0 | 0.365423 | 0.211332 | 0.000000 | 0.100000 | 0.222222 | 0.363636 | 0.534091 | 0.666667 | 0.769231 |
| rooted_displayed_trees_distance | 50.0 | 0.933333 | 0.242810 | 0.000000 | 1.000000 | 1.000000 | 1.000000 | 1.000000 | 1.000000 | 1.000000 |
| rooted_tripartition_distance | 50.0 | 0.402553 | 0.225834 | 0.000000 | 0.100000 | 0.222222 | 0.409091 | 0.583333 | 0.692308 | 0.800000 |
| rooted_path_multiplicity_distance | 50.0 | 0.226851 | 0.133888 | 0.000000 | 0.085714 | 0.107143 | 0.204545 | 0.333333 | 0.400000 | 0.461538 |
| rooted_nested_labels_distance | 50.0 | 0.519740 | 0.213260 | 0.000000 | 0.181818 | 0.333333 | 0.571429 | 0.666667 | 0.755000 | 0.888889 |
| runtime_raxml | 50.0 | 3.502760 | 1.415170 | 2.020000 | 2.145200 | 2.336000 | 2.673000 | 4.636500 | 5.660900 | 6.402000 |
| runtime_inference | 50.0 | 6.042900 | 3.791490 | 1.145000 | 1.485900 | 4.649500 | 5.901000 | 7.673750 | 8.537000 | 24.928000 |

| | count | mean | std | min | 10% | 25% | 50% | 75% | 90% | max |
|---|---|---|---|---|---|---|---|---|---|---|
| n_reticulations_inferred | 50.0 | 0.860000 | 0.350510 | 0.000000 | 0.000000 | 1.000000 | 1.000000 | 1.000000 | 1.000000 | 1.000000 |
| bic_diff | 50.0 | -1.744506 | 15.646837 | -67.708330 | -16.933045 | -2.259210 | 0.028310 | 1.432960 | 8.294548 | 39.836400 |
| bic_diff_relative | 50.0 | -0.000142 | 0.000858 | -0.004264 | -0.001045 | -0.000125 | 0.000002 | 0.000067 | 0.000375 | 0.001452 |
| aic_diff | 50.0 | -6.170459 | 15.078599 | -67.708330 | -23.283828 | -7.363610 | -0.053580 | 0.684085 | 2.615876 | 8.222450 |
| aic_diff_relative | 50.0 | -0.000358 | 0.000916 | -0.004351 | -0.001175 | -0.000341 | -0.000003 | 0.000032 | 0.000150 | 0.000316 |
| aicc_diff | 50.0 | -6.166135 | 15.075190 | -67.708340 | -23.252928 | -7.340452 | -0.053575 | 0.684078 | 2.615876 | 8.253330 |
| aicc_diff_relative | 50.0 | -0.000358 | 0.000916 | -0.004351 | -0.001175 | -0.000341 | -0.000003 | 0.000032 | 0.000150 | 0.000316 |
| lnL_diff | 50.0 | 3.645230 | 8.091366 | -2.838573 | -0.890046 | -0.275918 | 0.056281 | 3.900423 | 13.421074 | 36.782846 |
| lnL_diff_relative | 50.0 | -0.000414 | 0.000982 | -0.004585 | -0.001379 | -0.000342 | -0.000005 | 0.000028 | 0.000085 | 0.000318 |
| unrooted_softwired_network_distance | 50.0 | 0.088048 | 0.130223 | 0.000000 | 0.000000 | 0.000000 | 0.000000 | 0.125000 | 0.230000 | 0.583333 |
| unrooted_hardwired_network_distance | 50.0 | 0.195242 | 0.170311 | 0.000000 | 0.000000 | 0.111111 | 0.125000 | 0.287500 | 0.454545 | 0.666667 |
| unrooted_displayed_trees_distance | 50.0 | 1.000000 | 0.000000 | 1.000000 | 1.000000 | 1.000000 | 1.000000 | 1.000000 | 1.000000 | 1.000000 |
| rooted_softwired_network_distance | 50.0 | 0.404319 | 0.212943 | 0.000000 | 0.110000 | 0.229167 | 0.428571 | 0.556490 | 0.687981 | 0.800000 |
| rooted_hardwired_network_distance | 50.0 | 0.358841 | 0.208132 | 0.000000 | 0.111111 | 0.143750 | 0.363636 | 0.500000 | 0.639394 | 0.714286 |
| rooted_displayed_trees_distance | 50.0 | 0.900000 | 0.303046 | 0.000000 | 0.900000 | 1.000000 | 1.000000 | 1.000000 | 1.000000 | 1.000000 |
| rooted_tripartition_distance | 50.0 | 0.399995 | 0.205284 | 0.000000 | 0.111111 | 0.300000 | 0.400000 | 0.573864 | 0.692308 | 0.714286 |
| rooted_path_multiplicity_distance | 50.0 | 0.232137 | 0.117744 | 0.000000 | 0.095238 | 0.181818 | 0.260870 | 0.326087 | 0.400000 | 0.440000 |
| rooted_nested_labels_distance | 50.0 | 0.538218 | 0.200741 | 0.000000 | 0.318182 | 0.461538 | 0.571429 | 0.666667 | 0.755000 | 0.823529 |
| runtime_inference | 50.0 | 43.306300 | 21.315874 | 15.107000 | 24.399800 | 31.963250 | 41.320000 | 51.325250 | 59.937700 | 161.661000 |

**Table 2.** Percentiles for experiment A1, 10 taxa, 1 reticulation, `LhModel.AVERAGE`. Top: starting from RAxML-NG best tree, bottom: starting from 3 maximum parsimony and 3 random trees.

| | count | mean | std | min | 10% | 25% | 50% | 75% | 90% | max |
|---|---|---|---|---|---|---|---|---|---|---|
| n_reticulations_inferred | 50.0 | 0.760000 | 0.431419 | 0.000000 | 0.000000 | 1.000000 | 1.000000 | 1.000000 | 1.000000 | 1.000000 |
| bic_diff | 50.0 | -5.650328 | 18.994381 | -71.946240 | -25.182266 | -11.398798 | -0.149500 | 0.707488 | 8.160995 | 41.209700 |
| bic_diff_relative | 50.0 | -0.000348 | 0.001032 | -0.004531 | -0.001458 | -0.000497 | -0.000008 | 0.000034 | 0.000368 | 0.001502 |
| aic_diff | 50.0 | -13.237675 | 21.690441 | -71.946230 | -55.005788 | -19.029248 | -0.898900 | 0.336075 | 1.862326 | 9.595740 |
| aic_diff_relative | 50.0 | -0.000742 | 0.001265 | -0.004623 | -0.003150 | -0.001058 | -0.000039 | 0.000020 | 0.000068 | 0.000354 |
| aicc_diff | 50.0 | -13.230263 | 21.683785 | -71.946240 | -54.974899 | -19.029257 | -0.898910 | 0.336077 | 1.890128 | 9.626630 |
| aicc_diff_relative | 50.0 | -0.000741 | 0.001264 | -0.004623 | -0.003149 | -0.001057 | -0.000039 | 0.000020 | 0.000069 | 0.000355 |
| lnL_diff | 50.0 | 7.578838 | 11.802719 | -2.248006 | -0.609437 | -0.136858 | 1.112374 | 11.084812 | 31.378417 | 36.782846 |
| lnL_diff_relative | 50.0 | -0.000843 | 0.001380 | -0.004647 | -0.003511 | -0.001101 | -0.000121 | 0.000015 | 0.000059 | 0.000261 |
| unrooted_softwired_network_distance | 50.0 | 0.113429 | 0.117795 | 0.000000 | 0.000000 | 0.000000 | 0.105556 | 0.125000 | 0.300769 | 0.500000 |
| unrooted_hardwired_network_distance | 50.0 | 0.174636 | 0.163027 | 0.000000 | 0.000000 | 0.000000 | 0.125000 | 0.250000 | 0.340000 | 0.636364 |
| unrooted_displayed_trees_distance | 50.0 | 1.000000 | 0.000000 | 1.000000 | 1.000000 | 1.000000 | 1.000000 | 1.000000 | 1.000000 | 1.000000 |
| rooted_softwired_network_distance | 50.0 | 0.386995 | 0.211252 | 0.000000 | 0.075000 | 0.279545 | 0.416667 | 0.538462 | 0.602500 | 0.800000 |
| rooted_hardwired_network_distance | 50.0 | 0.358850 | 0.205181 | 0.000000 | 0.100000 | 0.222222 | 0.363636 | 0.500000 | 0.666667 | 0.769231 |
| rooted_displayed_trees_distance | 50.0 | 0.933333 | 0.242810 | 0.000000 | 1.000000 | 1.000000 | 1.000000 | 1.000000 | 1.000000 | 1.000000 |
| rooted_tripartition_distance | 50.0 | 0.406038 | 0.218661 | 0.000000 | 0.100000 | 0.241667 | 0.454545 | 0.583333 | 0.692308 | 0.769231 |
| rooted_path_multiplicity_distance | 50.0 | 0.228891 | 0.128696 | 0.000000 | 0.085714 | 0.152597 | 0.227273 | 0.326087 | 0.400000 | 0.461538 |
| rooted_nested_labels_distance | 50.0 | 0.524819 | 0.209380 | 0.000000 | 0.181818 | 0.333333 | 0.571429 | 0.666667 | 0.750000 | 0.888889 |
| runtime_raxml | 50.0 | 3.502760 | 1.415170 | 2.020000 | 2.145200 | 2.336000 | 2.673000 | 4.636500 | 5.660900 | 6.402000 |
| runtime_inference | 50.0 | 4.242960 | 2.932006 | 0.833000 | 1.009300 | 2.647500 | 4.180000 | 5.174500 | 6.696200 | 18.073000 |

| | count | mean | std | min | 10% | 25% | 50% | 75% | 90% | max |
|---|---|---|---|---|---|---|---|---|---|---|
| n_reticulations_inferred | 50.0 | 0.860000 | 0.350510 | 0.000000 | 0.000000 | 1.000000 | 1.000000e+00 | 1.000000 | 1.000000 | 1.000000 |
| bic_diff | 50.0 | -1.218126 | 15.049197 | -67.708330 | -16.131187 | -2.139918 | 5.551500e-02 | 1.432960 | 8.294548 | 41.209700 |
| bic_diff_relative | 50.0 | -0.000107 | 0.000813 | -0.004264 | -0.000931 | -0.000120 | 3.087947e-06 | 0.000067 | 0.000375 | 0.001502 |
| aic_diff | 50.0 | -5.644079 | 13.535077 | -67.708330 | -23.283828 | -7.363610 | -1.276500e-02 | 0.684085 | 2.615876 | 9.595740 |
| aic_diff_relative | 50.0 | -0.000311 | 0.000777 | -0.004351 | -0.001175 | -0.000341 | -7.099275e-07 | 0.000032 | 0.000150 | 0.000354 |
| aicc_diff | 50.0 | -5.639756 | 13.532211 | -67.708340 | -23.252928 | -7.340452 | -1.276500e-02 | 0.684078 | 2.615876 | 9.626630 |
| aicc_diff_relative | 50.0 | -0.000311 | 0.000777 | -0.004351 | -0.001175 | -0.000341 | -7.099209e-07 | 0.000032 | 0.000150 | 0.000355 |
| lnL_diff | 50.0 | 3.382040 | 7.266034 | -2.838573 | -0.890046 | -0.305445 | 2.678750e-02 | 3.900267 | 12.822320 | 33.854167 |
| lnL_diff_relative | 50.0 | -0.000364 | 0.000821 | -0.004373 | -0.001379 | -0.000342 | -2.852233e-06 | 0.000030 | 0.000085 | 0.000318 |
| unrooted_softwired_network_distance | 50.0 | 0.088326 | 0.121830 | 0.000000 | 0.000000 | 0.000000 | 0.000000e+00 | 0.125000 | 0.230769 | 0.454545 |
| unrooted_hardwired_network_distance | 50.0 | 0.197187 | 0.160398 | 0.000000 | 0.000000 | 0.111111 | 1.736111e-01 | 0.250000 | 0.405455 | 0.666667 |
| unrooted_displayed_trees_distance | 50.0 | 1.000000 | 0.000000 | 1.000000 | 1.000000 | 1.000000 | 1.000000e+00 | 1.000000 | 1.000000 | 1.000000 |
| rooted_softwired_network_distance | 50.0 | 0.415222 | 0.211019 | 0.000000 | 0.110000 | 0.279545 | 4.285714e-01 | 0.581801 | 0.692308 | 0.800000 |
| rooted_hardwired_network_distance | 50.0 | 0.373905 | 0.205427 | 0.000000 | 0.111111 | 0.241667 | 3.818182e-01 | 0.500000 | 0.666667 | 0.714286 |
| rooted_displayed_trees_distance | 50.0 | 0.920000 | 0.274048 | 0.000000 | 1.000000 | 1.000000 | 1.000000e+00 | 1.000000 | 1.000000 | 1.000000 |
| rooted_tripartition_distance | 50.0 | 0.410008 | 0.206137 | 0.000000 | 0.111111 | 0.300000 | 4.272727e-01 | 0.583333 | 0.692308 | 0.714286 |
| rooted_path_multiplicity_distance | 50.0 | 0.239388 | 0.121492 | 0.000000 | 0.095238 | 0.181818 | 2.608696e-01 | 0.333333 | 0.400000 | 0.461538 |
| rooted_nested_labels_distance | 50.0 | 0.541593 | 0.211115 | 0.000000 | 0.318182 | 0.461538 | 5.714286e-01 | 0.666667 | 0.800000 | 0.888889 |
| runtime_inference | 50.0 | 32.599280 | 16.695448 | 12.193000 | 17.566400 | 22.769250 | 3.042050e+01 | 38.626500 | 44.539600 | 122.241000 |

**Table 3.** Percentiles for experiment A1, 10 taxa, 1 reticulation, `LhModel.BEST`. Top: starting from RAxML-NG best tree, bottom: starting from 3 maximum parsimony and 3 random trees.

#### 6.2 Multiple starting trees, 20 taxa, 2 reticulations

In this experiment, we simulated 50 networks each for 20 taxa and 2 reticulation. In addition to the NetRAX inference starting from the RAxML-NG best ML tree, we also started another NetRAX inference using 3 random and 3 maximum parsimony starting trees.

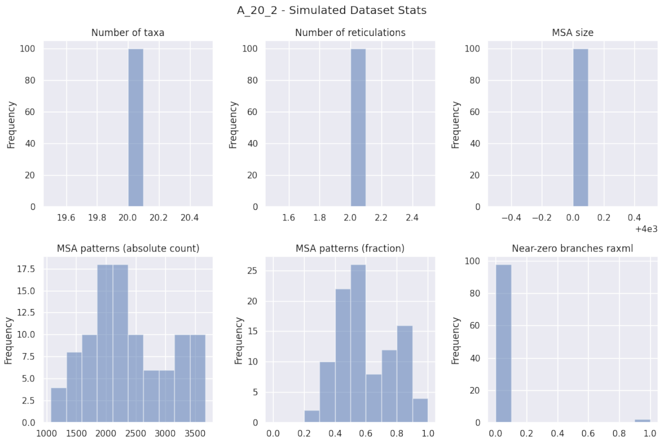


**Fig. 12.** Simulated dataset statistics for experiment A1, 20 taxa, 2 reticulations.

| A_20_2_norandom | LhType.AVERAGE | LhType.BEST |
|---|---|---|
| Inferred BIC better or equal | 4 (8.00 %) | 5 (10.00 %) |
| Inferred AIC better or equal | 3 (6.00 %) | 4 (8.00 %) |
| Inferred AICc better or equal | 3 (6.00 %) | 4 (8.00 %) |
| Inferred BIC worse | 46 (92.00 %) | 45 (90.00 %) |
| Inferred AIC worse | 47 (94.00 %) | 46 (92.00 %) |
| Inferred AICc worse | 47 (94.00 %) | 46 (92.00 %) |
| Inferred lnL better or equal | 3 (6.00 %) | 4 (8.00 %) |
| Inferred lnL worse | 47 (94.00 %) | 46 (92.00 %) |
| Inferred n_reticulations less | 4 (8.00 %) | 3 (6.00 %) |
| Inferred n_reticulations equal | 46 (92.00 %) | 47 (94.00 %) |
| Inferred n_reticulations more | 0 (0.00 %) | 0 (0.00 %) |
| Unrooted softwired distance zero | 26 (52.00 %) | 28 (56.00 %) |
| Good result | 28 (56.00 %) | 31 (62.00 %) |
| Passable result | 19 (38.00 %) | 17 (34.00 %) |
| Bad result | 3 (6.00 %) | 2 (4.00 %) |

| A_20_2_random | LhType.AVERAGE | LhType.BEST |
|---|---|---|
| Inferred BIC better or equal | 13 (26.00 %) | 14 (28.00 %) |
| Inferred AIC better or equal | 12 (24.00 %) | 13 (26.00 %) |
| Inferred AICc better or equal | 12 (24.00 %) | 13 (26.00 %) |
| Inferred BIC worse | 37 (74.00 %) | 36 (72.00 %) |
| Inferred AIC worse | 38 (76.00 %) | 37 (74.00 %) |
| Inferred AICc worse | 38 (76.00 %) | 37 (74.00 %) |
| Inferred lnL better or equal | 12 (24.00 %) | 13 (26.00 %) |
| Inferred lnL worse | 38 (76.00 %) | 37 (74.00 %) |
| Inferred n_reticulations less | 3 (6.00 %) | 3 (6.00 %) |
| Inferred n_reticulations equal | 47 (94.00 %) | 47 (94.00 %) |
| Inferred n_reticulations more | 0 (0.00 %) | 0 (0.00 %) |
| Unrooted softwired distance zero | 29 (58.00 %) | 28 (56.00 %) |
| Good result | 35 (70.00 %) | 35 (70.00 %) |
| Passable result | 13 (26.00 %) | 13 (26.00 %) |
| Bad result | 2 (4.00 %) | 2 (4.00 %) |

**Table 4.** Summary statistics for experiment A1, 29 taxa, 2 reticulations. Left: starting from RAxML-NG best tree, right: starting from 3 maximum parsimony and 3 random trees.

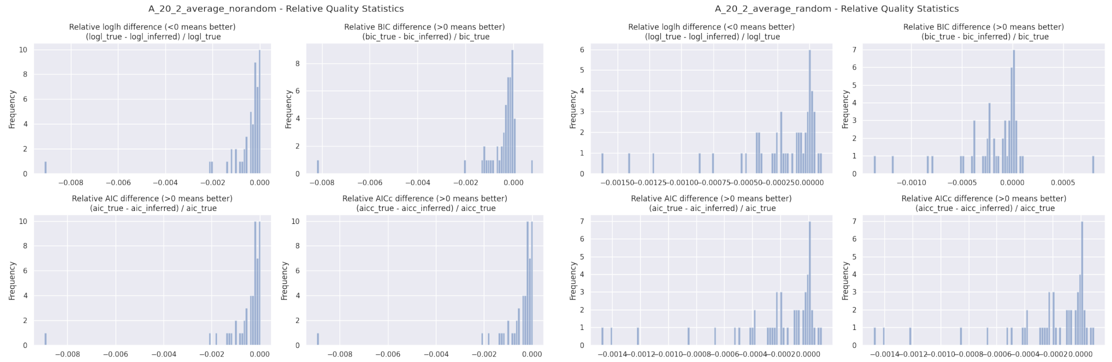


**Fig. 13.** Relative BIC, AIC, AICc, and loglikelihood differences for experiment A1, 20 taxa, 2 reticulations, `LhModel.AVERAGE`. Left: starting from RAxML-NG best tree, right: starting from 3 maximum parsimony and 3 random trees.

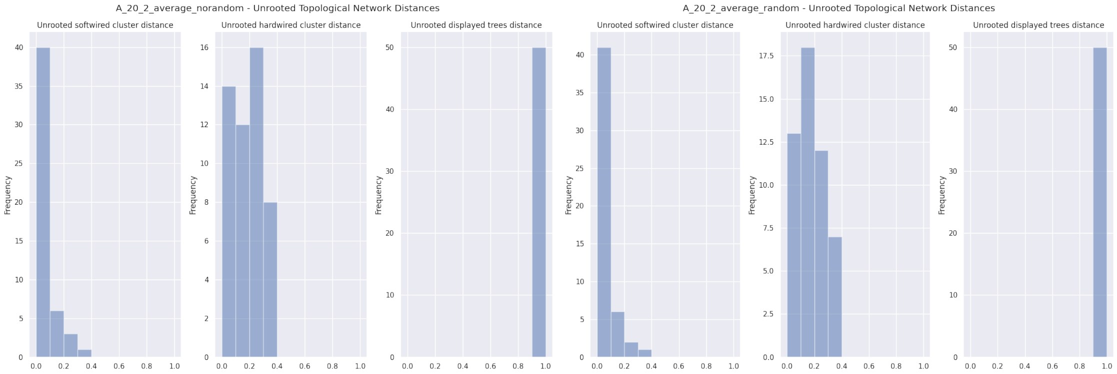


**Fig. 14.** Unrooted relative distances for experiment A1, 20 taxa, 2 reticulations, `LhModel.AVERAGE`. Left: starting from RAxML-NG best tree, right: starting from 3 maximum parsimony and 3 random trees.

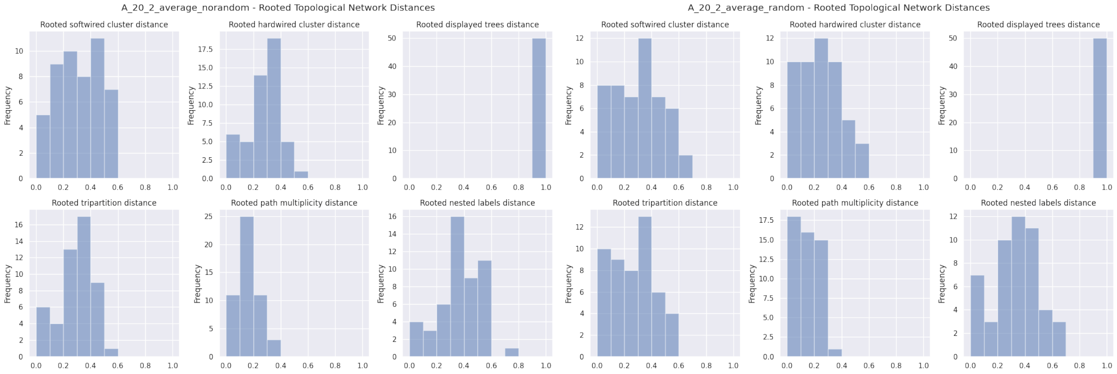


**Fig. 15.** Rooted relative distances for experiment A1, 20 taxa, 2 reticulations, `LhModel.AVERAGE`. Left: starting from RAxML-NG best tree, right: starting from 3 maximum parsimony and 3 random trees.

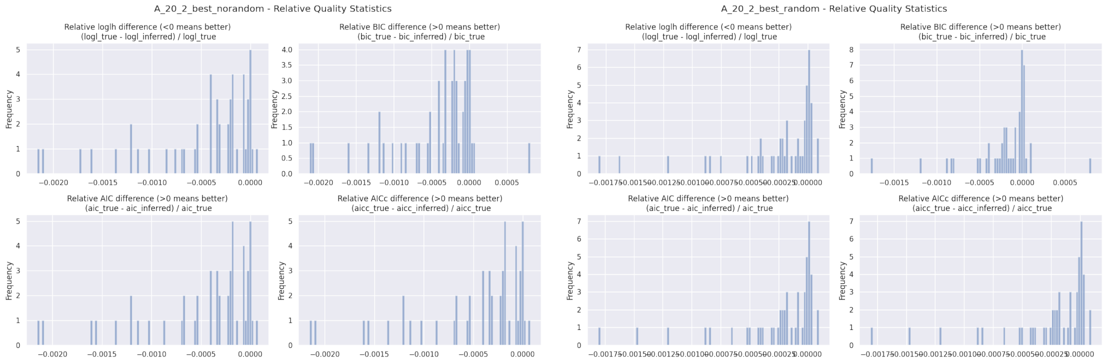


**Fig. 16.** Relative BIC, AIC, AICc, and loglikelihood differences for experiment A1, 20 taxa, 2 reticulations, `LhModel.BEST`. Left: starting from RAxML-NG best tree, right: starting from 3 maximum parsimony and 3 random trees.

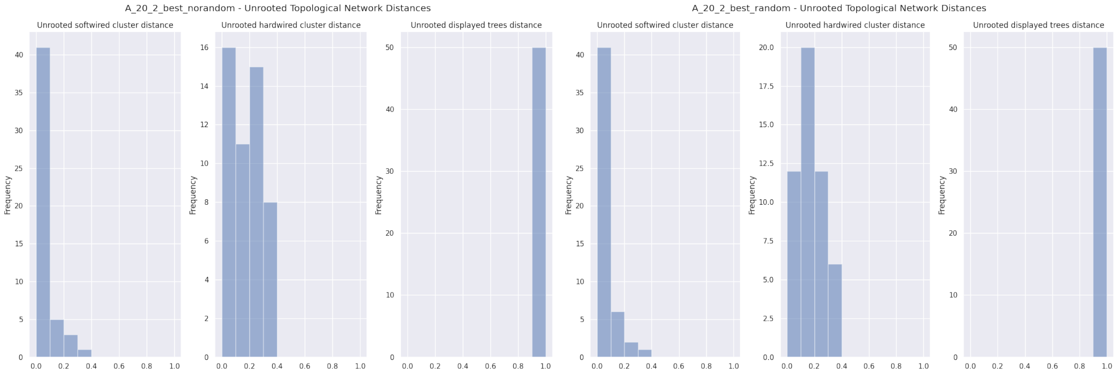


**Fig. 17.** Unrooted relative distances for experiment A1, 20 taxa, 2 reticulations, `LhModel.BEST`. Left: starting from RAxML-NG best tree, right: starting from 3 maximum parsimony and 3 random trees.

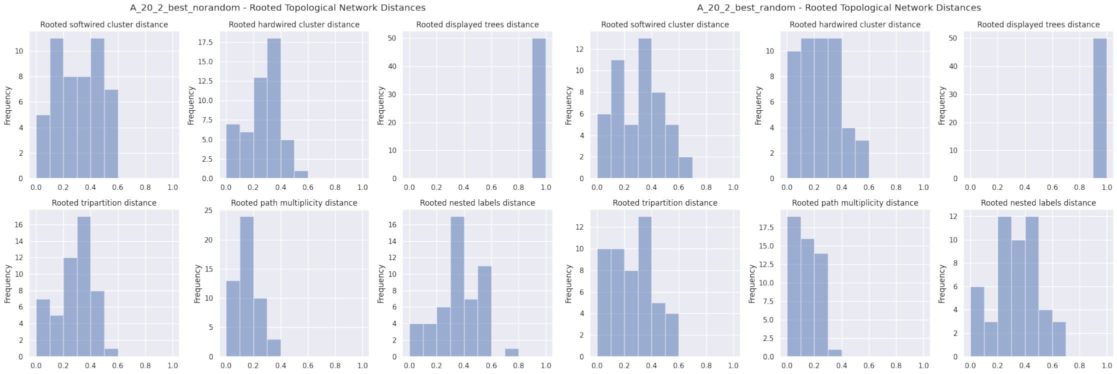


**Fig. 18.** Rooted relative distances for experiment A1, 20 taxa, 2 reticulations, `LhModel.BEST`. Left: starting from RAxML-NG best tree, right: starting from 3 maximum parsimony and 3 random trees.

| | count | mean | std | min | 10% | 25% | 50% | 75% | 90% | max |
|---|---|---|---|---|---|---|---|---|---|---|
| n_reticulations_inferred | 50.0 | 1.920000 | 0.274048 | 1.000000 | 2.000000 | 2.000000 | 2.000000 | 2.000000 | 2.000000 | 2.000000 |
| bic_diff | 50.0 | -32.749270 | 72.413860 | -487.196230 | -66.180779 | -30.159462 | -16.694520 | -5.641420 | -0.341186 | 30.193600 |
| bic_diff_relative | 50.0 | -0.000517 | 0.001204 | -0.008214 | -0.001176 | -0.000522 | -0.000222 | -0.000070 | -0.000006 | 0.000807 |
| aic_diff | 50.0 | -35.722001 | 77.274469 | -524.355360 | -66.352113 | -34.832762 | -16.694515 | -6.116095 | -0.756531 | 1.741200 |
| aic_diff_relative | 50.0 | -0.000578 | 0.001299 | -0.008955 | -0.001216 | -0.000549 | -0.000225 | -0.000088 | -0.000013 | 0.000026 |
| aicc_diff | 50.0 | -35.720710 | 77.272135 | -524.339230 | -66.352114 | -34.832770 | -16.694520 | -6.112055 | -0.756532 | 1.741200 |
| aicc_diff_relative | 50.0 | -0.000578 | 0.001299 | -0.008955 | -0.001216 | -0.000549 | -0.000225 | -0.000088 | -0.000013 | 0.000026 |
| lnL_diff | 50.0 | 18.181001 | 39.227272 | -0.870600 | 0.378262 | 3.098370 | 8.347260 | 17.416385 | 33.176057 | 266.177670 |
| lnL_diff_relative | 50.0 | -0.000592 | 0.001323 | -0.009118 | -0.001220 | -0.000551 | -0.000228 | -0.000088 | -0.000013 | 0.000026 |
| unrooted_softwired_network_distance | 50.0 | 0.050142 | 0.076995 | 0.000000 | 0.000000 | 0.000000 | 0.000000 | 0.068391 | 0.163231 | 0.323529 |
| unrooted_hardwired_network_distance | 50.0 | 0.171090 | 0.114029 | 0.000000 | 0.000000 | 0.063283 | 0.190476 | 0.272727 | 0.319697 | 0.347826 |
| unrooted_displayed_trees_distance | 50.0 | 1.000000 | 0.000000 | 1.000000 | 1.000000 | 1.000000 | 1.000000 | 1.000000 | 1.000000 | 1.000000 |
| rooted_softwired_network_distance | 50.0 | 0.303446 | 0.156742 | 0.000000 | 0.101437 | 0.185185 | 0.312229 | 0.428571 | 0.511220 | 0.595238 |
| rooted_hardwired_network_distance | 50.0 | 0.267436 | 0.114829 | 0.000000 | 0.095238 | 0.227273 | 0.282609 | 0.333333 | 0.400000 | 0.538462 |
| rooted_displayed_trees_distance | 50.0 | 1.000000 | 0.000000 | 1.000000 | 1.000000 | 1.000000 | 1.000000 | 1.000000 | 1.000000 | 1.000000 |
| rooted_tripartition_distance | 50.0 | 0.285602 | 0.121471 | 0.000000 | 0.095238 | 0.227273 | 0.333333 | 0.364583 | 0.416667 | 0.555556 |
| rooted_path_multiplicity_distance | 50.0 | 0.161847 | 0.075801 | 0.000000 | 0.047619 | 0.136364 | 0.177778 | 0.213043 | 0.257447 | 0.326531 |
| rooted_nested_labels_distance | 50.0 | 0.369169 | 0.155558 | 0.000000 | 0.166667 | 0.256923 | 0.370370 | 0.482759 | 0.533333 | 0.777778 |
| runtime_raxml | 50.0 | 14.842580 | 4.011155 | 7.775000 | 10.976800 | 11.905250 | 13.959000 | 17.272000 | 20.114400 | 25.962000 |
| runtime_inference | 50.0 | 130.985120 | 66.529495 | 19.802000 | 67.308700 | 91.242750 | 120.047000 | 156.379250 | 199.049800 | 389.484000 |

| | count | mean | std | min | 10% | 25% | 50% | 75% | 90% | max |
|---|---|---|---|---|---|---|---|---|---|---|
| n_reticulations_inferred | 50.0 | 1.940000 | 0.239898 | 1.000000 | 2.000000 | 2.000000 | 2.000000 | 2.000000e+00 | 2.000000 | 2.000000 |
| bic_diff | 50.0 | -10.604955 | 16.690075 | -63.082280 | -26.629026 | -17.398125 | -5.738295 | 1.980500e-02 | 1.930973 | 30.193610 |
| bic_diff_relative | 50.0 | -0.000178 | 0.000335 | -0.001373 | -0.000491 | -0.000257 | -0.000084 | 3.142927e-07 | 0.000030 | 0.000807 |
| aic_diff | 50.0 | -12.834502 | 19.009067 | -82.257800 | -32.535050 | -17.398115 | -6.240205 | -3.773175e-01 | 1.557329 | 5.538010 |
| aic_diff_relative | 50.0 | -0.000223 | 0.000353 | -0.001478 | -0.000542 | -0.000267 | -0.000092 | -6.467662e-06 | 0.000018 | 0.000096 |
| aicc_diff | 50.0 | -12.833535 | 19.007217 | -82.241680 | -32.535042 | -17.398125 | -6.240205 | -3.773175e-01 | 1.557329 | 5.538010 |
| aicc_diff_relative | 50.0 | -0.000223 | 0.000352 | -0.001478 | -0.000542 | -0.000267 | -0.000092 | -6.467643e-06 | 0.000018 | 0.000096 |
| lnL_diff | 50.0 | 6.657252 | 9.999492 | -2.769000 | -0.778669 | 0.188660 | 3.120105 | 8.699058e+00 | 16.267525 | 45.128910 |
| lnL_diff_relative | 50.0 | -0.000233 | 0.000368 | -0.001626 | -0.000553 | -0.000284 | -0.000092 | -6.485812e-06 | 0.000018 | 0.000096 |
| unrooted_softwired_network_distance | 50.0 | 0.043575 | 0.074555 | 0.000000 | 0.000000 | 0.000000 | 0.000000 | 4.347826e-02 | 0.125833 | 0.323529 |
| unrooted_hardwired_network_distance | 50.0 | 0.169085 | 0.114350 | 0.000000 | 0.000000 | 0.096429 | 0.186147 | 2.443182e-01 | 0.321146 | 0.391304 |
| unrooted_displayed_trees_distance | 50.0 | 1.000000 | 0.000000 | 1.000000 | 1.000000 | 1.000000 | 1.000000 | 1.000000e+00 | 1.000000 | 1.000000 |
| rooted_softwired_network_distance | 50.0 | 0.294714 | 0.177157 | 0.000000 | 0.033333 | 0.164118 | 0.319659 | 4.265625e-01 | 0.524762 | 0.617021 |
| rooted_hardwired_network_distance | 50.0 | 0.242747 | 0.144391 | 0.000000 | 0.045000 | 0.142857 | 0.244071 | 3.645833e-01 | 0.416667 | 0.560000 |
| rooted_displayed_trees_distance | 50.0 | 1.000000 | 0.000000 | 1.000000 | 1.000000 | 1.000000 | 1.000000 | 1.000000e+00 | 1.000000 | 1.000000 |
| rooted_tripartition_distance | 50.0 | 0.255407 | 0.150930 | 0.000000 | 0.045000 | 0.152597 | 0.260870 | 3.645833e-01 | 0.421154 | 0.592593 |
| rooted_path_multiplicity_distance | 50.0 | 0.147934 | 0.088537 | 0.000000 | 0.042857 | 0.093023 | 0.136364 | 2.173913e-01 | 0.255319 | 0.380000 |
| rooted_nested_labels_distance | 50.0 | 0.331557 | 0.162499 | 0.000000 | 0.086957 | 0.240000 | 0.307692 | 4.285714e-01 | 0.535172 | 0.666667 |
| runtime_inference | 50.0 | 1012.825840 | 642.587280 | 233.054000 | 577.440900 | 723.144000 | 926.345500 | 1.102426e+03 | 1330.331200 | 4703.129000 |

**Table 5.** Percentiles for experiment A1, 20 taxa, 2 reticulations, `LhModel.AVERAGE`. Top: starting from RAxML-NG best tree, bottom: starting from 3 maximum parsimony and 3 random trees.

| | count | mean | std | min | 10% | 25% | 50% | 75% | 90% | max |
|---|---|---|---|---|---|---|---|---|---|---|
| n_reticulations_inferred | 50.0 | 1.940000 | 0.239898 | 1.000000 | 2.000000 | 2.000000 | 2.000000 | 2.000000 | 2.000000 | 2.000000 |
| bic_diff | 50.0 | -26.975549 | 35.470443 | -172.916000 | -63.675523 | -38.109370 | -18.151355 | -3.854223 | -0.225679 | 30.193650 |
| bic_diff_relative | 50.0 | -0.000416 | 0.000553 | -0.002112 | -0.001189 | -0.000541 | -0.000223 | -0.000066 | -0.000002 | 0.000807 |
| aic_diff | 50.0 | -29.205098 | 35.693534 | -172.916010 | -69.443798 | -41.411020 | -18.151350 | -5.059700 | -0.341177 | 4.171590 |
| aic_diff_relative | 50.0 | -0.000464 | 0.000552 | -0.002153 | -0.001228 | -0.000636 | -0.000228 | -0.000070 | -0.000006 | 0.000075 |
| aicc_diff | 50.0 | -29.204129 | 35.692955 | -172.916000 | -69.443806 | -41.411027 | -18.151355 | -5.059698 | -0.341186 | 4.171580 |
| aicc_diff_relative | 50.0 | -0.000463 | 0.000552 | -0.002153 | -0.001228 | -0.000636 | -0.000228 | -0.000070 | -0.000006 | 0.000075 |
| lnL_diff | 50.0 | 14.842549 | 18.015534 | -2.085790 | 0.170597 | 2.529855 | 9.075675 | 20.705510 | 34.721903 | 86.458000 |
| lnL_diff_relative | 50.0 | -0.000474 | 0.000559 | -0.002162 | -0.001232 | -0.000638 | -0.000273 | -0.000070 | -0.000006 | 0.000075 |
| unrooted_softwired_network_distance | 50.0 | 0.045153 | 0.074779 | 0.000000 | 0.000000 | 0.000000 | 0.000000 | 0.060870 | 0.128500 | 0.323529 |
| unrooted_hardwired_network_distance | 50.0 | 0.165683 | 0.114727 | 0.000000 | 0.000000 | 0.050658 | 0.190476 | 0.264069 | 0.319697 | 0.347826 |
| unrooted_displayed_trees_distance | 50.0 | 1.000000 | 0.000000 | 1.000000 | 1.000000 | 1.000000 | 1.000000 | 1.000000 | 1.000000 | 1.000000 |
| rooted_softwired_network_distance | 50.0 | 0.299620 | 0.159197 | 0.000000 | 0.101437 | 0.174345 | 0.312229 | 0.428571 | 0.511220 | 0.595238 |
| rooted_hardwired_network_distance | 50.0 | 0.261673 | 0.117804 | 0.000000 | 0.095238 | 0.186364 | 0.260870 | 0.333333 | 0.400000 | 0.538462 |
| rooted_displayed_trees_distance | 50.0 | 1.000000 | 0.000000 | 1.000000 | 1.000000 | 1.000000 | 1.000000 | 1.000000 | 1.000000 | 1.000000 |
| rooted_tripartition_distance | 50.0 | 0.277125 | 0.123054 | 0.000000 | 0.095238 | 0.227273 | 0.318841 | 0.333333 | 0.401667 | 0.555556 |
| rooted_path_multiplicity_distance | 50.0 | 0.157933 | 0.077783 | 0.000000 | 0.047619 | 0.103858 | 0.177778 | 0.207488 | 0.257447 | 0.326531 |
| rooted_nested_labels_distance | 50.0 | 0.360012 | 0.157165 | 0.000000 | 0.166667 | 0.240000 | 0.370370 | 0.482759 | 0.533333 | 0.777778 |
| runtime_raxml | 50.0 | 14.842580 | 4.011155 | 7.775000 | 10.976800 | 11.905250 | 13.959000 | 17.272000 | 20.114400 | 25.962000 |
| runtime_inference | 50.0 | 108.859280 | 58.129259 | 14.173000 | 50.402000 | 69.631750 | 100.586000 | 129.810750 | 171.611100 | 361.816000 |

| | count | mean | std | min | 10% | 25% | 50% | 75% | 90% | max |
|---|---|---|---|---|---|---|---|---|---|---|
| n_reticulations_inferred | 50.0 | 1.940000 | 0.239898 | 1.000000 | 2.000000 | 2.000000 | 2.000000 | 2.000000e+00 | 2.000000 | 2.000000 |
| bic_diff | 50.0 | -12.128115 | 19.911911 | -73.436460 | -31.691011 | -19.046660 | -4.776710 | 1.849150e-01 | 2.244084 | 30.193660 |
| bic_diff_relative | 50.0 | -0.000198 | 0.000381 | -0.001772 | -0.000547 | -0.000256 | -0.000084 | 4.087443e-06 | 0.000032 | 0.000807 |
| aic_diff | 50.0 | -14.357664 | 21.733376 | -82.257800 | -53.811095 | -19.794323 | -5.109795 | 1.980500e-02 | 1.930973 | 5.538010 |
| aic_diff_relative | 50.0 | -0.000243 | 0.000396 | -0.001807 | -0.000683 | -0.000282 | -0.000092 | 3.180079e-07 | 0.000030 | 0.000096 |
| aicc_diff | 50.0 | -14.356697 | 21.731825 | -82.241680 | -53.809491 | -19.794297 | -5.109785 | 1.980500e-02 | 1.930973 | 5.538010 |
| aicc_diff_relative | 50.0 | -0.000243 | 0.000396 | -0.001807 | -0.000683 | -0.000282 | -0.000092 | 3.180071e-07 | 0.000030 | 0.000096 |
| lnL_diff | 50.0 | 7.418833 | 11.285635 | -2.769000 | -0.965486 | -0.009903 | 2.554895 | 9.897157e+00 | 27.246850 | 45.128910 |
| lnL_diff_relative | 50.0 | -0.000253 | 0.000410 | -0.001814 | -0.000769 | -0.000312 | -0.000092 | 3.188187e-07 | 0.000030 | 0.000097 |
| unrooted_softwired_network_distance | 50.0 | 0.045714 | 0.074944 | 0.000000 | 0.000000 | 0.000000 | 0.000000 | 6.086957e-02 | 0.129167 | 0.323529 |
| unrooted_hardwired_network_distance | 50.0 | 0.170981 | 0.104951 | 0.000000 | 0.050000 | 0.100000 | 0.181818 | 2.272727e-01 | 0.318182 | 0.391304 |
| unrooted_displayed_trees_distance | 50.0 | 1.000000 | 0.000000 | 1.000000 | 1.000000 | 1.000000 | 1.000000 | 1.000000e+00 | 1.000000 | 1.000000 |
| rooted_softwired_network_distance | 50.0 | 0.297809 | 0.167512 | 0.000000 | 0.095097 | 0.164071 | 0.314145 | 4.265625e-01 | 0.514309 | 0.617021 |
| rooted_hardwired_network_distance | 50.0 | 0.241865 | 0.140233 | 0.000000 | 0.050000 | 0.142857 | 0.232684 | 3.333333e-01 | 0.416667 | 0.560000 |
| rooted_displayed_trees_distance | 50.0 | 1.000000 | 0.000000 | 1.000000 | 1.000000 | 1.000000 | 1.000000 | 1.000000e+00 | 1.000000 | 1.000000 |
| rooted_tripartition_distance | 50.0 | 0.253856 | 0.145855 | 0.000000 | 0.050000 | 0.152597 | 0.260870 | 3.333333e-01 | 0.421154 | 0.592593 |
| rooted_path_multiplicity_distance | 50.0 | 0.148259 | 0.084708 | 0.000000 | 0.047619 | 0.093023 | 0.136364 | 2.173913e-01 | 0.255319 | 0.380000 |
| rooted_nested_labels_distance | 50.0 | 0.339758 | 0.157793 | 0.086957 | 0.086957 | 0.240000 | 0.370370 | 4.692118e-01 | 0.535172 | 0.666667 |
| runtime_inference | 50.0 | 872.722040 | 498.178203 | 186.327000 | 486.574700 | 622.678750 | 803.013000 | 1.020157e+03 | 1232.181100 | 3430.972000 |

**Table 6.** Percentiles for experiment A1, 20 taxa, 2 reticulations, `LhModel.BEST`. Top: starting from RAxML-NG best tree, bottom: starting from 3 maximum parsimony and 3 random trees.

##### 6.3 A1: Multiple starting trees, 30 taxa, 3 reticulations

In this experiment, we simulated 50 networks each for 30 taxa and 3 reticulation. In addition to the NetRAX inference starting from the RAxML-NG best ML tree, we also started another NetRAX inference using 3 random and 3 maximum parsimony starting trees.

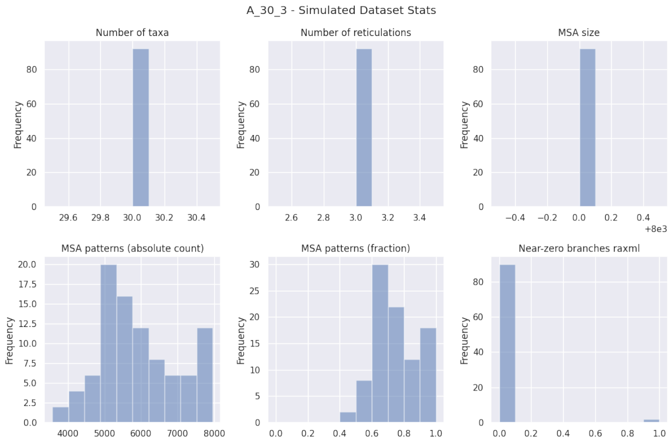


**Fig. 19.** Simulated dataset statistics for experiment A1, 30 taxa, 3 reticulations.

| A_30_3_norandom | LhType.AVERAGE | LhType.BEST |
|---|---|---|
| Inferred BIC better or equal | 2 (4.35 %) | 3 (6.52 %) |
| Inferred AIC better or equal | 0 (0.00 %) | 1 (2.17 %) |
| Inferred AICc better or equal | 0 (0.00 %) | 1 (2.17 %) |
| Inferred BIC worse | 44 (95.65 %) | 43 (93.48 %) |
| Inferred AIC worse | 46 (100.00 %) | 45 (97.83 %) |
| Inferred AICc worse | 46 (100.00 %) | 45 (97.83 %) |
| Inferred lnL better or equal | 0 (0.00 %) | 1 (2.17 %) |
| Inferred lnL worse | 46 (100.00 %) | 45 (97.83 %) |
| Inferred n_reticulations less | 9 (19.57 %) | 9 (19.57 %) |
| Inferred n_reticulations equal | 34 (73.91 %) | 37 (80.43 %) |
| Inferred n_reticulations more | 3 (6.52 %) | 0 (0.00 %) |
| Unrooted softwired distance zero | 14 (30.43 %) | 17 (36.96 %) |
| Good result | 16 (34.78 %) | 19 (41.30 %) |
| Passable result | 20 (43.48 %) | 20 (43.48 %) |
| Bad result | 10 (21.74 %) | 7 (15.22 %) |

| A_30_3_random | LhType.AVERAGE | LhType.BEST |
|---|---|---|
| Inferred BIC better or equal | 6 (13.04 %) | 5 (10.87 %) |
| Inferred AIC better or equal | 4 (8.70 %) | 3 (6.52 %) |
| Inferred AICc better or equal | 4 (8.70 %) | 3 (6.52 %) |
| Inferred BIC worse | 40 (86.96 %) | 41 (89.13 %) |
| Inferred AIC worse | 42 (91.30 %) | 43 (93.48 %) |
| Inferred AICc worse | 42 (91.30 %) | 43 (93.48 %) |
| Inferred lnL better or equal | 4 (8.70 %) | 3 (6.52 %) |
| Inferred lnL worse | 42 (91.30 %) | 43 (93.48 %) |
| Inferred n_reticulations less | 6 (13.04 %) | 5 (10.87 %) |
| Inferred n_reticulations equal | 39 (84.78 %) | 40 (86.96 %) |
| Inferred n_reticulations more | 1 (2.17 %) | 1 (2.17 %) |
| Unrooted softwired distance zero | 17 (36.96 %) | 18 (39.13 %) |
| Good result | 21 (45.65 %) | 21 (45.65 %) |
| Passable result | 20 (43.48 %) | 21 (45.65 %) |
| Bad result | 5 (10.87 %) | 4 (8.70 %) |

**Table 7.** Summary statistics for experiment A1, 39 taxa, 3 reticulations. Left: starting from RAxML-NG best tree, right: starting from 3 maximum parsimony and 3 random trees.

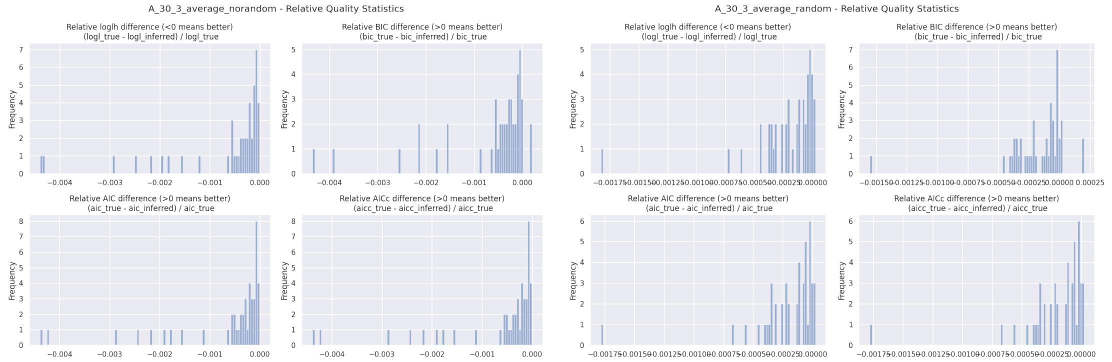


**Fig. 20.** Relative BIC, AIC, AICc, and loglikelihood differences for experiment A1, 30 taxa, 3 reticulations, `LhModel.AVERAGE`. Left: starting from RAxML-NG best tree, right: starting from 3 maximum parsimony and 3 random trees.

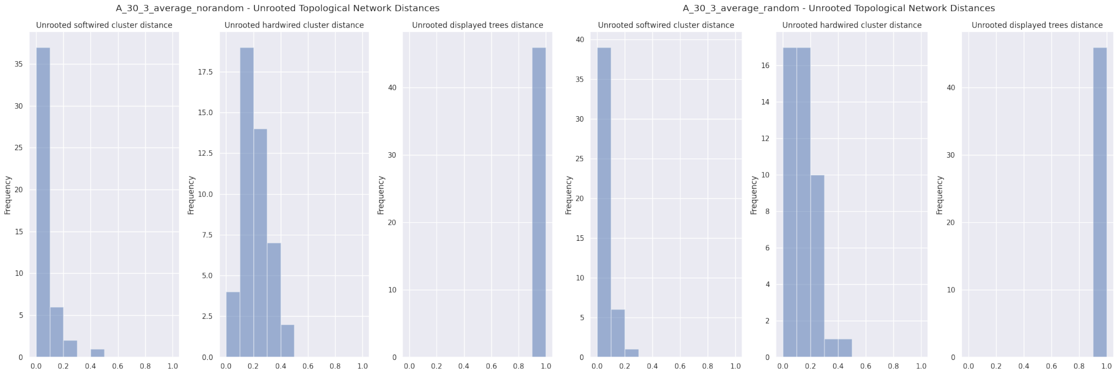


**Fig. 21.** Unrooted relative distances for experiment A1, 30 taxa, 3 reticulations, `LhModel.AVERAGE`. Left: starting from RAxML-NG best tree, right: starting from 3 maximum parsimony and 3 random trees.

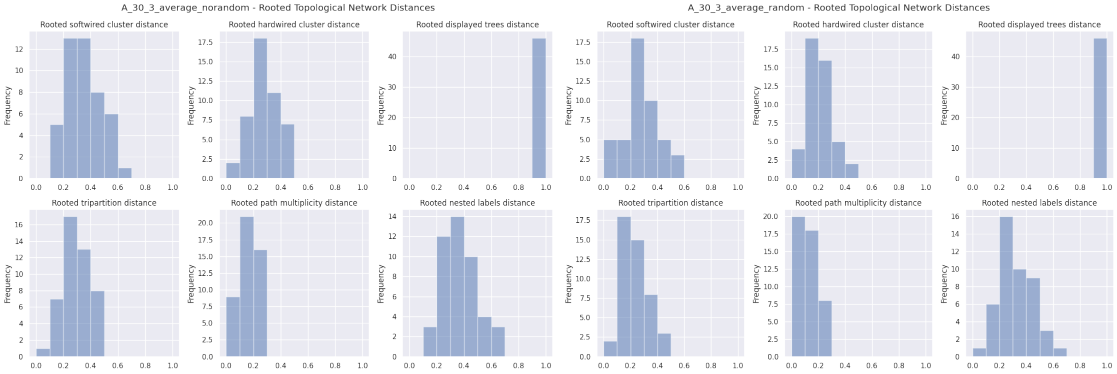


**Fig. 22.** Rooted relative distances for experiment A1, 30 taxa, 3 reticulations, `LhModel.AVERAGE`. Left: starting from RAxML-NG best tree, right: starting from 3 maximum parsimony and 3 random trees.

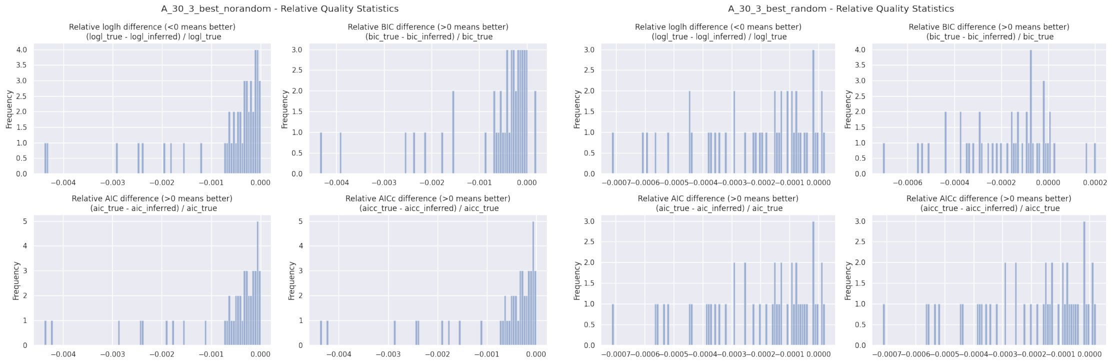


**Fig. 23.** Relative BIC, AIC, AICc, and loglikelihood differences for experiment A1, 30 taxa, 3 reticulations, `LhModel.BEST`. Left: starting from RAxML-NG best tree, right: starting from 3 maximum parsimony and 3 random trees.

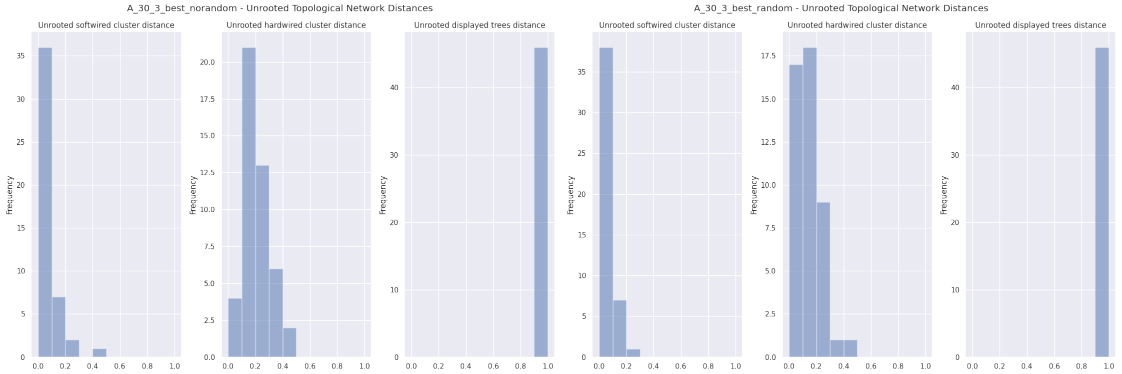


**Fig. 24.** Unrooted relative distances for experiment A1, 30 taxa, 3 reticulations, `LhModel.BEST`. Left: starting from RAxML-NG best tree, right: starting from 3 maximum parsimony and 3 random trees.

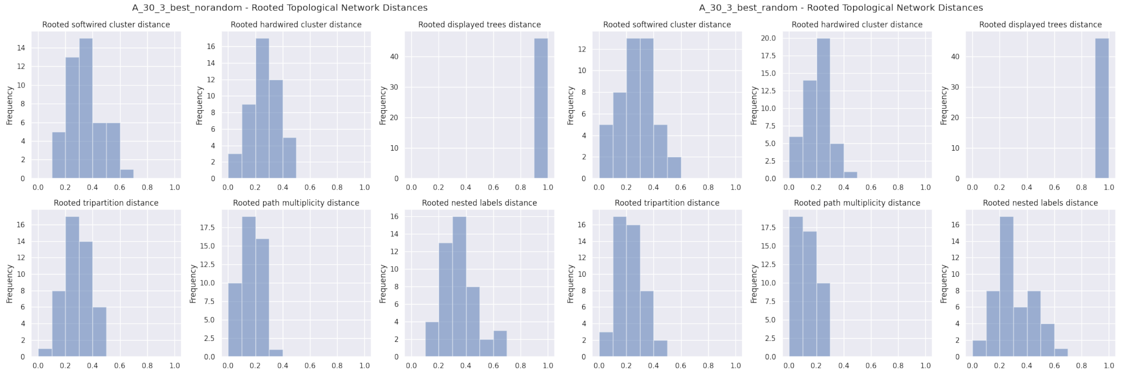


**Fig. 25.** Rooted relative distances for experiment A1, 30 taxa, 3 reticulations, `LhModel.BEST`. Left: starting from RAxML-NG best tree, right: starting from 3 maximum parsimony and 3 random trees.

| | count | mean | std | min | 10% | 25% | 50% | 75% | 90% | max |
|---|---|---|---|---|---|---|---|---|---|---|
| n_reticulations_inferred | 46.0 | 2.869565 | 0.499275 | 2.000000 | 2.000000 | 3.000000 | 3.000000 | 3.000000 | 3.000000 | 4.000000 |
| bic_diff | 46.0 | -108.899826 | 171.349289 | -777.532600 | -334.381100 | -89.461825 | -49.125350 | -16.360475 | -6.426600 | 31.452700 |
| bic_diff_relative | 46.0 | -0.000633 | 0.000982 | -0.004348 | -0.001945 | -0.000560 | -0.000280 | -0.000079 | -0.000033 | 0.000200 |
| aic_diff | 46.0 | -114.319867 | 179.539495 | -777.532600 | -355.157800 | -85.479025 | -45.772850 | -16.360450 | -9.799700 | -0.957800 |
| aic_diff_relative | 46.0 | -0.000673 | 0.001041 | -0.004384 | -0.002031 | -0.000540 | -0.000264 | -0.000091 | -0.000055 | -0.000006 |
| aicc_diff | 46.0 | -114.318650 | 179.537405 | -777.532500 | -355.153100 | -85.479025 | -45.777700 | -16.360525 | -9.799700 | -0.957800 |
| aicc_diff_relative | 46.0 | -0.000673 | 0.001041 | -0.004384 | -0.002031 | -0.000540 | -0.000264 | -0.000091 | -0.000055 | -0.000006 |
| lnL_diff | 46.0 | 57.681671 | 90.673216 | 0.478920 | 4.899850 | 9.074590 | 20.886430 | 42.739507 | 179.578920 | 388.766290 |
| lnL_diff_relative | 46.0 | -0.000680 | 0.001054 | -0.004391 | -0.002047 | -0.000559 | -0.000259 | -0.000101 | -0.000055 | -0.000006 |
| unrooted_softwired_network_distance | 46.0 | 0.068673 | 0.084258 | 0.000000 | 0.000000 | 0.000000 | 0.048897 | 0.091990 | 0.162791 | 0.444444 |
| unrooted_hardwired_network_distance | 46.0 | 0.205806 | 0.090940 | 0.034483 | 0.121212 | 0.151515 | 0.196691 | 0.257143 | 0.309921 | 0.461538 |
| unrooted_displayed_trees_distance | 46.0 | 1.000000 | 0.000000 | 1.000000 | 1.000000 | 1.000000 | 1.000000 | 1.000000 | 1.000000 | 1.000000 |
| rooted_softwired_network_distance | 46.0 | 0.354961 | 0.127619 | 0.137255 | 0.194057 | 0.262153 | 0.361630 | 0.451623 | 0.527001 | 0.636364 |
| rooted_hardwired_network_distance | 46.0 | 0.274168 | 0.104729 | 0.062500 | 0.138258 | 0.212121 | 0.267460 | 0.346847 | 0.428812 | 0.475000 |
| rooted_displayed_trees_distance | 46.0 | 1.000000 | 0.000000 | 1.000000 | 1.000000 | 1.000000 | 1.000000 | 1.000000 | 1.000000 | 1.000000 |
| rooted_tripartition_distance | 46.0 | 0.294405 | 0.105141 | 0.062500 | 0.176471 | 0.228571 | 0.277778 | 0.368421 | 0.450000 | 0.487805 |
| rooted_path_multiplicity_distance | 46.0 | 0.170868 | 0.065153 | 0.031746 | 0.092308 | 0.121212 | 0.176471 | 0.213768 | 0.253521 | 0.291667 |
| rooted_nested_labels_distance | 46.0 | 0.372184 | 0.121725 | 0.111111 | 0.236313 | 0.300000 | 0.380952 | 0.426080 | 0.510638 | 0.653061 |
| runtime_raxml | 46.0 | 57.612826 | 17.096124 | 30.138000 | 38.317000 | 42.942250 | 56.206500 | 70.436250 | 79.729500 | 100.538000 |
| runtime_inference | 46.0 | 1264.042000 | 747.795029 | 275.449000 | 523.873000 | 791.349500 | 1212.247000 | 1491.949250 | 1967.520500 | 4478.980000 |

| | count | mean | std | min | 10% | 25% | 50% | 75% | 90% | max |
|---|---|---|---|---|---|---|---|---|---|---|
| n_reticulations_inferred | 46.0 | 2.891304 | 0.378785 | 2.000000 | 2.000000 | 3.000000 | 3.000000 | 3.000000 | 3.000000 | 4.000000 |
| bic_diff | 46.0 | -28.438228 | 47.624974 | -299.285900 | -56.382300 | -46.034150 | -18.370250 | -5.736600 | 0.911100 | 32.317500 |
| bic_diff_relative | 46.0 | -0.000169 | 0.000260 | -0.001552 | -0.000369 | -0.000276 | -0.000096 | -0.000026 | 0.000005 | 0.000201 |
| aic_diff | 46.0 | -32.954937 | 52.803962 | -340.839500 | -60.739950 | -48.283800 | -18.370200 | -7.346975 | -1.079600 | 3.440200 |
| aic_diff_relative | 46.0 | -0.000201 | 0.000290 | -0.001781 | -0.000390 | -0.000300 | -0.000112 | -0.000030 | -0.000008 | 0.000022 |
| aicc_diff | 46.0 | -32.953909 | 52.802303 | -340.830000 | -60.739950 | -48.283725 | -18.370200 | -7.346975 | -1.079650 | 3.440200 |
| aicc_diff_relative | 46.0 | -0.000201 | 0.000290 | -0.001781 | -0.000390 | -0.000300 | -0.000112 | -0.000030 | -0.000008 | 0.000022 |
| lnL_diff | 46.0 | 16.912244 | 27.135175 | -1.720090 | 0.539805 | 3.673492 | 9.431220 | 24.141872 | 31.153100 | 174.419720 |
| lnL_diff_relative | 46.0 | -0.000207 | 0.000300 | -0.001825 | -0.000419 | -0.000301 | -0.000108 | -0.000030 | -0.000008 | 0.000022 |
| unrooted_softwired_network_distance | 46.0 | 0.047036 | 0.056707 | 0.000000 | 0.000000 | 0.000000 | 0.026084 | 0.068770 | 0.127144 | 0.243243 |
| unrooted_hardwired_network_distance | 46.0 | 0.153599 | 0.088535 | 0.000000 | 0.065591 | 0.096774 | 0.125000 | 0.201287 | 0.275210 | 0.421053 |
| unrooted_displayed_trees_distance | 46.0 | 1.000000 | 0.000000 | 1.000000 | 1.000000 | 1.000000 | 1.000000 | 1.000000 | 1.000000 | 1.000000 |
| rooted_softwired_network_distance | 46.0 | 0.281389 | 0.133146 | 0.000000 | 0.105455 | 0.205622 | 0.267669 | 0.376425 | 0.464953 | 0.562500 |
| rooted_hardwired_network_distance | 46.0 | 0.208401 | 0.092858 | 0.000000 | 0.121212 | 0.125000 | 0.191176 | 0.274510 | 0.333333 | 0.421053 |
| rooted_displayed_trees_distance | 46.0 | 1.000000 | 0.000000 | 1.000000 | 1.000000 | 1.000000 | 1.000000 | 1.000000 | 1.000000 | 1.000000 |
| rooted_tripartition_distance | 46.0 | 0.226450 | 0.098934 | 0.000000 | 0.121212 | 0.151515 | 0.205882 | 0.285714 | 0.351351 | 0.450000 |
| rooted_path_multiplicity_distance | 46.0 | 0.131187 | 0.063909 | 0.000000 | 0.062500 | 0.092308 | 0.121212 | 0.176471 | 0.222878 | 0.267606 |
| rooted_nested_labels_distance | 46.0 | 0.318545 | 0.133841 | 0.000000 | 0.162162 | 0.210526 | 0.320732 | 0.418605 | 0.466403 | 0.680000 |
| runtime_inference | 46.0 | 9259.401630 | 3802.198244 | 3065.790000 | 4965.726500 | 6874.954250 | 8458.074000 | 11196.525750 | 13492.716000 | 23662.042000 |

**Table 8.** Percentiles for experiment A1, 30 taxa, 3 reticulations, `LhModel.AVERAGE`. Top: starting from RAxML-NG best tree, bottom: starting from 3 maximum parsimony and 3 random trees.

| | count | mean | std | min | 10% | 25% | 50% | 75% | 90% | max |
|---|---|---|---|---|---|---|---|---|---|---|
| n_reticulations_inferred | 46.0 | 2.804348 | 0.401085 | 2.000000 | 2.000000 | 3.000000 | 3.000000 | 3.000000 | 3.000000 | 3.000000 |
| bic_diff | 46.0 | -115.396902 | 171.113977 | -777.532600 | -336.725600 | -92.715550 | -53.045500 | -21.023725 | -7.147200 | 31.746800 |
| bic_diff_relative | 46.0 | -0.000669 | 0.000980 | -0.004348 | -0.001945 | -0.000619 | -0.000316 | -0.000131 | -0.000033 | 0.000197 |
| aic_diff | 46.0 | -123.526952 | 178.063258 | -777.532600 | -373.797950 | -92.715550 | -54.797200 | -21.023675 | -11.249600 | 0.314600 |
| aic_diff_relative | 46.0 | -0.000727 | 0.001031 | -0.004384 | -0.002144 | -0.000626 | -0.000329 | -0.000147 | -0.000059 | 0.000002 |
| aicc_diff | 46.0 | -123.525113 | 178.061528 | -777.532500 | -373.793250 | -92.715600 | -54.797200 | -21.023675 | -11.249600 | 0.314600 |
| aicc_diff_relative | 46.0 | -0.000727 | 0.001031 | -0.004384 | -0.002144 | -0.000626 | -0.000329 | -0.000147 | -0.000059 | 0.000002 |
| lnL_diff | 46.0 | 62.546084 | 89.773959 | -0.157280 | 6.338185 | 11.094985 | 27.398605 | 46.357785 | 188.898985 | 388.766290 |
| lnL_diff_relative | 46.0 | -0.000738 | 0.001042 | -0.004391 | -0.002160 | -0.000627 | -0.000329 | -0.000149 | -0.000067 | 0.000002 |
| unrooted_softwired_network_distance | 46.0 | 0.067717 | 0.085834 | 0.000000 | 0.000000 | 0.000000 | 0.048897 | 0.094684 | 0.162791 | 0.444444 |
| unrooted_hardwired_network_distance | 46.0 | 0.200775 | 0.090094 | 0.062500 | 0.110606 | 0.151515 | 0.179144 | 0.251681 | 0.309921 | 0.461538 |
| unrooted_displayed_trees_distance | 46.0 | 1.000000 | 0.000000 | 1.000000 | 1.000000 | 1.000000 | 1.000000 | 1.000000 | 1.000000 | 1.000000 |
| rooted_softwired_network_distance | 46.0 | 0.349875 | 0.130077 | 0.100000 | 0.191303 | 0.237695 | 0.370485 | 0.422847 | 0.527001 | 0.636364 |
| rooted_hardwired_network_distance | 46.0 | 0.268010 | 0.105630 | 0.062500 | 0.123106 | 0.189394 | 0.271242 | 0.328571 | 0.422368 | 0.475000 |
| rooted_displayed_trees_distance | 46.0 | 1.000000 | 0.000000 | 1.000000 | 1.000000 | 1.000000 | 1.000000 | 1.000000 | 1.000000 | 1.000000 |
| rooted_tripartition_distance | 46.0 | 0.283885 | 0.102465 | 0.062500 | 0.163993 | 0.228571 | 0.277778 | 0.368421 | 0.430128 | 0.475000 |
| rooted_path_multiplicity_distance | 46.0 | 0.164854 | 0.064969 | 0.031746 | 0.085216 | 0.121212 | 0.170325 | 0.202899 | 0.241046 | 0.301370 |
| rooted_nested_labels_distance | 46.0 | 0.353808 | 0.126303 | 0.111111 | 0.210526 | 0.256410 | 0.365476 | 0.418605 | 0.494444 | 0.653061 |
| runtime_raxml | 46.0 | 57.612826 | 17.096124 | 30.138000 | 38.317000 | 42.942250 | 56.206500 | 70.436250 | 79.729500 | 100.538000 |
| runtime_inference | 46.0 | 1083.018087 | 646.928006 | 263.758000 | 480.736500 | 730.942500 | 966.784500 | 1280.208500 | 1541.123000 | 4218.982000 |

| | count | mean | std | min | 10% | 25% | 50% | 75% | 90% | max |
|---|---|---|---|---|---|---|---|---|---|---|
| n_reticulations_inferred | 46.0 | 2.913043 | 0.354406 | 2.000000 | 2.500000 | 3.000000 | 3.000000 | 3.000000 | 3.000000 | 4.000000 |
| bic_diff | 46.0 | -29.385015 | 30.084049 | -94.035500 | -76.918350 | -48.656325 | -20.275550 | -10.126675 | 0.216400 | 32.611600 |
| bic_diff_relative | 46.0 | -0.000179 | 0.000190 | -0.000706 | -0.000437 | -0.000291 | -0.000131 | -0.000054 | 0.000001 | 0.000203 |
| aic_diff | 46.0 | -32.998367 | 28.692402 | -94.035600 | -80.474650 | -51.248575 | -21.964650 | -10.744400 | -3.267650 | 3.440200 |
| aic_diff_relative | 46.0 | -0.000206 | 0.000185 | -0.000714 | -0.000480 | -0.000336 | -0.000150 | -0.000067 | -0.000015 | 0.000022 |
| aicc_diff | 46.0 | -32.997557 | 28.691870 | -94.035600 | -80.470000 | -51.241425 | -21.964700 | -10.744325 | -3.267700 | 3.440200 |
| aicc_diff_relative | 46.0 | -0.000206 | 0.000185 | -0.000714 | -0.000480 | -0.000336 | -0.000150 | -0.000067 | -0.000015 | 0.000022 |
| lnL_diff | 46.0 | 16.847012 | 14.639829 | -1.720090 | 1.633875 | 5.709847 | 12.097380 | 26.047238 | 42.026225 | 47.017780 |
| lnL_diff_relative | 46.0 | -0.000211 | 0.000190 | -0.000716 | -0.000483 | -0.000336 | -0.000150 | -0.000073 | -0.000015 | 0.000022 |
| unrooted_softwired_network_distance | 46.0 | 0.047388 | 0.057814 | 0.000000 | 0.000000 | 0.000000 | 0.024390 | 0.068770 | 0.139212 | 0.243243 |
| unrooted_hardwired_network_distance | 46.0 | 0.150973 | 0.085691 | 0.000000 | 0.064516 | 0.096774 | 0.138258 | 0.187500 | 0.246218 | 0.421053 |
| unrooted_displayed_trees_distance | 46.0 | 1.000000 | 0.000000 | 1.000000 | 1.000000 | 1.000000 | 1.000000 | 1.000000 | 1.000000 | 1.000000 |
| rooted_softwired_network_distance | 46.0 | 0.272687 | 0.127051 | 0.000000 | 0.099026 | 0.193250 | 0.267492 | 0.354041 | 0.423389 | 0.562500 |
| rooted_hardwired_network_distance | 46.0 | 0.201937 | 0.092985 | 0.000000 | 0.093750 | 0.122159 | 0.205882 | 0.264706 | 0.333333 | 0.421053 |
| rooted_displayed_trees_distance | 46.0 | 1.000000 | 0.000000 | 1.000000 | 1.000000 | 1.000000 | 1.000000 | 1.000000 | 1.000000 | 1.000000 |
| rooted_tripartition_distance | 46.0 | 0.219215 | 0.098803 | 0.000000 | 0.121212 | 0.128788 | 0.217227 | 0.283730 | 0.351351 | 0.435897 |
| rooted_path_multiplicity_distance | 46.0 | 0.129339 | 0.068435 | 0.000000 | 0.062500 | 0.069952 | 0.121212 | 0.176471 | 0.228571 | 0.291667 |
| rooted_nested_labels_distance | 46.0 | 0.306013 | 0.149560 | 0.000000 | 0.136637 | 0.210526 | 0.278205 | 0.418605 | 0.489130 | 0.680000 |
| runtime_inference | 46.0 | 10300.961261 | 9669.241115 | 3632.946000 | 5132.770000 | 6600.156250 | 8340.384500 | 10500.451750 | 14526.196000 | 68019.888000 |

**Table 9.** Percentiles for experiment A1, 30 taxa, 3 reticulations, `LhModel.BEST`. Top: starting from RAxML-NG best tree, bottom: starting from 3 maximum parsimony and 3 random trees.

##### 6.4 A2: 40 taxa, 1 reticulation

In this experiment, we simulated 50 networks each for 40 taxa and 1 reticulation. We ran NetRAX inference starting from the RAxML-NG best ML tree.

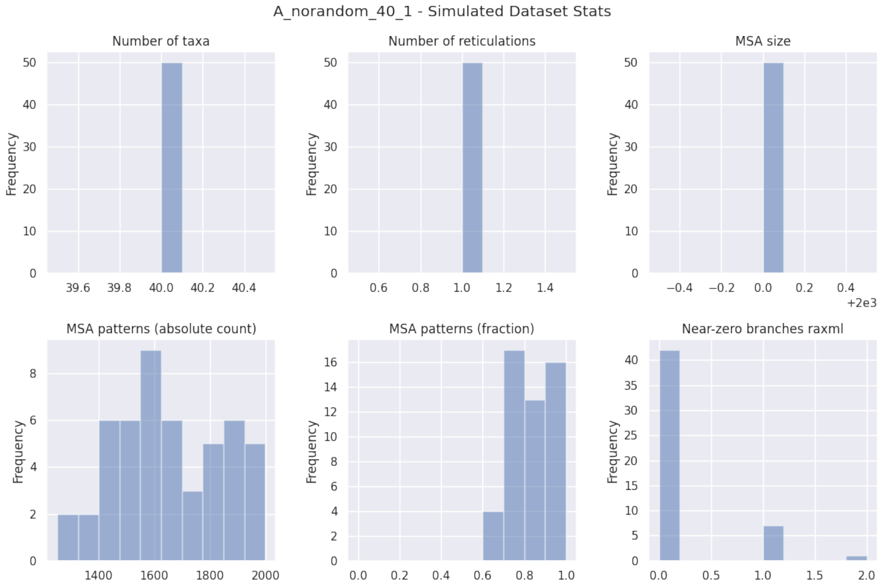


**Fig. 26.** Simulated dataset statistics for experiment A2, 40 taxa, 1 reticulation.

| A_norandom_40_1_norandom | LhType.AVERAGE | LhType.BEST |
|---|---|---|
| Inferred BIC better or equal | 23 (46.00 %) | 21 (42.00 %) |
| Inferred AIC better or equal | 21 (42.00 %) | 19 (38.00 %) |
| Inferred AICc better or equal | 21 (42.00 %) | 19 (38.00 %) |
| Inferred BIC worse | 27 (54.00 %) | 29 (58.00 %) |
| Inferred AIC worse | 29 (58.00 %) | 31 (62.00 %) |
| Inferred AICc worse | 29 (58.00 %) | 31 (62.00 %) |
| Inferred lnL better or equal | 21 (42.00 %) | 19 (38.00 %) |
| Inferred lnL worse | 29 (58.00 %) | 31 (62.00 %) |
| Inferred n_reticulations less | 2 (4.00 %) | 2 (4.00 %) |
| Inferred n_reticulations equal | 48 (96.00 %) | 48 (96.00 %) |
| Inferred n_reticulations more | 0 (0.00 %) | 0 (0.00 %) |
| Unrooted softwired distance zero | 13 (26.00 %) | 13 (26.00 %) |
| Good result | 34 (68.00 %) | 33 (66.00 %) |
| Passable result | 16 (32.00 %) | 17 (34.00 %) |
| Bad result | 0 (0.00 %) | 0 (0.00 %) |

**Table 10.** Summary statistics for experiment A1, 40 taxa, 1 reticulation, starting from RAxML-NG best tree.

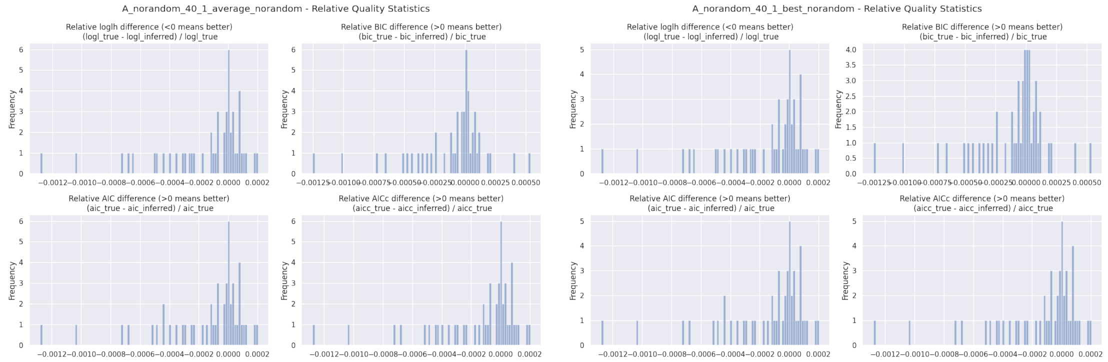


**Fig. 27.** Relative BIC, AIC, AICc, and loglikelihood differences for experiment A2, 40 taxa, 1 reticulation, starting from RAxML-NG best tree. Left: `LhType.AVERAGE`, right: `LhType.BEST`.

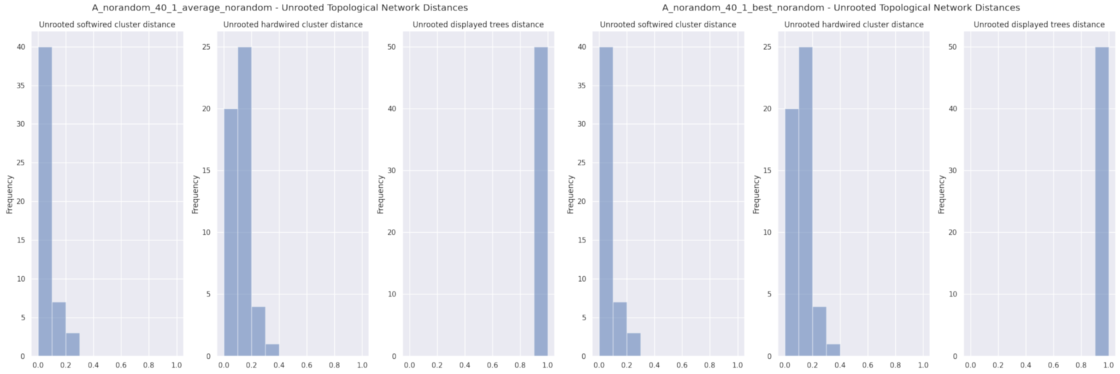


**Fig. 28.** Unrooted relative distances for experiment A2, 40 taxa, 1 reticulation, starting from RAxML-NG best tree. Left: `LhType.AVERAGE`, right: `LhType.BEST`.

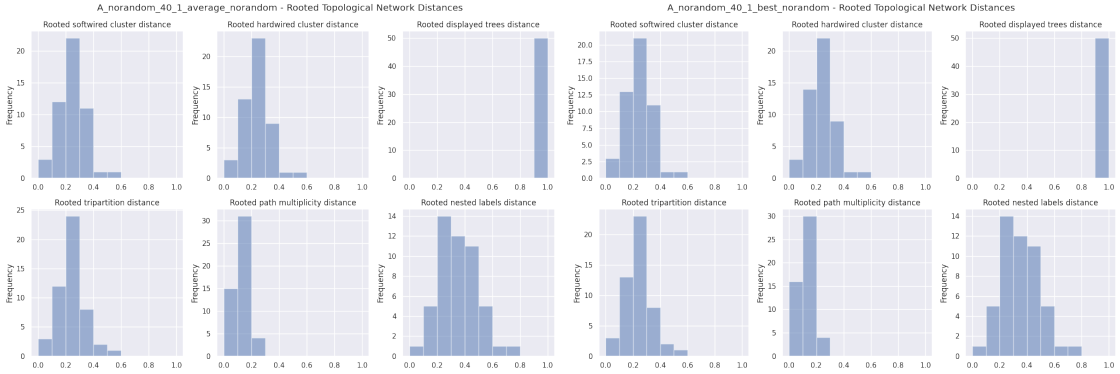


**Fig. 29.** Rooted relative distances for experiment A2, 40 taxa, 1 reticulation, starting from RAxML-NG best tree. Left: `LhType.AVERAGE`, right: `LhType.BEST`.

| | count | mean | std | min | 10% | 25% | 50% | 75% | 90% | max |
|---|---|---|---|---|---|---|---|---|---|---|
| n_reticulations_inferred | 50.0 | 0.960000 | 0.197949 | 0.000000 | 1.000000 | 1.000000 | 1.000000 | 1.000000 | 1.000000 | 1.000000 |
| bic_diff | 50.0 | -5.724288 | 16.059622 | -58.566990 | -22.014517 | -9.743140 | -0.514435 | 1.986412 | 5.593636 | 27.506480 |
| bic_diff_relative | 50.0 | -0.000111 | 0.000314 | -0.001253 | -0.000479 | -0.000221 | -0.000010 | 0.000042 | 0.000110 | 0.000538 |
| aic_diff | 50.0 | -7.210654 | 14.880970 | -58.566990 | -22.014517 | -11.586745 | -0.888630 | 1.463027 | 4.007042 | 9.341350 |
| aic_diff_relative | 50.0 | -0.000144 | 0.000299 | -0.001279 | -0.000489 | -0.000245 | -0.000021 | 0.000029 | 0.000083 | 0.000203 |
| aicc_diff | 50.0 | -7.209864 | 14.880655 | -58.566990 | -22.014507 | -11.586745 | -0.888630 | 1.463035 | 4.007042 | 9.341340 |
| aicc_diff_relative | 50.0 | -0.000144 | 0.000299 | -0.001279 | -0.000489 | -0.000245 | -0.000021 | 0.000029 | 0.000083 | 0.000203 |
| lnL_diff | 50.0 | 3.765326 | 7.545662 | -4.670670 | -2.003521 | -0.731520 | 0.444315 | 6.819280 | 12.597144 | 29.283500 |
| lnL_diff_relative | 50.0 | -0.000151 | 0.000306 | -0.001284 | -0.000524 | -0.000256 | -0.000021 | 0.000029 | 0.000083 | 0.000203 |
| unrooted_softwired_network_distance | 50.0 | 0.066663 | 0.063218 | 0.000000 | 0.000000 | 0.005952 | 0.048780 | 0.093023 | 0.142248 | 0.260000 |
| unrooted_hardwired_network_distance | 50.0 | 0.109468 | 0.070759 | 0.000000 | 0.023077 | 0.051282 | 0.100000 | 0.166667 | 0.196540 | 0.311111 |
| unrooted_displayed_trees_distance | 50.0 | 1.000000 | 0.000000 | 1.000000 | 1.000000 | 1.000000 | 1.000000 | 1.000000 | 1.000000 | 1.000000 |
| rooted_softwired_network_distance | 50.0 | 0.251583 | 0.090266 | 0.062500 | 0.153953 | 0.186410 | 0.264615 | 0.299691 | 0.353817 | 0.564516 |
| rooted_hardwired_network_distance | 50.0 | 0.236054 | 0.084804 | 0.050000 | 0.140767 | 0.186047 | 0.227273 | 0.266667 | 0.326087 | 0.519231 |
| rooted_displayed_trees_distance | 50.0 | 1.000000 | 0.000000 | 1.000000 | 1.000000 | 1.000000 | 1.000000 | 1.000000 | 1.000000 | 1.000000 |
| rooted_tripartition_distance | 50.0 | 0.241981 | 0.085600 | 0.050000 | 0.142857 | 0.186047 | 0.227273 | 0.266667 | 0.340426 | 0.519231 |
| rooted_path_multiplicity_distance | 50.0 | 0.129596 | 0.049015 | 0.024691 | 0.072289 | 0.095238 | 0.123529 | 0.155573 | 0.181818 | 0.297872 |
| rooted_nested_labels_distance | 50.0 | 0.348541 | 0.141230 | 0.095238 | 0.181818 | 0.222222 | 0.333333 | 0.431373 | 0.521212 | 0.730159 |
| runtime_raxml | 50.0 | 29.700360 | 4.005916 | 20.857000 | 24.137200 | 26.426250 | 30.535000 | 32.407500 | 34.081600 | 39.654000 |
| runtime_inference | 50.0 | 61.959200 | 17.524851 | 7.762000 | 47.098500 | 53.656000 | 61.351000 | 69.426500 | 84.238300 | 100.261000 |

| | count | mean | std | min | 10% | 25% | 50% | 75% | 90% | max |
|---|---|---|---|---|---|---|---|---|---|---|
| n_reticulations_inferred | 50.0 | 0.960000 | 0.197949 | 0.000000 | 1.000000 | 1.000000 | 1.000000 | 1.000000 | 1.000000 | 1.000000 |
| bic_diff | 50.0 | -5.809648 | 16.037390 | -58.566990 | -22.014517 | -9.743140 | -0.693200 | 1.986412 | 5.593636 | 27.506480 |
| bic_diff_relative | 50.0 | -0.000112 | 0.000314 | -0.001253 | -0.000479 | -0.000221 | -0.000015 | 0.000042 | 0.000110 | 0.000538 |
| aic_diff | 50.0 | -7.296015 | 14.848257 | -58.566990 | -22.014517 | -11.586745 | -1.542485 | 1.463018 | 4.007042 | 9.341350 |
| aic_diff_relative | 50.0 | -0.000145 | 0.000299 | -0.001279 | -0.000489 | -0.000245 | -0.000026 | 0.000029 | 0.000083 | 0.000203 |
| aicc_diff | 50.0 | -7.295225 | 14.847946 | -58.566990 | -22.014507 | -11.586745 | -1.542485 | 1.463025 | 4.007042 | 9.341340 |
| aicc_diff_relative | 50.0 | -0.000145 | 0.000299 | -0.001279 | -0.000489 | -0.000245 | -0.000026 | 0.000029 | 0.000083 | 0.000203 |
| lnL_diff | 50.0 | 3.808006 | 7.528609 | -4.670670 | -2.003521 | -0.731518 | 0.771240 | 6.819280 | 12.597144 | 29.283500 |
| lnL_diff_relative | 50.0 | -0.000152 | 0.000306 | -0.001284 | -0.000524 | -0.000256 | -0.000026 | 0.000029 | 0.000083 | 0.000203 |
| unrooted_softwired_network_distance | 50.0 | 0.066663 | 0.063218 | 0.000000 | 0.000000 | 0.005952 | 0.048780 | 0.093023 | 0.142248 | 0.260000 |
| unrooted_hardwired_network_distance | 50.0 | 0.109468 | 0.070759 | 0.000000 | 0.023077 | 0.051282 | 0.100000 | 0.166667 | 0.196540 | 0.311111 |
| unrooted_displayed_trees_distance | 50.0 | 1.000000 | 0.000000 | 1.000000 | 1.000000 | 1.000000 | 1.000000 | 1.000000 | 1.000000 | 1.000000 |
| rooted_softwired_network_distance | 50.0 | 0.250889 | 0.090562 | 0.062500 | 0.153953 | 0.186410 | 0.264615 | 0.299691 | 0.353817 | 0.564516 |
| rooted_hardwired_network_distance | 50.0 | 0.235229 | 0.085091 | 0.050000 | 0.140767 | 0.186047 | 0.227273 | 0.266667 | 0.326087 | 0.519231 |
| rooted_displayed_trees_distance | 50.0 | 1.000000 | 0.000000 | 1.000000 | 1.000000 | 1.000000 | 1.000000 | 1.000000 | 1.000000 | 1.000000 |
| rooted_tripartition_distance | 50.0 | 0.241157 | 0.085942 | 0.050000 | 0.142857 | 0.186047 | 0.227273 | 0.266667 | 0.340426 | 0.519231 |
| rooted_path_multiplicity_distance | 50.0 | 0.129148 | 0.049229 | 0.024691 | 0.072289 | 0.095238 | 0.123529 | 0.155573 | 0.181818 | 0.297872 |
| rooted_nested_labels_distance | 50.0 | 0.347311 | 0.140491 | 0.095238 | 0.181818 | 0.222222 | 0.333333 | 0.431373 | 0.521212 | 0.730159 |
| runtime_raxml | 50.0 | 29.700360 | 4.005916 | 20.857000 | 24.137200 | 26.426250 | 30.535000 | 32.407500 | 34.081600 | 39.654000 |
| runtime_inference | 50.0 | 56.187740 | 19.695786 | 6.416000 | 40.013900 | 47.885250 | 53.911500 | 61.748750 | 73.931100 | 124.420000 |

**Table 11.** Percentiles for experiment A1, 40 taxa, 1 reticulation, `LhModel.AVERAGE`, starting from RAxML-NG best tree. Top: `LhType.AVERAGE`, bottom: `LhType.BEST`.

##### 6.5 A2: 40 taxa, 2 reticulations

In this experiment, we simulated 50 networks each for 40 taxa and 2 reticulations. We ran NetRAX inference starting from the RAxML-NG best ML tree.

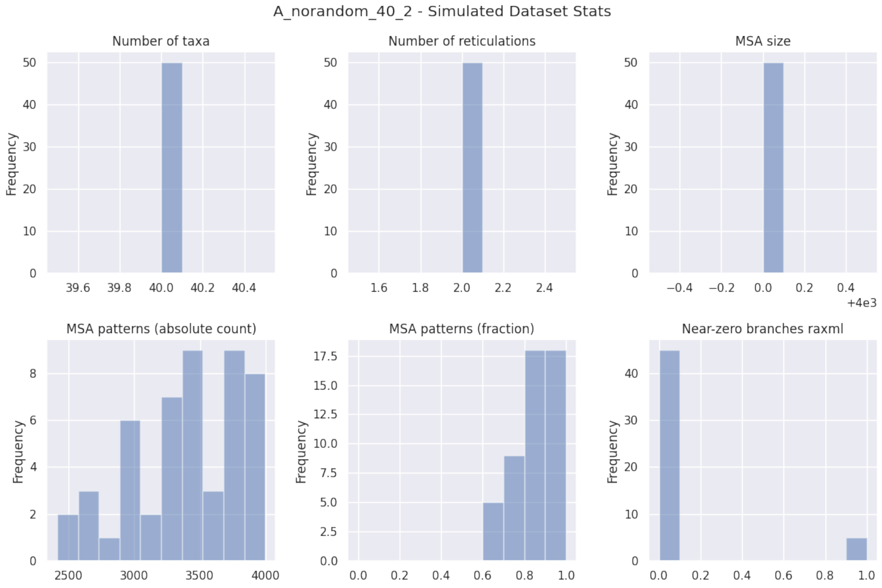


**Fig. 30.** Simulated dataset statistics for experiment A2, 40 taxa, 2 reticulations.

| A_norandom_40_2_norandom | LhType.AVERAGE | LhType.BEST |
|---|---|---|
| Inferred BIC better or equal | 3 (6.00 %) | 1 (2.00 %) |
| Inferred AIC better or equal | 2 (4.00 %) | 0 (0.00 %) |
| Inferred AICc better or equal | 2 (4.00 %) | 0 (0.00 %) |
| Inferred BIC worse | 47 (94.00 %) | 49 (98.00 %) |
| Inferred AIC worse | 48 (96.00 %) | 50 (100.00 %) |
| Inferred AICc worse | 48 (96.00 %) | 50 (100.00 %) |
| Inferred lnL better or equal | 2 (4.00 %) | 0 (0.00 %) |
| Inferred lnL worse | 48 (96.00 %) | 50 (100.00 %) |
| Inferred n_reticulations less | 6 (12.00 %) | 6 (12.00 %) |
| Inferred n_reticulations equal | 43 (86.00 %) | 43 (86.00 %) |
| Inferred n_reticulations more | 1 (2.00 %) | 1 (2.00 %) |
| Unrooted softwired distance zero | 18 (36.00 %) | 18 (36.00 %) |
| Good result | 21 (42.00 %) | 19 (38.00 %) |
| Passable result | 23 (46.00 %) | 25 (50.00 %) |
| Bad result | 6 (12.00 %) | 6 (12.00 %) |

**Table 12.** Summary statistics for experiment A1, 40 taxa, 2 reticulations, starting from RAxML-NG best tree.

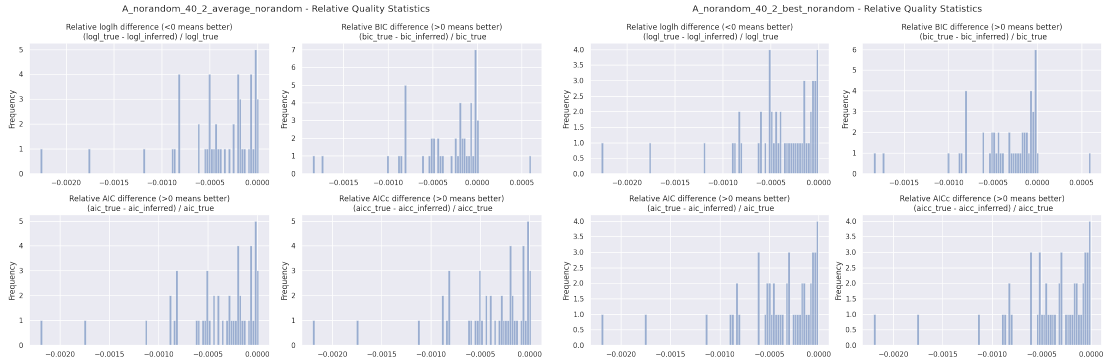


**Fig. 31.** Relative BIC, AIC, AICc, and loglikelihood differences for experiment A2, 40 taxa, 2 reticulations, starting from RAxML-NG best tree. Left: `LhType.AVERAGE`, right: `LhType.BEST`.

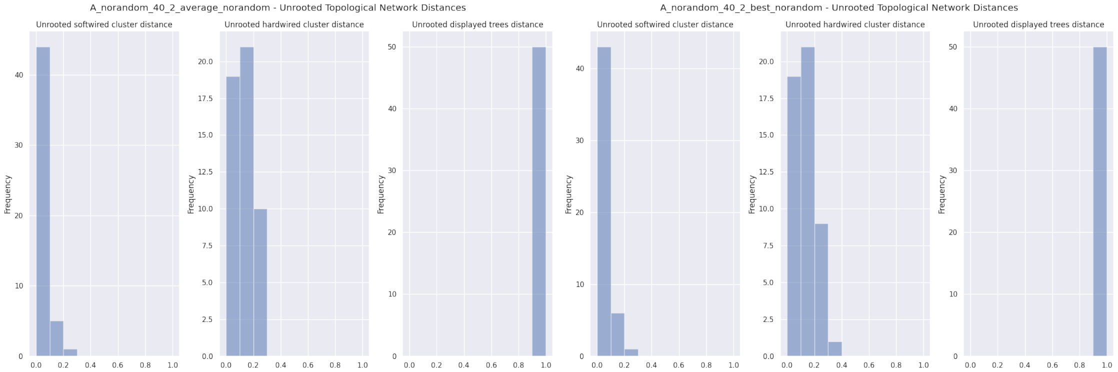


**Fig. 32.** Unrooted relative distances for experiment A2, 40 taxa, 2 reticulations, starting from RAxML-NG best tree. Left: `LhType.AVERAGE`, right: `LhType.BEST`.

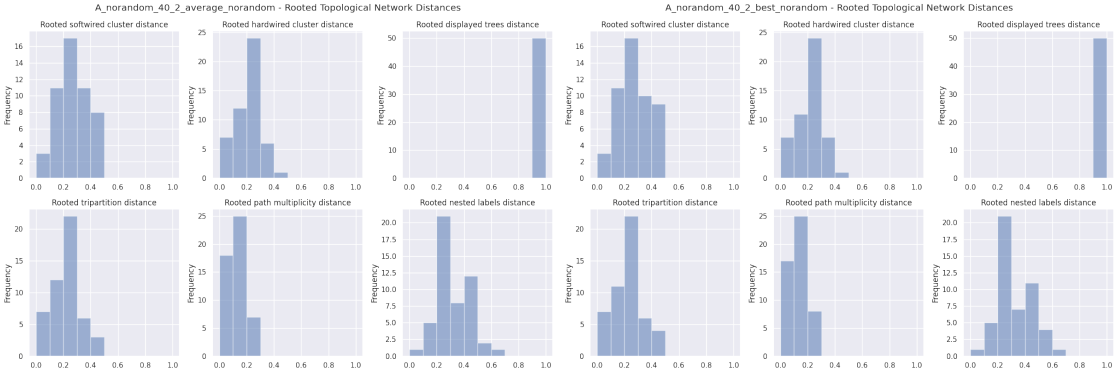


**Fig. 33.** Rooted relative distances for experiment A2, 40 taxa, 2 reticulations, starting from RAxML-NG best tree. Left: `LhType.AVERAGE`, right: `LhType.BEST`.

| | count | mean | std | min | 10% | 25% | 50% | 75% | 90% | max |
|---|---|---|---|---|---|---|---|---|---|---|
| n_reticulations_inferred | 50.0 | 1.880000 | 0.435187 | 0.000000 | 1.000000 | 2.000000 | 2.000000 | 2.000000 | 2.000000 | 3.000000 |
| bic_diff | 50.0 | -37.709614 | 48.502210 | -222.277300 | -77.526266 | -53.553940 | -25.309150 | -5.598025 | -1.143936 | 52.535530 |
| bic_diff_relative | 50.0 | -0.000353 | 0.000434 | -0.001846 | -0.000821 | -0.000504 | -0.000197 | -0.000061 | -0.000012 | 0.000602 |
| aic_diff | 50.0 | -42.501420 | 50.271065 | -262.209000 | -71.519624 | -53.553940 | -28.041510 | -11.288600 | -1.500384 | 1.057930 |
| aic_diff_relative | 50.0 | -0.000400 | 0.000436 | -0.002200 | -0.000833 | -0.000525 | -0.000277 | -0.000079 | -0.000014 | 0.000013 |
| aicc_diff | 50.0 | -42.500000 | 50.269637 | -262.197000 | -71.519624 | -53.553940 | -28.029645 | -11.288700 | -1.500384 | 1.057920 |
| aicc_diff_relative | 50.0 | -0.000400 | 0.000436 | -0.002200 | -0.000833 | -0.000525 | -0.000277 | -0.000079 | -0.000014 | 0.000013 |
| lnL_diff | 50.0 | 21.730711 | 25.665176 | -0.528960 | 0.750182 | 5.644330 | 15.402685 | 26.846483 | 35.759821 | 135.104510 |
| lnL_diff_relative | 50.0 | -0.000410 | 0.000444 | -0.002271 | -0.000836 | -0.000520 | -0.000277 | -0.000079 | -0.000014 | 0.000013 |
| unrooted_softwired_network_distance | 50.0 | 0.043439 | 0.049751 | 0.000000 | 0.000000 | 0.000000 | 0.039608 | 0.053730 | 0.111905 | 0.220000 |
| unrooted_hardwired_network_distance | 50.0 | 0.128095 | 0.068762 | 0.000000 | 0.050000 | 0.073171 | 0.119048 | 0.189369 | 0.222727 | 0.266667 |
| unrooted_displayed_trees_distance | 50.0 | 1.000000 | 0.000000 | 1.000000 | 1.000000 | 1.000000 | 1.000000 | 1.000000 | 1.000000 | 1.000000 |
| rooted_softwired_network_distance | 50.0 | 0.269006 | 0.115464 | 0.058824 | 0.122917 | 0.190617 | 0.254630 | 0.361768 | 0.447400 | 0.485714 |
| rooted_hardwired_network_distance | 50.0 | 0.217970 | 0.082133 | 0.048780 | 0.095238 | 0.181818 | 0.222222 | 0.260870 | 0.326812 | 0.408163 |
| rooted_displayed_trees_distance | 50.0 | 1.000000 | 0.000000 | 1.000000 | 1.000000 | 1.000000 | 1.000000 | 1.000000 | 1.000000 | 1.000000 |
| rooted_tripartition_distance | 50.0 | 0.229516 | 0.090206 | 0.048780 | 0.095238 | 0.181818 | 0.222222 | 0.282609 | 0.354167 | 0.431373 |
| rooted_path_multiplicity_distance | 50.0 | 0.127845 | 0.055200 | 0.024390 | 0.048193 | 0.094118 | 0.122093 | 0.159091 | 0.211111 | 0.247312 |
| rooted_nested_labels_distance | 50.0 | 0.324620 | 0.118991 | 0.090909 | 0.169855 | 0.250000 | 0.285714 | 0.415094 | 0.472727 | 0.610169 |
| runtime_raxml | 50.0 | 49.038660 | 9.746339 | 29.761000 | 37.481300 | 42.748000 | 47.852500 | 55.477750 | 58.564600 | 74.907000 |
| runtime_inference | 50.0 | 406.090140 | 333.840487 | 15.712000 | 154.637400 | 271.284500 | 351.776000 | 468.459750 | 549.047500 | 2414.109000 |

| | count | mean | std | min | 10% | 25% | 50% | 75% | 90% | max |
|---|---|---|---|---|---|---|---|---|---|---|
| n_reticulations_inferred | 50.0 | 1.880000 | 0.435187 | 0.000000 | 1.000000 | 2.000000 | 2.000000 | 2.000000 | 2.000000 | 3.000000 |
| bic_diff | 50.0 | -39.339389 | 48.237157 | -222.277300 | -77.749176 | -55.689435 | -26.130650 | -7.340350 | -2.932601 | 52.555280 |
| bic_diff_relative | 50.0 | -0.000363 | 0.000426 | -0.001846 | -0.000821 | -0.000504 | -0.000245 | -0.000064 | -0.000028 | 0.000603 |
| aic_diff | 50.0 | -44.131193 | 49.855173 | -262.209000 | -71.519624 | -55.689435 | -30.905160 | -13.263445 | -3.806742 | -0.373260 |
| aic_diff_relative | 50.0 | -0.000410 | 0.000426 | -0.002200 | -0.000833 | -0.000525 | -0.000314 | -0.000103 | -0.000041 | -0.000005 |
| aicc_diff | 50.0 | -44.129773 | 49.853782 | -262.197000 | -71.519624 | -55.689435 | -30.905210 | -13.263520 | -3.806752 | -0.373250 |
| aicc_diff_relative | 50.0 | -0.000410 | 0.000426 | -0.002200 | -0.000833 | -0.000525 | -0.000314 | -0.000103 | -0.000041 | -0.000005 |
| lnL_diff | 50.0 | 22.545597 | 25.445814 | 0.186630 | 1.903375 | 6.631742 | 16.447120 | 27.844717 | 35.759821 | 135.104510 |
| lnL_diff_relative | 50.0 | -0.000420 | 0.000435 | -0.002271 | -0.000836 | -0.000520 | -0.000337 | -0.000103 | -0.000041 | -0.000005 |
| unrooted_softwired_network_distance | 50.0 | 0.046303 | 0.052469 | 0.000000 | 0.000000 | 0.000000 | 0.040833 | 0.060069 | 0.119388 | 0.220000 |
| unrooted_hardwired_network_distance | 50.0 | 0.131629 | 0.074169 | 0.000000 | 0.050000 | 0.073628 | 0.119048 | 0.189369 | 0.222727 | 0.361702 |
| unrooted_displayed_trees_distance | 50.0 | 1.000000 | 0.000000 | 1.000000 | 1.000000 | 1.000000 | 1.000000 | 1.000000 | 1.000000 | 1.000000 |
| rooted_softwired_network_distance | 50.0 | 0.271294 | 0.116556 | 0.058824 | 0.122917 | 0.190617 | 0.254630 | 0.364719 | 0.447400 | 0.485714 |
| rooted_hardwired_network_distance | 50.0 | 0.221278 | 0.084798 | 0.048780 | 0.095238 | 0.181818 | 0.222222 | 0.277174 | 0.334043 | 0.408163 |
| rooted_displayed_trees_distance | 50.0 | 1.000000 | 0.000000 | 1.000000 | 1.000000 | 1.000000 | 1.000000 | 1.000000 | 1.000000 | 1.000000 |
| rooted_tripartition_distance | 50.0 | 0.233835 | 0.093854 | 0.048780 | 0.095238 | 0.181818 | 0.222222 | 0.294056 | 0.357526 | 0.431373 |
| rooted_path_multiplicity_distance | 50.0 | 0.131061 | 0.058868 | 0.024390 | 0.048193 | 0.094118 | 0.122093 | 0.159091 | 0.211978 | 0.276596 |
| rooted_nested_labels_distance | 50.0 | 0.329632 | 0.124857 | 0.090909 | 0.169855 | 0.250000 | 0.285714 | 0.415094 | 0.475455 | 0.610169 |
| runtime_raxml | 50.0 | 49.038660 | 9.746339 | 29.761000 | 37.481300 | 42.748000 | 47.852500 | 55.477750 | 58.564600 | 74.907000 |
| runtime_inference | 50.0 | 366.414520 | 309.771546 | 11.472000 | 146.491800 | 248.360250 | 324.411500 | 393.384000 | 492.009900 | 2248.341000 |

**Table 13.** Percentiles for experiment A1, 40 taxa, 2 reticulations, `LhModel.AVERAGE`, starting from RAxML-NG best tree. Top: `LhType.AVERAGE`, bottom: `LhType.BEST`.

#### 6.6 A2: 40 taxa, 3 reticulations

In this experiment, we simulated 50 networks each for 40 taxa and 3 reticulations. We ran NetRAX inference starting from the RAxML-NG best ML tree.

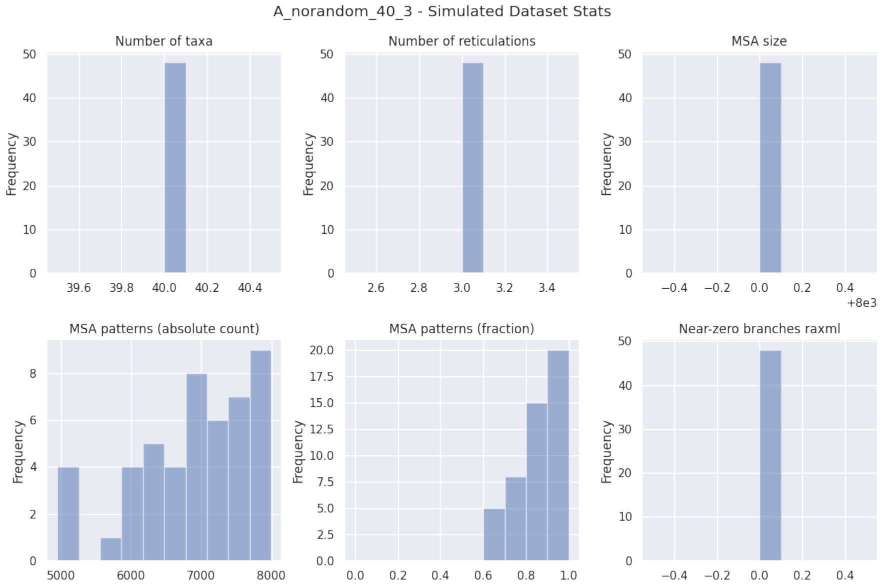


**Fig. 34.** Simulated dataset statistics for experiment A2, 40 taxa, 3 reticulations.

| A_norandom_40_3_norandom | LhType.AVERAGE | LhType.BEST |
|---|---|---|
| Inferred BIC better or equal | 2 (4.17 %) | 1 (2.08 %) |
| Inferred AIC better or equal | 2 (4.17 %) | 1 (2.08 %) |
| Inferred AICc better or equal | 2 (4.17 %) | 1 (2.08 %) |
| Inferred BIC worse | 46 (95.83 %) | 47 (97.92 %) |
| Inferred AIC worse | 46 (95.83 %) | 47 (97.92 %) |
| Inferred AICc worse | 46 (95.83 %) | 47 (97.92 %) |
| Inferred lnL better or equal | 2 (4.17 %) | 1 (2.08 %) |
| Inferred lnL worse | 46 (95.83 %) | 47 (97.92 %) |
| Inferred n_reticulations less | 2 (4.17 %) | 4 (8.33 %) |
| Inferred n_reticulations equal | 44 (91.67 %) | 44 (91.67 %) |
| Inferred n_reticulations more | 2 (4.17 %) | 0 (0.00 %) |
| Unrooted softwired distance zero | 26 (54.17 %) | 26 (54.17 %) |
| Good result | 28 (58.33 %) | 27 (56.25 %) |
| Passable result | 16 (33.33 %) | 17 (35.42 %) |
| Bad result | 4 (8.33 %) | 4 (8.33 %) |

**Table 14.** Summary statistics for experiment A1, 40 taxa, 3 reticulations, starting from RAxML-NG best tree.

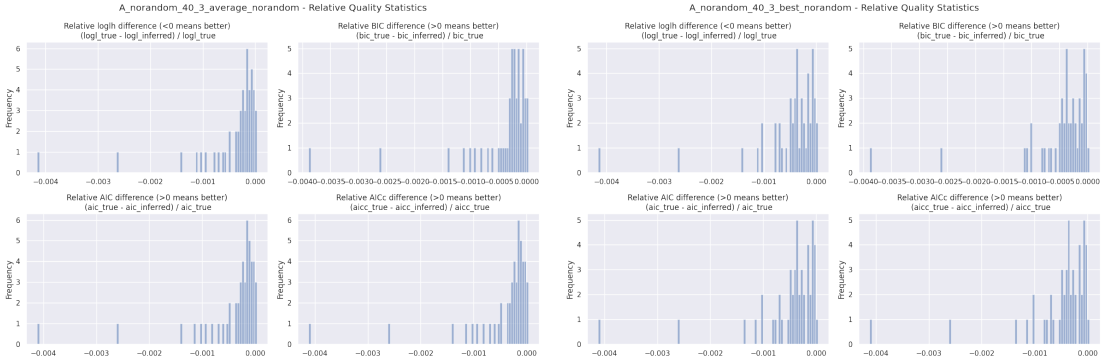


**Fig. 35.** Relative BIC, AIC, AICc, and loglikelihood differences for experiment A2, 40 taxa, 3 reticulations, starting from RAxML-NG best tree. Left: `LhType.AVERAGE`, right: `LhType.BEST`.

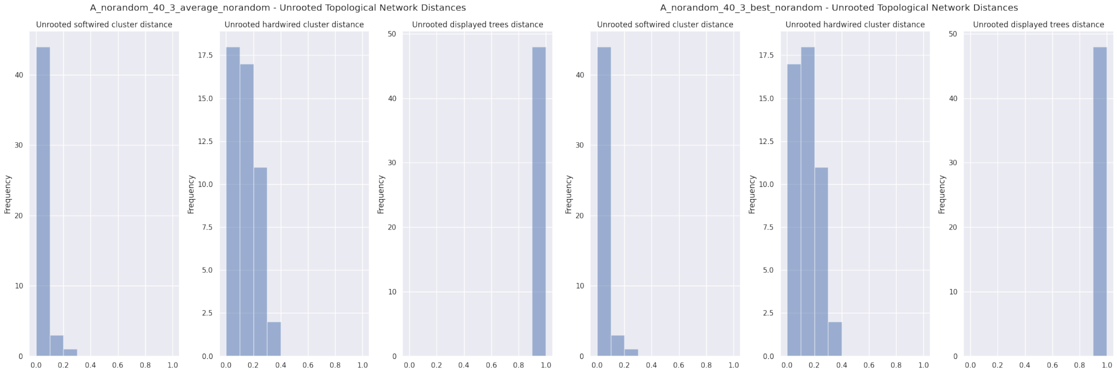


**Fig. 36.** Unrooted relative distances for experiment A2, 40 taxa, 3 reticulations, starting from RAxML-NG best tree. Left: `LhType.AVERAGE`, right: `LhType.BEST`.

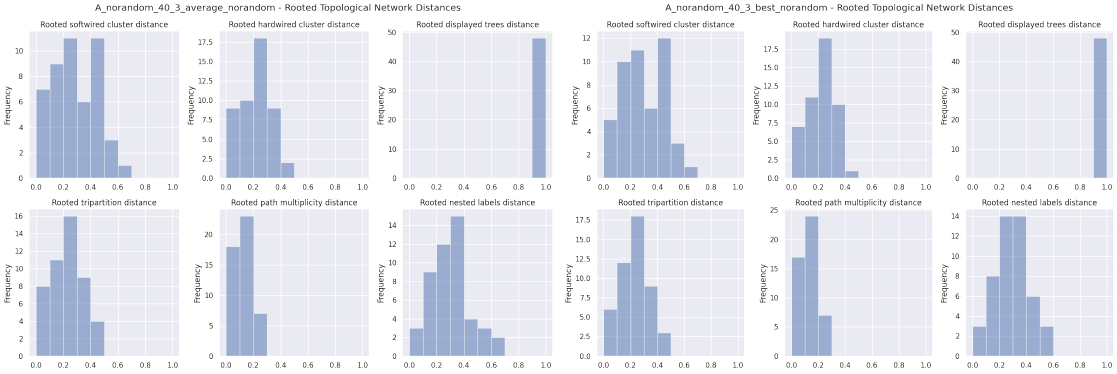


**Fig. 37.** Rooted relative distances for experiment A2, 40 taxa, 3 reticulations, starting from RAxML-NG best tree. Left: `LhType.AVERAGE`, right: `LhType.BEST`.

| | count | mean | std | min | 10% | 25% | 50% | 75% | 90% | max |
|---|---|---|---|---|---|---|---|---|---|---|
| n_reticulations_inferred | 48.0 | 3.000000 | 0.291730 | 2.000000 | 3.000000 | 3.000000 | 3.000000 | 3.000000 | 3.000000 | 4.000000 |
| bic_diff | 48.0 | -90.513285 | 145.973592 | -847.506900 | -188.465780 | -85.553025 | -50.222250 | -21.911725 | -5.932000 | 7.254900 |
| bic_diff_relative | 48.0 | -0.000421 | 0.000683 | -0.003887 | -0.000965 | -0.000373 | -0.000218 | -0.000109 | -0.000023 | 0.000034 |
| aic_diff | 48.0 | -90.513290 | 151.536831 | -890.211100 | -188.465880 | -82.270300 | -46.621100 | -21.911725 | -5.932000 | 7.254900 |
| aic_diff_relative | 48.0 | -0.000425 | 0.000712 | -0.004116 | -0.000975 | -0.000392 | -0.000209 | -0.000107 | -0.000023 | 0.000034 |
| aicc_diff | 48.0 | -90.513300 | 151.535704 | -890.203100 | -188.465880 | -82.270300 | -46.621050 | -21.911650 | -5.932070 | 7.255000 |
| aicc_diff_relative | 48.0 | -0.000425 | 0.000712 | -0.004116 | -0.000975 | -0.000392 | -0.000209 | -0.000107 | -0.000023 | 0.000034 |
| lnL_diff | 48.0 | 45.256640 | 76.334788 | -3.627400 | 2.965962 | 10.187545 | 23.310500 | 41.135150 | 94.232899 | 449.105600 |
| lnL_diff_relative | 48.0 | -0.000426 | 0.000718 | -0.004159 | -0.000977 | -0.000393 | -0.000210 | -0.000087 | -0.000023 | 0.000034 |
| unrooted_softwired_network_distance | 48.0 | 0.032212 | 0.049277 | 0.000000 | 0.000000 | 0.000000 | 0.000000 | 0.044807 | 0.091761 | 0.218750 |
| unrooted_hardwired_network_distance | 48.0 | 0.142585 | 0.084580 | 0.000000 | 0.047619 | 0.071429 | 0.137949 | 0.200000 | 0.258723 | 0.361702 |
| unrooted_displayed_trees_distance | 48.0 | 1.000000 | 0.000000 | 1.000000 | 1.000000 | 1.000000 | 1.000000 | 1.000000 | 1.000000 | 1.000000 |
| rooted_softwired_network_distance | 48.0 | 0.293043 | 0.157367 | 0.000000 | 0.089157 | 0.174860 | 0.275448 | 0.426108 | 0.493307 | 0.630435 |
| rooted_hardwired_network_distance | 48.0 | 0.216645 | 0.107189 | 0.000000 | 0.079402 | 0.131342 | 0.228261 | 0.297872 | 0.345833 | 0.411765 |
| rooted_displayed_trees_distance | 48.0 | 1.000000 | 0.000000 | 1.000000 | 1.000000 | 1.000000 | 1.000000 | 1.000000 | 1.000000 | 1.000000 |
| rooted_tripartition_distance | 48.0 | 0.228262 | 0.114580 | 0.000000 | 0.079402 | 0.153409 | 0.230676 | 0.312500 | 0.383647 | 0.452830 |
| rooted_path_multiplicity_distance | 48.0 | 0.132675 | 0.072350 | 0.000000 | 0.040562 | 0.087415 | 0.136364 | 0.177778 | 0.223142 | 0.298969 |
| rooted_nested_labels_distance | 48.0 | 0.291235 | 0.142217 | 0.000000 | 0.127660 | 0.194728 | 0.291101 | 0.370370 | 0.474471 | 0.636364 |
| runtime_raxml | 48.0 | 107.258396 | 25.381060 | 58.505000 | 77.675900 | 93.946500 | 104.149500 | 122.216500 | 144.933600 | 154.649000 |
| runtime_inference | 48.0 | 2195.188500 | 828.036918 | 811.862000 | 1270.799500 | 1655.701750 | 2008.101000 | 2518.836000 | 3226.490200 | 4744.682000 |

| | count | mean | std | min | 10% | 25% | 50% | 75% | 90% | max |
|---|---|---|---|---|---|---|---|---|---|---|
| n_reticulations_inferred | 48.0 | 2.916667 | 0.279310 | 2.000000 | 3.000000 | 3.000000 | 3.000000 | 3.000000 | 3.000000 | 3.000000 |
| bic_diff | 48.0 | -104.731731 | 144.885607 | -847.506900 | -185.244280 | -132.494275 | -64.956600 | -29.706950 | -8.470790 | 7.254900 |
| bic_diff_relative | 48.0 | -0.000477 | 0.000666 | -0.003887 | -0.001011 | -0.000489 | -0.000349 | -0.000138 | -0.000045 | 0.000034 |
| aic_diff | 48.0 | -108.290425 | 150.870389 | -890.211100 | -200.080970 | -142.470700 | -64.956550 | -29.706950 | -8.470790 | 7.254900 |
| aic_diff_relative | 48.0 | -0.000497 | 0.000700 | -0.004116 | -0.001021 | -0.000510 | -0.000352 | -0.000138 | -0.000045 | 0.000034 |
| aicc_diff | 48.0 | -108.289756 | 150.869195 | -890.203100 | -200.078500 | -142.470700 | -64.956600 | -29.706950 | -8.470790 | 7.255000 |
| aicc_diff_relative | 48.0 | -0.000497 | 0.000700 | -0.004116 | -0.001021 | -0.000510 | -0.000352 | -0.000138 | -0.000045 | 0.000034 |
| lnL_diff | 48.0 | 54.478536 | 76.034717 | -3.627400 | 4.235380 | 14.853462 | 32.478290 | 71.235350 | 101.240485 | 449.105600 |
| lnL_diff_relative | 48.0 | -0.000500 | 0.000707 | -0.004159 | -0.001023 | -0.000517 | -0.000353 | -0.000138 | -0.000045 | 0.000034 |
| unrooted_softwired_network_distance | 48.0 | 0.034945 | 0.052129 | 0.000000 | 0.000000 | 0.000000 | 0.000000 | 0.065801 | 0.093927 | 0.218750 |
| unrooted_hardwired_network_distance | 48.0 | 0.142872 | 0.081553 | 0.000000 | 0.048432 | 0.072735 | 0.130682 | 0.200000 | 0.247707 | 0.361702 |
| unrooted_displayed_trees_distance | 48.0 | 1.000000 | 0.000000 | 1.000000 | 1.000000 | 1.000000 | 1.000000 | 1.000000 | 1.000000 | 1.000000 |
| rooted_softwired_network_distance | 48.0 | 0.301610 | 0.155180 | 0.000000 | 0.104643 | 0.179165 | 0.281319 | 0.428955 | 0.493307 | 0.630435 |
| rooted_hardwired_network_distance | 48.0 | 0.217851 | 0.102573 | 0.000000 | 0.093023 | 0.136364 | 0.219807 | 0.293218 | 0.349728 | 0.400000 |
| rooted_displayed_trees_distance | 48.0 | 1.000000 | 0.000000 | 1.000000 | 1.000000 | 1.000000 | 1.000000 | 1.000000 | 1.000000 | 1.000000 |
| rooted_tripartition_distance | 48.0 | 0.229139 | 0.109174 | 0.000000 | 0.093023 | 0.153409 | 0.219807 | 0.296875 | 0.383647 | 0.431373 |
| rooted_path_multiplicity_distance | 48.0 | 0.129951 | 0.063213 | 0.000000 | 0.047619 | 0.087415 | 0.136364 | 0.162422 | 0.217391 | 0.265957 |
| rooted_nested_labels_distance | 48.0 | 0.280507 | 0.123465 | 0.000000 | 0.127660 | 0.204082 | 0.274510 | 0.370370 | 0.416842 | 0.542373 |
| runtime_raxml | 48.0 | 107.258396 | 25.381060 | 58.505000 | 77.675900 | 93.946500 | 104.149500 | 122.216500 | 144.933600 | 154.649000 |
| runtime_inference | 48.0 | 1854.266771 | 612.670255 | 483.695000 | 1232.943900 | 1529.690750 | 1840.635500 | 2090.032750 | 2540.603700 | 3756.555000 |

**Table 15.** Percentiles for experiment A1, 40 taxa, 3 reticulations, `LhModel.AVERAGE`, starting from RAxML-NG best tree. Top: `LhType.AVERAGE`, bottom: `LhType.BEST`.

#### 6.7 A2: 40 taxa, 4 reticulations

In this experiment, we simulated 50 networks each for 40 taxa and 4 reticulations. We ran NetRAX inference starting from the RAxML-NG best ML tree.

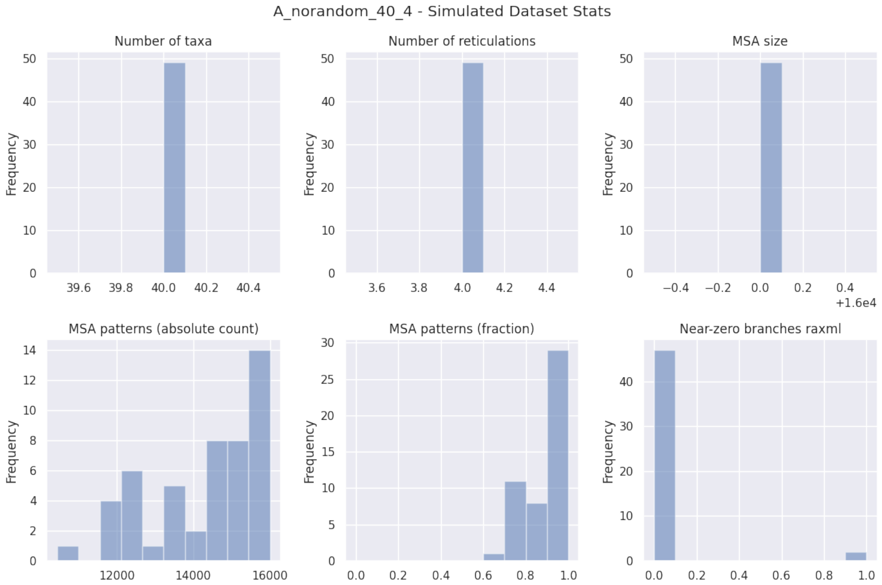


**Fig. 38.** Simulated dataset statistics for experiment A2, 40 taxa, 4 reticulations.

| A_norandom_40_4_norandom | LhType.AVERAGE | LhType.BEST |
|---|---|---|
| Inferred BIC better or equal | 0 (0.00 %) | 0 (0.00 %) |
| Inferred AIC better or equal | 0 (0.00 %) | 0 (0.00 %) |
| Inferred AICc better or equal | 0 (0.00 %) | 0 (0.00 %) |
| Inferred BIC worse | 49 (100.00 %) | 49 (100.00 %) |
| Inferred AIC worse | 49 (100.00 %) | 49 (100.00 %) |
| Inferred AICc worse | 49 (100.00 %) | 49 (100.00 %) |
| Inferred lnL better or equal | 0 (0.00 %) | 0 (0.00 %) |
| Inferred lnL worse | 49 (100.00 %) | 49 (100.00 %) |
| Inferred n_reticulations less | 5 (10.20 %) | 10 (20.41 %) |
| Inferred n_reticulations equal | 43 (87.76 %) | 38 (77.55 %) |
| Inferred n_reticulations more | 1 (2.04 %) | 1 (2.04 %) |
| Unrooted softwired distance zero | 23 (46.94 %) | 19 (38.78 %) |
| Good result | 23 (46.94 %) | 19 (38.78 %) |
| Passable result | 20 (40.82 %) | 19 (38.78 %) |
| Bad result | 6 (12.24 %) | 11 (22.45 %) |

**Table 16.** Summary statistics for experiment A1, 40 taxa, 4 reticulations, starting from RAxML-NG best tree.

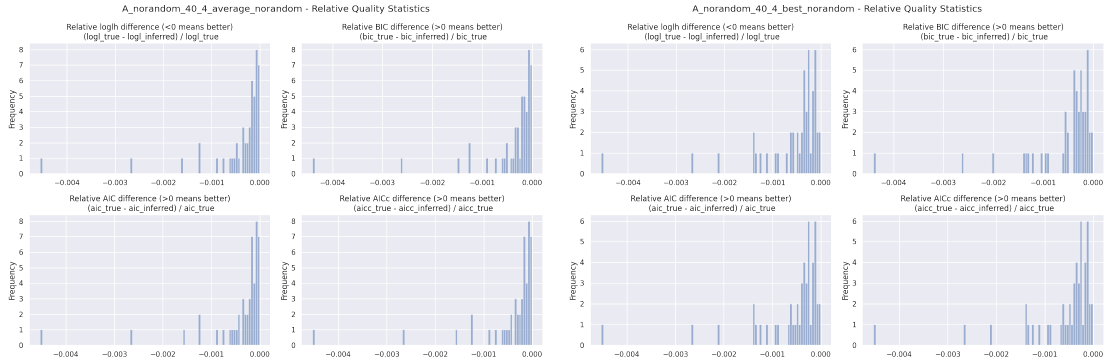


**Fig. 39.** Relative BIC, AIC, AICc, and loglikelihood differences for experiment A2, 40 taxa, 4 reticulations, starting from RAxML-NG best tree. Left: `LhType.AVERAGE`, right: `LhType.BEST`.

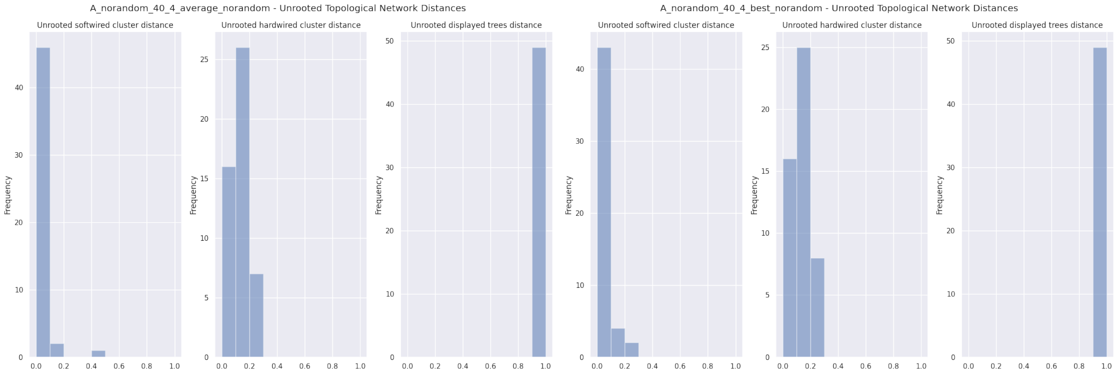


**Fig. 40.** Unrooted relative distances for experiment A2, 40 taxa, 4 reticulations, starting from RAxML-NG best tree. Left: `LhType.AVERAGE`, right: `LhType.BEST`.

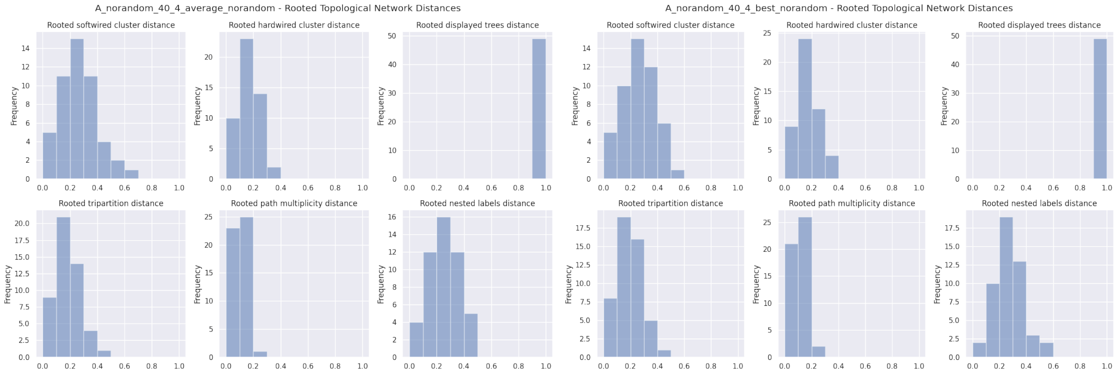


**Fig. 41.** Rooted relative distances for experiment A2, 40 taxa, 4 reticulations, starting from RAxML-NG best tree. Left: `LhType.AVERAGE`, right: `LhType.BEST`.

| | count | mean | std | min | 10% | 25% | 50% | 75% | 90% | max |
|---|---|---|---|---|---|---|---|---|---|---|
| n_reticulations_inferred | 49.0 | 3.938776 | 0.428571 | 3.000000 | 3.800000 | 4.000000 | 4.000000 | 4.000000 | 4.000000 | 6.000000 |
| bic_diff | 49.0 | -220.167016 | 426.863881 | -2252.498000 | -461.412760 | -168.759700 | -92.817200 | -36.963700 | -14.076940 | -2.016500 |
| bic_diff_relative | 49.0 | -0.000422 | 0.000745 | -0.004400 | -0.000959 | -0.000360 | -0.000182 | -0.000075 | -0.000032 | -0.000005 |
| aic_diff | 49.0 | -222.951302 | 436.370801 | -2297.974900 | -461.412760 | -165.345600 | -91.871100 | -36.963700 | -14.076940 | -2.016500 |
| aic_diff_relative | 49.0 | -0.000430 | 0.000767 | -0.004529 | -0.000963 | -0.000427 | -0.000183 | -0.000075 | -0.000032 | -0.000005 |
| aicc_diff | 49.0 | -222.950943 | 436.369523 | -2297.969000 | -461.412760 | -165.345500 | -91.883200 | -36.963700 | -14.076940 | -2.016500 |
| aicc_diff_relative | 49.0 | -0.000430 | 0.000767 | -0.004529 | -0.000963 | -0.000427 | -0.000183 | -0.000075 | -0.000032 | -0.000005 |
| lnL_diff | 49.0 | 111.720539 | 219.055762 | 1.008300 | 7.038420 | 18.481800 | 44.665400 | 84.379800 | 230.706420 | 1152.987400 |
| lnL_diff_relative | 49.0 | -0.000432 | 0.000771 | -0.004551 | -0.000964 | -0.000428 | -0.000183 | -0.000076 | -0.000032 | -0.000005 |
| unrooted_softwired_network_distance | 49.0 | 0.034855 | 0.071979 | 0.000000 | 0.000000 | 0.000000 | 0.014493 | 0.050847 | 0.070833 | 0.467213 |
| unrooted_hardwired_network_distance | 49.0 | 0.126314 | 0.062339 | 0.000000 | 0.047398 | 0.071429 | 0.133333 | 0.159091 | 0.202553 | 0.244444 |
| unrooted_displayed_trees_distance | 49.0 | 1.000000 | 0.000000 | 1.000000 | 1.000000 | 1.000000 | 1.000000 | 1.000000 | 1.000000 | 1.000000 |
| rooted_softwired_network_distance | 49.0 | 0.268700 | 0.137870 | 0.000000 | 0.103030 | 0.173913 | 0.258427 | 0.354839 | 0.434827 | 0.650000 |
| rooted_hardwired_network_distance | 49.0 | 0.169417 | 0.080610 | 0.000000 | 0.065116 | 0.133333 | 0.177778 | 0.212766 | 0.275000 | 0.380000 |
| rooted_displayed_trees_distance | 49.0 | 1.000000 | 0.000000 | 1.000000 | 1.000000 | 1.000000 | 1.000000 | 1.000000 | 1.000000 | 1.000000 |
| rooted_tripartition_distance | 49.0 | 0.181081 | 0.087584 | 0.000000 | 0.065116 | 0.133333 | 0.195652 | 0.212766 | 0.289796 | 0.425926 |
| rooted_path_multiplicity_distance | 49.0 | 0.104506 | 0.052896 | 0.000000 | 0.042409 | 0.069767 | 0.113636 | 0.134831 | 0.175824 | 0.268041 |
| rooted_nested_labels_distance | 49.0 | 0.256530 | 0.108021 | 0.000000 | 0.122449 | 0.196078 | 0.264151 | 0.327273 | 0.391531 | 0.476190 |
| runtime_raxml | 49.0 | 213.969286 | 48.157515 | 128.989000 | 154.762000 | 177.813000 | 215.941000 | 242.467000 | 262.700000 | 384.130000 |
| runtime_inference | 49.0 | 10122.056653 | 7290.758495 | 3193.147000 | 4469.655200 | 7205.559000 | 8979.243000 | 11031.439000 | 13325.616800 | 49184.610000 |

| | count | mean | std | min | 10% | 25% | 50% | 75% | 90% | max |
|---|---|---|---|---|---|---|---|---|---|---|
| n_reticulations_inferred | 49.0 | 3.816327 | 0.441280 | 3.000000 | 3.000000 | 4.000000 | 4.000000 | 4.000000 | 4.000000 | 5.000000 |
| bic_diff | 49.0 | -287.359763 | 428.140486 | -2252.493900 | -623.965080 | -313.744200 | -125.809900 | -93.061500 | -45.427360 | -3.417400 |
| bic_diff_relative | 49.0 | -0.000566 | 0.000762 | -0.004400 | -0.001299 | -0.000540 | -0.000322 | -0.000183 | -0.000108 | -0.000008 |
| aic_diff | 49.0 | -295.712667 | 437.945292 | -2297.970800 | -633.060460 | -313.744200 | -156.278600 | -93.061500 | -45.427360 | -3.417300 |
| aic_diff_relative | 49.0 | -0.000588 | 0.000786 | -0.004529 | -0.001353 | -0.000599 | -0.000327 | -0.000184 | -0.000109 | -0.000008 |
| aicc_diff | 49.0 | -295.711584 | 437.943982 | -2297.965000 | -633.059280 | -313.744100 | -156.278600 | -93.061500 | -45.427360 | -3.417300 |
| aicc_diff_relative | 49.0 | -0.000588 | 0.000786 | -0.004529 | -0.001353 | -0.000599 | -0.000327 | -0.000184 | -0.000109 | -0.000008 |
| lnL_diff | 49.0 | 148.591016 | 219.871108 | 1.708600 | 22.713640 | 46.530700 | 78.139300 | 156.872000 | 317.330180 | 1152.985500 |
| lnL_diff_relative | 49.0 | -0.000592 | 0.000790 | -0.004551 | -0.001356 | -0.000603 | -0.000328 | -0.000184 | -0.000109 | -0.000008 |
| unrooted_softwired_network_distance | 49.0 | 0.039767 | 0.054524 | 0.000000 | 0.000000 | 0.000000 | 0.016129 | 0.053333 | 0.105000 | 0.206897 |
| unrooted_hardwired_network_distance | 49.0 | 0.131180 | 0.059587 | 0.000000 | 0.047398 | 0.090909 | 0.136364 | 0.173913 | 0.213691 | 0.239130 |
| unrooted_displayed_trees_distance | 49.0 | 1.000000 | 0.000000 | 1.000000 | 1.000000 | 1.000000 | 1.000000 | 1.000000 | 1.000000 | 1.000000 |
| rooted_softwired_network_distance | 49.0 | 0.268398 | 0.120987 | 0.011236 | 0.103598 | 0.186047 | 0.263158 | 0.354839 | 0.428879 | 0.582609 |
| rooted_hardwired_network_distance | 49.0 | 0.175654 | 0.080515 | 0.023810 | 0.065116 | 0.133333 | 0.177778 | 0.212766 | 0.291667 | 0.392157 |
| rooted_displayed_trees_distance | 49.0 | 1.000000 | 0.000000 | 1.000000 | 1.000000 | 1.000000 | 1.000000 | 1.000000 | 1.000000 | 1.000000 |
| rooted_tripartition_distance | 49.0 | 0.191131 | 0.087214 | 0.023810 | 0.065116 | 0.133333 | 0.195652 | 0.234043 | 0.306122 | 0.425926 |
| rooted_path_multiplicity_distance | 49.0 | 0.109924 | 0.052649 | 0.023810 | 0.042409 | 0.069767 | 0.113636 | 0.134831 | 0.178022 | 0.260417 |
| rooted_nested_labels_distance | 49.0 | 0.272643 | 0.105583 | 0.042553 | 0.152490 | 0.200000 | 0.264151 | 0.327273 | 0.391531 | 0.593750 |
| runtime_raxml | 49.0 | 213.969286 | 48.157515 | 128.989000 | 154.762000 | 177.813000 | 215.941000 | 242.467000 | 262.700000 | 384.130000 |
| runtime_inference | 49.0 | 8576.075673 | 4941.566578 | 2048.477000 | 3836.533400 | 6411.044000 | 7452.927000 | 9907.035000 | 13058.279800 | 31124.775000 |

**Table 17.** Percentiles for experiment A1, 40 taxa, 4 reticulations, `LhModel.AVERAGE`, starting from RAxML-NG best tree. Top: `LhType.AVERAGE`, bottom: `LhType.BEST`.

#### 6.8 B: Reticulation probability 0.1

We simulated 50 data sets with 20 taxa and 1 reticulation each. We varied the first-parent probability of the simulated reticulation to be in $\{0.1, 0.2, 0.3, 0.4, 0.5\}$. Here, we present the plots and tables for reticulation probability 0.1.

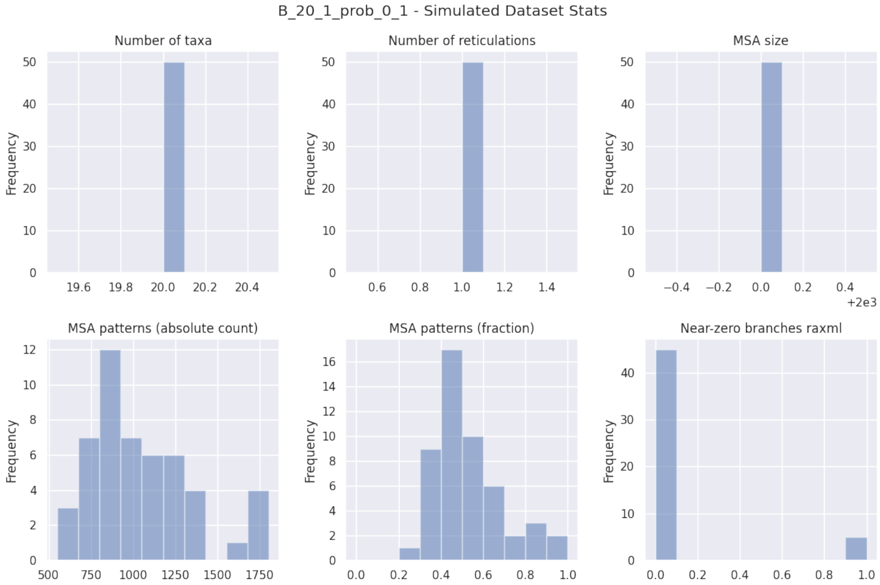


**Fig. 42.** Simulated dataset statistics for experiment B, reticulation probability 0.1.

| B_20_1_prob_0.1_norandom | LhType.AVERAGE | LhType.BEST |
|---|---|---|
| Inferred BIC better or equal | 32 (64.00 %) | 32 (64.00 %) |
| Inferred AIC better or equal | 21 (42.00 %) | 21 (42.00 %) |
| Inferred AICc better or equal | 21 (42.00 %) | 21 (42.00 %) |
| Inferred BIC worse | 18 (36.00 %) | 18 (36.00 %) |
| Inferred AIC worse | 29 (58.00 %) | 29 (58.00 %) |
| Inferred AICc worse | 29 (58.00 %) | 29 (58.00 %) |
| Inferred lnL better or equal | 18 (36.00 %) | 18 (36.00 %) |
| Inferred lnL worse | 32 (64.00 %) | 32 (64.00 %) |
| Inferred n_reticulations less | 19 (38.00 %) | 18 (36.00 %) |
| Inferred n_reticulations equal | 31 (62.00 %) | 32 (64.00 %) |
| Inferred n_reticulations more | 0 (0.00 %) | 0 (0.00 %) |
| Unrooted softwired distance zero | 11 (22.00 %) | 12 (24.00 %) |
| Good result | 36 (72.00 %) | 37 (74.00 %) |
| Passable result | 9 (18.00 %) | 9 (18.00 %) |
| Bad result | 5 (10.00 %) | 4 (8.00 %) |

**Table 18.** Summary statistics for experiment B, reticulation probability 0.1, starting from RAxML-NG best tree.

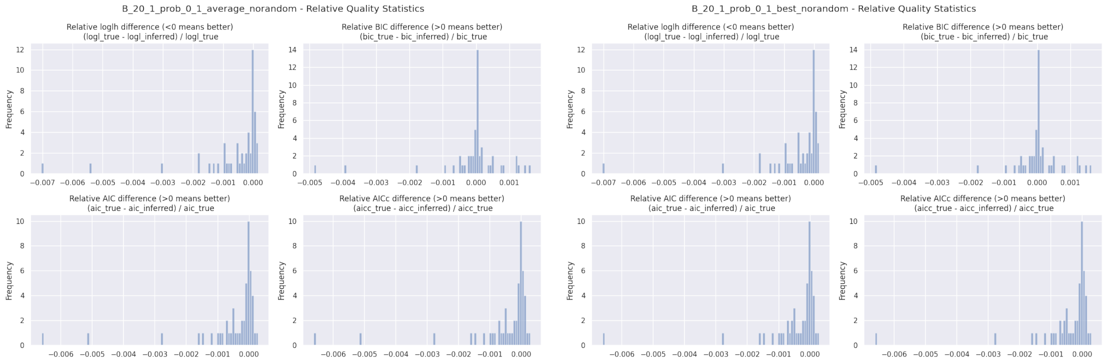


**Fig. 43.** Relative BIC, AIC, AICc, and loglikelihood differences for experiment B, reticulation probability 0.1, starting from RAxML-NG best tree. Left: `LhType.AVERAGE`, right: `LhType.BEST`.

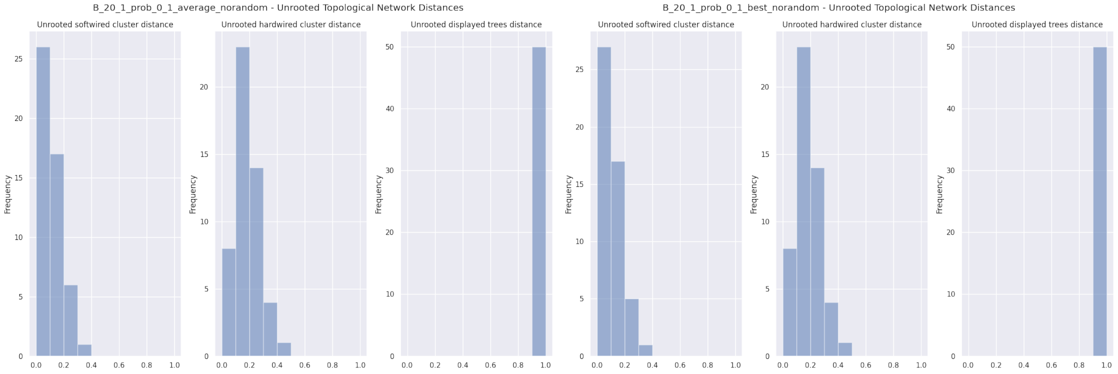


**Fig. 44.** Unrooted relative distances for experiment B, reticulation probability 0.1, starting from RAxML-NG best tree. Left: `LhType.AVERAGE`, right: `LhType.BEST`.

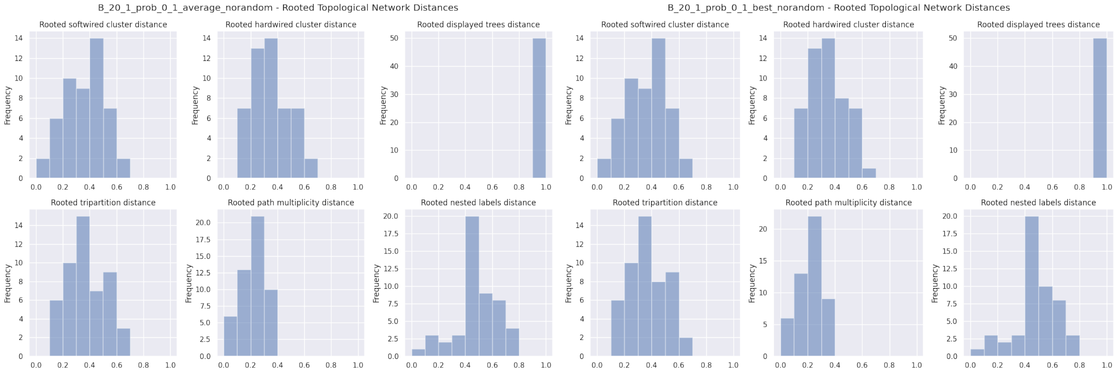


**Fig. 45.** Rooted relative distances for experiment B, reticulation probability 0.1, starting from RAxML-NG best tree. Left: `LhType.AVERAGE`, right: `LhType.BEST`.

| | count | mean | std | min | 10% | 25% | 50% | 75% | 90% | max |
|---|---|---|---|---|---|---|---|---|---|---|
| n_reticulations_inferred | 50.0 | 0.620000 | 0.490314 | 0.000000 | 0.000000 | 0.000000 | 1.000000 | 1.000000 | 1.000000 | 1.000000 |
| bic_diff | 50.0 | -2.517126 | 28.972683 | -124.743940 | -20.117562 | -4.319848 | 0.476320 | 4.958723 | 25.000693 | 41.734630 |
| bic_diff_relative | 50.0 | -0.000078 | 0.001066 | -0.004869 | -0.000511 | -0.000142 | 0.000013 | 0.000147 | 0.000863 | 0.001631 |
| aic_diff | 50.0 | -15.584010 | 33.140278 | -159.130480 | -33.122178 | -19.023172 | -2.308470 | 0.641825 | 3.032259 | 7.348100 |
| aic_diff_relative | 50.0 | -0.000533 | 0.001241 | -0.006613 | -0.001212 | -0.000576 | -0.000085 | 0.000014 | 0.000106 | 0.000293 |
| aicc_diff | 50.0 | -15.575098 | 33.134690 | -159.107020 | -33.098728 | -19.017310 | -2.296745 | 0.641825 | 3.055710 | 7.371550 |
| aicc_diff_relative | 50.0 | -0.000533 | 0.001241 | -0.006612 | -0.001211 | -0.000575 | -0.000085 | 0.000014 | 0.000107 | 0.000294 |
| lnL_diff | 50.0 | 9.312006 | 17.606977 | -2.616710 | -0.598476 | -0.152090 | 2.512375 | 12.281012 | 20.561090 | 83.565240 |
| lnL_diff_relative | 50.0 | -0.000641 | 0.001321 | -0.007031 | -0.001517 | -0.000779 | -0.000174 | 0.000011 | 0.000044 | 0.000161 |
| unrooted_softwired_network_distance | 50.0 | 0.100379 | 0.078549 | 0.000000 | 0.000000 | 0.043972 | 0.095238 | 0.150000 | 0.202727 | 0.318182 |
| unrooted_hardwired_network_distance | 50.0 | 0.176654 | 0.097821 | 0.052632 | 0.055556 | 0.105263 | 0.157895 | 0.247024 | 0.301818 | 0.478261 |
| unrooted_displayed_trees_distance | 50.0 | 1.000000 | 0.000000 | 1.000000 | 1.000000 | 1.000000 | 1.000000 | 1.000000 | 1.000000 | 1.000000 |
| rooted_softwired_network_distance | 50.0 | 0.361319 | 0.147697 | 0.041667 | 0.153462 | 0.252315 | 0.372685 | 0.477187 | 0.520000 | 0.685714 |
| rooted_hardwired_network_distance | 50.0 | 0.340342 | 0.138985 | 0.100000 | 0.145526 | 0.238095 | 0.347826 | 0.430254 | 0.520000 | 0.629630 |
| rooted_displayed_trees_distance | 50.0 | 1.000000 | 0.000000 | 1.000000 | 1.000000 | 1.000000 | 1.000000 | 1.000000 | 1.000000 | 1.000000 |
| rooted_tripartition_distance | 50.0 | 0.374285 | 0.145424 | 0.100000 | 0.186429 | 0.272727 | 0.391304 | 0.458333 | 0.542308 | 0.678571 |
| rooted_path_multiplicity_distance | 50.0 | 0.220109 | 0.090619 | 0.048780 | 0.095238 | 0.145349 | 0.222222 | 0.297872 | 0.320567 | 0.387755 |
| rooted_nested_labels_distance | 50.0 | 0.483328 | 0.158376 | 0.095238 | 0.260870 | 0.400000 | 0.480000 | 0.592593 | 0.689655 | 0.774194 |
| runtime_raxml | 50.0 | 7.897520 | 2.103311 | 4.854000 | 5.683300 | 6.233250 | 7.493500 | 8.863250 | 10.943200 | 13.061000 |
| runtime_inference | 50.0 | 17.014820 | 13.206688 | 2.248000 | 2.684400 | 3.291000 | 17.905000 | 25.517000 | 32.567200 | 58.396000 |

| | count | mean | std | min | 10% | 25% | 50% | 75% | 90% | max |
|---|---|---|---|---|---|---|---|---|---|---|
| n_reticulations_inferred | 50.0 | 0.640000 | 0.484873 | 0.000000 | 0.000000 | 0.000000 | 1.000000 | 1.000000 | 1.000000 | 1.000000 |
| bic_diff | 50.0 | -0.358814 | 23.107696 | -105.544970 | -17.108099 | -4.319848 | 0.476320 | 4.958723 | 25.000172 | 41.732630 |
| bic_diff_relative | 50.0 | -0.000010 | 0.000911 | -0.004869 | -0.000498 | -0.000142 | 0.000013 | 0.000147 | 0.000863 | 0.001631 |
| aic_diff | 50.0 | -12.737967 | 25.875181 | -139.931520 | -32.174991 | -17.199645 | -2.307325 | 0.641825 | 3.042495 | 7.346100 |
| aic_diff_relative | 50.0 | -0.000441 | 0.001050 | -0.006613 | -0.001000 | -0.000533 | -0.000085 | 0.000014 | 0.000106 | 0.000293 |
| aicc_diff | 50.0 | -12.729525 | 25.869732 | -139.908060 | -32.151541 | -17.182057 | -2.295605 | 0.641825 | 3.065946 | 7.369550 |
| aicc_diff_relative | 50.0 | -0.000441 | 0.001050 | -0.006612 | -0.000999 | -0.000533 | -0.000085 | 0.000014 | 0.000107 | 0.000294 |
| lnL_diff | 50.0 | 7.808984 | 13.971256 | -2.616710 | -0.598476 | -0.152090 | 2.509370 | 10.804457 | 20.087491 | 73.965760 |
| lnL_diff_relative | 50.0 | -0.000544 | 0.001128 | -0.007031 | -0.001343 | -0.000696 | -0.000174 | 0.000011 | 0.000044 | 0.000161 |
| unrooted_softwired_network_distance | 50.0 | 0.095834 | 0.077627 | 0.000000 | 0.000000 | 0.042120 | 0.095238 | 0.150000 | 0.200000 | 0.318182 |
| unrooted_hardwired_network_distance | 50.0 | 0.175654 | 0.097310 | 0.052632 | 0.055556 | 0.105263 | 0.157895 | 0.238095 | 0.301818 | 0.478261 |
| unrooted_displayed_trees_distance | 50.0 | 1.000000 | 0.000000 | 1.000000 | 1.000000 | 1.000000 | 1.000000 | 1.000000 | 1.000000 | 1.000000 |
| rooted_softwired_network_distance | 50.0 | 0.359595 | 0.145505 | 0.041667 | 0.153462 | 0.252315 | 0.372685 | 0.477187 | 0.501852 | 0.685714 |
| rooted_hardwired_network_distance | 50.0 | 0.336916 | 0.133720 | 0.100000 | 0.145526 | 0.238095 | 0.347826 | 0.430254 | 0.520000 | 0.615385 |
| rooted_displayed_trees_distance | 50.0 | 1.000000 | 0.000000 | 1.000000 | 1.000000 | 1.000000 | 1.000000 | 1.000000 | 1.000000 | 1.000000 |
| rooted_tripartition_distance | 50.0 | 0.369880 | 0.139222 | 0.100000 | 0.186429 | 0.272727 | 0.391304 | 0.458333 | 0.538462 | 0.678571 |
| rooted_path_multiplicity_distance | 50.0 | 0.217571 | 0.087553 | 0.048780 | 0.095238 | 0.145349 | 0.222222 | 0.294056 | 0.319149 | 0.387755 |
| rooted_nested_labels_distance | 50.0 | 0.479031 | 0.154318 | 0.095238 | 0.260870 | 0.400000 | 0.480000 | 0.587302 | 0.668966 | 0.774194 |
| runtime_raxml | 50.0 | 7.897520 | 2.103311 | 4.854000 | 5.683300 | 6.233250 | 7.493500 | 8.863250 | 10.943200 | 13.061000 |
| runtime_inference | 50.0 | 14.380420 | 11.052755 | 1.778000 | 2.135000 | 2.921250 | 14.658500 | 21.309500 | 26.829800 | 51.939000 |

**Table 19.** Percentiles for experiment B, reticulation probability 0.1, starting from RAxML-NG best tree. Top: `LhType.AVERAGE`, bottom: `LhType.BEST`.

#### 6.9 B: Reticulation probability 0.2

We simulated 50 data sets with 20 taxa and 1 reticulation each. We varied the first-parent probability of the simulated reticulation to be in $\{0.1, 0.2, 0.3, 0.4, 0.5\}$. Here, we present the plots and tables for reticulation probability 0.2.

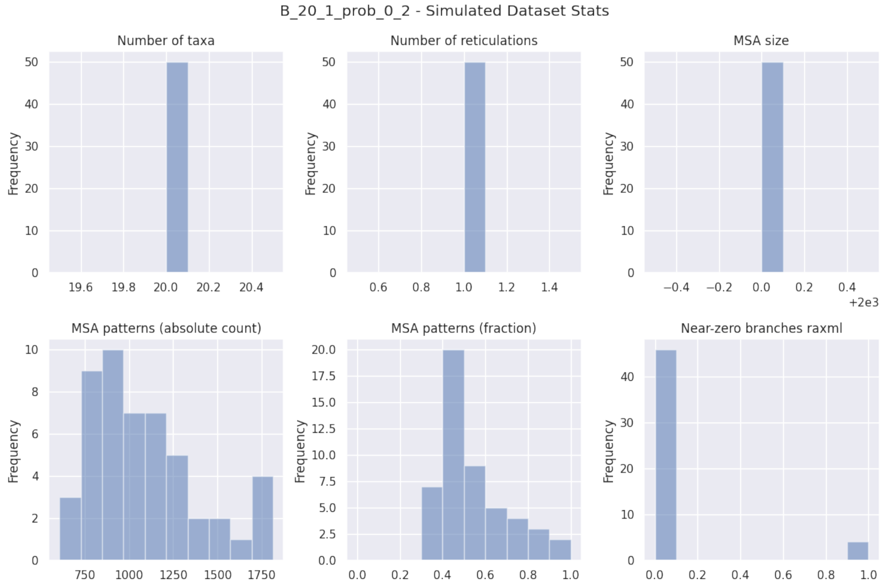


**Fig. 46.** Simulated dataset statistics for experiment B, reticulation probability 0.2.

| B_20_1_prob_0.2_norandom | LhType.AVERAGE | LhType.BEST |
|---|---|---|
| Inferred BIC better or equal | 23 (46.00 %) | 27 (54.00 %) |
| Inferred AIC better or equal | 18 (36.00 %) | 20 (40.00 %) |
| Inferred AICc better or equal | 18 (36.00 %) | 20 (40.00 %) |
| Inferred BIC worse | 27 (54.00 %) | 23 (46.00 %) |
| Inferred AIC worse | 32 (64.00 %) | 30 (60.00 %) |
| Inferred AICc worse | 32 (64.00 %) | 30 (60.00 %) |
| Inferred lnL better or equal | 18 (36.00 %) | 20 (40.00 %) |
| Inferred lnL worse | 32 (64.00 %) | 30 (60.00 %) |
| Inferred n_reticulations less | 12 (24.00 %) | 12 (24.00 %) |
| Inferred n_reticulations equal | 38 (76.00 %) | 38 (76.00 %) |
| Inferred n_reticulations more | 0 (0.00 %) | 0 (0.00 %) |
| Unrooted softwired distance zero | 17 (34.00 %) | 17 (34.00 %) |
| Good result | 36 (72.00 %) | 38 (76.00 %) |
| Passable result | 7 (14.00 %) | 7 (14.00 %) |
| Bad result | 7 (14.00 %) | 5 (10.00 %) |

**Table 20.** Summary statistics for experiment B, reticulation probability 0.2, starting from RAxML-NG best tree.

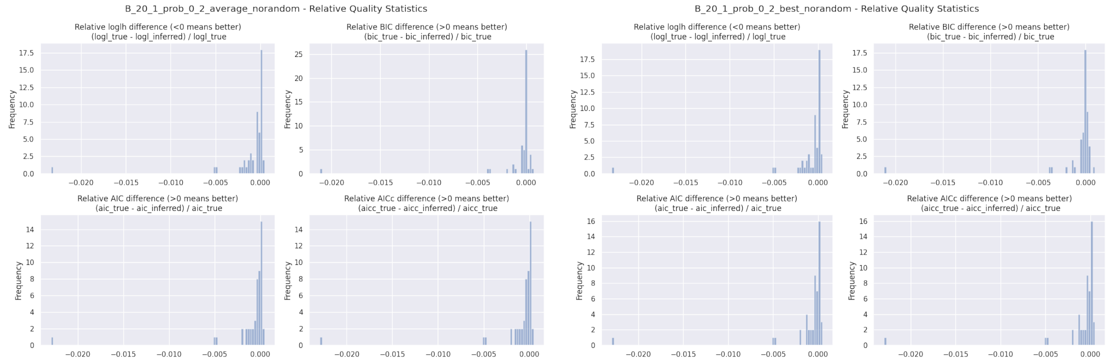


**Fig. 47.** Relative BIC, AIC, AICc, and loglikelihood differences for experiment B, reticulation probability 0.2, starting from RAxML-NG best tree. Left: `LhType.AVERAGE`, right: `LhType.BEST`.

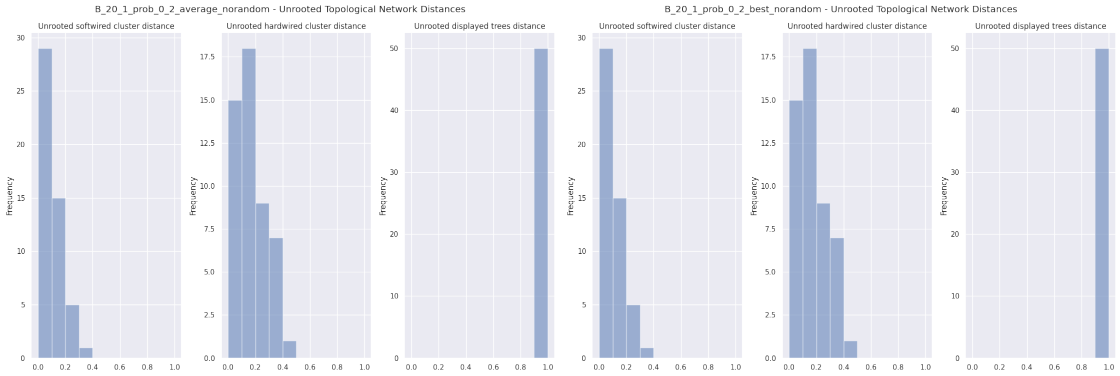


**Fig. 48.** Unrooted relative distances for experiment B, reticulation probability 0.2, starting from RAxML-NG best tree. Left: `LhType.AVERAGE`, right: `LhType.BEST`.

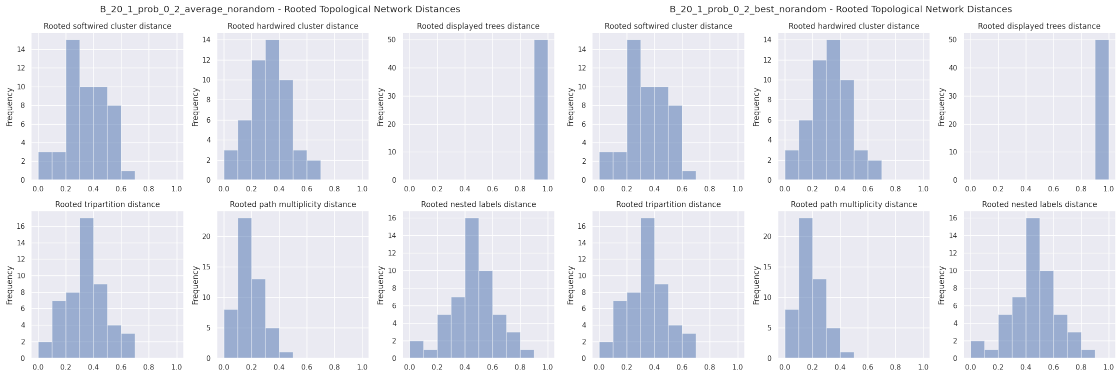


**Fig. 49.** Rooted relative distances for experiment B, reticulation probability 0.2, starting from RAxML-NG best tree. Left: `LhType.AVERAGE`, right: `LhType.BEST`.

| | count | mean | std | min | 10% | 25% | 50% | 75% | 90% | max |
|---|---|---|---|---|---|---|---|---|---|---|
| n_reticulations_inferred | 50.0 | 0.760000 | 0.431419 | 0.000000 | 0.000000 | 1.000000 | 1.000000 | 1.000000 | 1.000000 | 1.000000 |
| bic_diff | 50.0 | -20.665995 | 82.835832 | -559.281350 | -38.793778 | -9.851595 | -0.130090 | 1.362605 | 3.671201 | 20.180490 |
| bic_diff_relative | 50.0 | -0.000704 | 0.003075 | -0.021181 | -0.001345 | -0.000353 | -0.000004 | 0.000050 | 0.000127 | 0.000760 |
| aic_diff | 50.0 | -28.918766 | 88.996210 | -593.667900 | -40.188900 | -24.108600 | -4.043935 | 0.793175 | 2.018602 | 10.477780 |
| aic_diff_relative | 50.0 | -0.000991 | 0.003337 | -0.022932 | -0.001495 | -0.000821 | -0.000110 | 0.000020 | 0.000059 | 0.000469 |
| aicc_diff | 50.0 | -28.913137 | 88.991310 | -593.644440 | -40.186546 | -24.085150 | -4.043935 | 0.793175 | 2.018602 | 10.477780 |
| aicc_diff_relative | 50.0 | -0.000991 | 0.003337 | -0.022930 | -0.001494 | -0.000820 | -0.000110 | 0.000020 | 0.000059 | 0.000469 |
| lnL_diff | 50.0 | 15.419383 | 45.358871 | -5.238890 | -1.009301 | -0.396588 | 2.021970 | 16.054300 | 21.928024 | 300.833950 |
| lnL_diff_relative | 50.0 | -0.001059 | 0.003409 | -0.023349 | -0.001767 | -0.001031 | -0.000110 | 0.000020 | 0.000060 | 0.000472 |
| unrooted_softwired_network_distance | 50.0 | 0.090563 | 0.085399 | 0.000000 | 0.000000 | 0.000000 | 0.088933 | 0.155921 | 0.200833 | 0.320000 |
| unrooted_hardwired_network_distance | 50.0 | 0.157603 | 0.119296 | 0.000000 | 0.000000 | 0.055556 | 0.157895 | 0.238095 | 0.336364 | 0.434783 |
| unrooted_displayed_trees_distance | 50.0 | 1.000000 | 0.000000 | 1.000000 | 1.000000 | 1.000000 | 1.000000 | 1.000000 | 1.000000 | 1.000000 |
| rooted_softwired_network_distance | 50.0 | 0.342201 | 0.142879 | 0.000000 | 0.160000 | 0.243386 | 0.346154 | 0.440000 | 0.521846 | 0.606061 |
| rooted_hardwired_network_distance | 50.0 | 0.323705 | 0.139250 | 0.000000 | 0.186429 | 0.238095 | 0.318182 | 0.430254 | 0.484000 | 0.629630 |
| rooted_displayed_trees_distance | 50.0 | 1.000000 | 0.000000 | 1.000000 | 1.000000 | 1.000000 | 1.000000 | 1.000000 | 1.000000 | 1.000000 |
| rooted_tripartition_distance | 50.0 | 0.345880 | 0.145925 | 0.000000 | 0.186429 | 0.272727 | 0.347826 | 0.458333 | 0.521846 | 0.678571 |
| rooted_path_multiplicity_distance | 50.0 | 0.196894 | 0.090695 | 0.000000 | 0.095238 | 0.139535 | 0.181818 | 0.256763 | 0.322651 | 0.420000 |
| rooted_nested_labels_distance | 50.0 | 0.456839 | 0.161864 | 0.095238 | 0.260870 | 0.333333 | 0.461538 | 0.538462 | 0.668966 | 0.848485 |
| runtime_raxml | 50.0 | 8.168720 | 2.264444 | 4.673000 | 5.734900 | 6.323250 | 7.643000 | 9.643750 | 11.734100 | 14.073000 |
| runtime_inference | 50.0 | 18.324500 | 16.329085 | 2.453000 | 2.883700 | 8.247500 | 18.034500 | 22.535500 | 26.765400 | 110.669000 |

| | count | mean | std | min | 10% | 25% | 50% | 75% | 90% | max |
|---|---|---|---|---|---|---|---|---|---|---|
| n_reticulations_inferred | 50.0 | 0.760000 | 0.431419 | 0.000000 | 0.000000 | 1.000000 | 1.000000 | 1.000000 | 1.000000 | 1.000000 |
| bic_diff | 50.0 | -19.628502 | 83.094761 | -559.281350 | -38.793778 | -9.851595 | 0.186135 | 2.979415 | 7.769495 | 25.708880 |
| bic_diff_relative | 50.0 | -0.000672 | 0.003084 | -0.021181 | -0.001280 | -0.000353 | 0.000006 | 0.000099 | 0.000291 | 0.000968 |
| aic_diff | 50.0 | -27.881272 | 89.153979 | -593.667900 | -39.316886 | -22.556590 | -4.043935 | 1.362607 | 5.317257 | 10.477780 |
| aic_diff_relative | 50.0 | -0.000959 | 0.003343 | -0.022932 | -0.001445 | -0.000754 | -0.000110 | 0.000049 | 0.000133 | 0.000469 |
| aicc_diff | 50.0 | -27.875644 | 89.149145 | -593.644440 | -39.295780 | -22.533140 | -4.043935 | 1.362608 | 5.317257 | 10.477780 |
| aicc_diff_relative | 50.0 | -0.000959 | 0.003343 | -0.022930 | -0.001445 | -0.000753 | -0.000110 | 0.000049 | 0.000133 | 0.000469 |
| lnL_diff | 50.0 | 14.900636 | 45.426751 | -5.238890 | -2.658638 | -0.681297 | 2.021970 | 15.278303 | 20.962588 | 300.833950 |
| lnL_diff_relative | 50.0 | -0.001027 | 0.003414 | -0.023349 | -0.001721 | -0.001016 | -0.000110 | 0.000049 | 0.000134 | 0.000472 |
| unrooted_softwired_network_distance | 50.0 | 0.090563 | 0.085399 | 0.000000 | 0.000000 | 0.000000 | 0.088933 | 0.155921 | 0.200833 | 0.320000 |
| unrooted_hardwired_network_distance | 50.0 | 0.157603 | 0.119296 | 0.000000 | 0.000000 | 0.055556 | 0.157895 | 0.238095 | 0.336364 | 0.434783 |
| unrooted_displayed_trees_distance | 50.0 | 1.000000 | 0.000000 | 1.000000 | 1.000000 | 1.000000 | 1.000000 | 1.000000 | 1.000000 | 1.000000 |
| rooted_softwired_network_distance | 50.0 | 0.342201 | 0.142879 | 0.000000 | 0.160000 | 0.243386 | 0.346154 | 0.440000 | 0.521846 | 0.606061 |
| rooted_hardwired_network_distance | 50.0 | 0.323705 | 0.139250 | 0.000000 | 0.186429 | 0.238095 | 0.318182 | 0.430254 | 0.484000 | 0.629630 |
| rooted_displayed_trees_distance | 50.0 | 1.000000 | 0.000000 | 1.000000 | 1.000000 | 1.000000 | 1.000000 | 1.000000 | 1.000000 | 1.000000 |
| rooted_tripartition_distance | 50.0 | 0.345880 | 0.145925 | 0.000000 | 0.186429 | 0.272727 | 0.347826 | 0.458333 | 0.521846 | 0.678571 |
| rooted_path_multiplicity_distance | 50.0 | 0.196894 | 0.090695 | 0.000000 | 0.095238 | 0.139535 | 0.181818 | 0.256763 | 0.322651 | 0.420000 |
| rooted_nested_labels_distance | 50.0 | 0.456839 | 0.161864 | 0.095238 | 0.260870 | 0.333333 | 0.461538 | 0.538462 | 0.668966 | 0.848485 |
| runtime_raxml | 50.0 | 8.168720 | 2.264444 | 4.673000 | 5.734900 | 6.323250 | 7.643000 | 9.643750 | 11.734100 | 14.073000 |
| runtime_inference | 50.0 | 15.093820 | 14.503797 | 1.994000 | 2.253700 | 6.000500 | 13.486500 | 18.380500 | 22.333700 | 98.793000 |

**Table 21.** Percentiles for experiment B, reticulation probability 0.2, starting from RAxML-NG best tree. Top: `LhType.AVERAGE`, bottom: `LhType.BEST`.

#### 6.10 B: Reticulation probability 0.3

We simulated 50 data sets with 20 taxa and 1 reticulation each. We varied the first-parent probability of the simulated reticulation to be in $\{0.1, 0.2, 0.3, 0.4, 0.5\}$. Here, we present the plots and tables for reticulation probability 0.3.

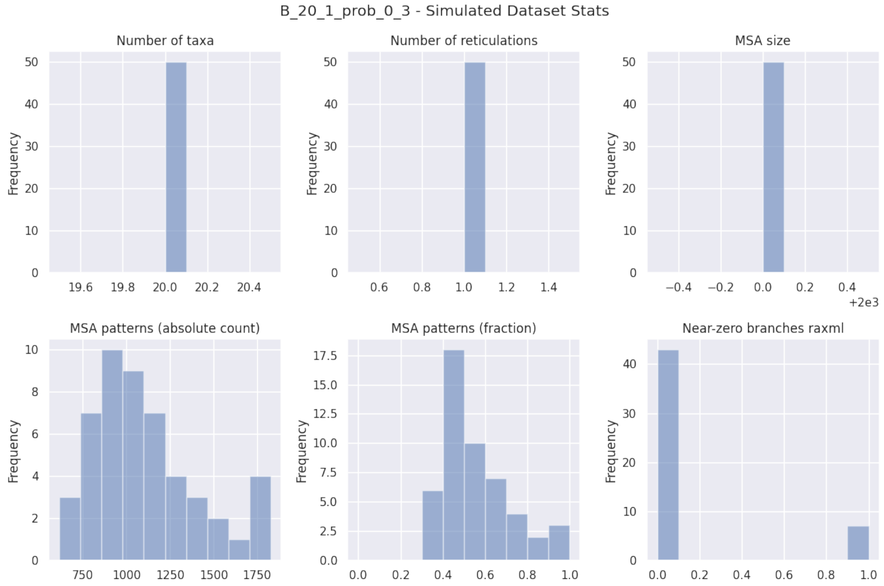


**Fig. 50.** Simulated dataset statistics for experiment B, reticulation probability 0.3.

| B_20_1_prob_0.3_norandom | LhType.AVERAGE | LhType.BEST |
|---|---|---|
| Inferred BIC better or equal | 22 (44.00 %) | 28 (56.00 %) |
| Inferred AIC better or equal | 22 (44.00 %) | 28 (56.00 %) |
| Inferred AICc better or equal | 22 (44.00 %) | 28 (56.00 %) |
| Inferred BIC worse | 28 (56.00 %) | 22 (44.00 %) |
| Inferred AIC worse | 28 (56.00 %) | 22 (44.00 %) |
| Inferred AICc worse | 28 (56.00 %) | 22 (44.00 %) |
| Inferred lnL better or equal | 22 (44.00 %) | 28 (56.00 %) |
| Inferred lnL worse | 28 (56.00 %) | 22 (44.00 %) |
| Inferred n_reticulations less | 4 (8.00 %) | 1 (2.00 %) |
| Inferred n_reticulations equal | 46 (92.00 %) | 49 (98.00 %) |
| Inferred n_reticulations more | 0 (0.00 %) | 0 (0.00 %) |
| Unrooted softwired distance zero | 26 (52.00 %) | 29 (58.00 %) |
| Good result | 36 (72.00 %) | 42 (84.00 %) |
| Passable result | 10 (20.00 %) | 7 (14.00 %) |
| Bad result | 4 (8.00 %) | 1 (2.00 %) |

**Table 22.** Summary statistics for experiment B, reticulation probability 0.3, starting from RAxML-NG best tree.

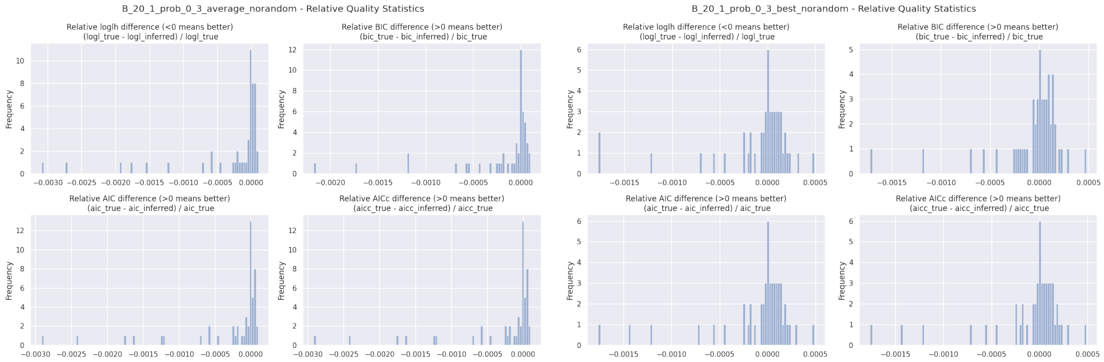


**Fig. 51.** Relative BIC, AIC, AICc, and loglikelihood differences for experiment B, reticulation probability 0.3, starting from RAxML-NG best tree. Left: `LhType.AVERAGE`, right: `LhType.BEST`.

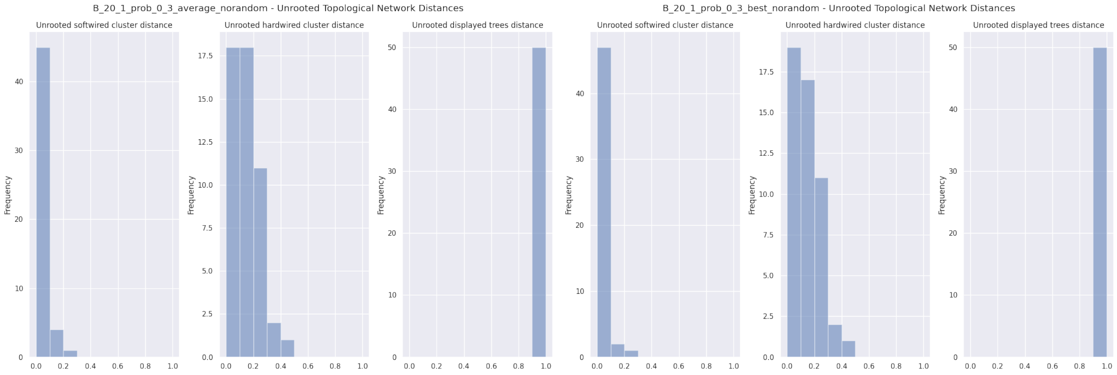


**Fig. 52.** Unrooted relative distances for experiment B, reticulation probability 0.3, starting from RAxML-NG best tree. Left: `LhType.AVERAGE`, right: `LhType.BEST`.

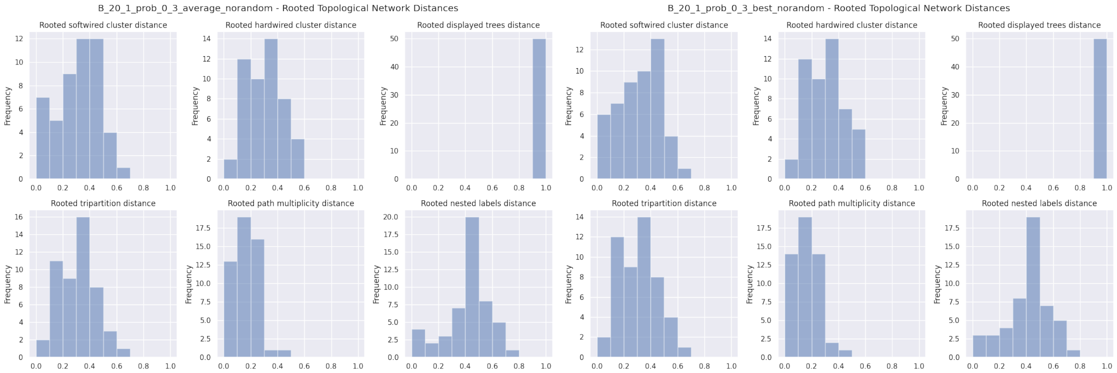


**Fig. 53.** Rooted relative distances for experiment B, reticulation probability 0.3, starting from RAxML-NG best tree. Left: `LhType.AVERAGE`, right: `LhType.BEST`.

| | count | mean | std | min | 10% | 25% | 50% | 75% | 90% | max |
|---|---|---|---|---|---|---|---|---|---|---|
| n_reticulations_inferred | 50.0 | 0.920000 | 0.274048 | 0.000000 | 1.000000 | 1.000000 | 1.000000 | 1.000000 | 1.000000 | 1.000000 |
| bic_diff | 50.0 | -6.055445 | 16.911902 | -104.874400 | -17.526060 | -5.369480 | -0.158660 | 0.634770 | 1.795278 | 3.831890 |
| bic_diff_relative | 50.0 | -0.000194 | 0.000460 | -0.002172 | -0.000585 | -0.000184 | -0.000006 | 0.000021 | 0.000053 | 0.000099 |
| aic_diff | 50.0 | -8.806368 | 23.445771 | -139.260950 | -31.465800 | -6.179330 | -0.158655 | 0.634770 | 1.795278 | 3.831900 |
| aic_diff_relative | 50.0 | -0.000286 | 0.000651 | -0.002915 | -0.001219 | -0.000205 | -0.000006 | 0.000022 | 0.000055 | 0.000101 |
| aicc_diff | 50.0 | -8.804492 | 23.440644 | -139.237490 | -31.463456 | -6.179330 | -0.158655 | 0.634770 | 1.795278 | 3.831900 |
| aicc_diff_relative | 50.0 | -0.000286 | 0.000651 | -0.002915 | -0.001219 | -0.000205 | -0.000006 | 0.000022 | 0.000055 | 0.000101 |
| lnL_diff | 50.0 | 4.723184 | 12.614658 | -1.915950 | -0.897634 | -0.317385 | 0.079335 | 3.089658 | 15.734859 | 73.630470 |
| lnL_diff_relative | 50.0 | -0.000308 | 0.000709 | -0.003091 | -0.001253 | -0.000206 | -0.000006 | 0.000022 | 0.000055 | 0.000101 |
| unrooted_softwired_network_distance | 50.0 | 0.040191 | 0.051885 | 0.000000 | 0.000000 | 0.000000 | 0.000000 | 0.079106 | 0.095714 | 0.227273 |
| unrooted_hardwired_network_distance | 50.0 | 0.127607 | 0.102095 | 0.000000 | 0.000000 | 0.053363 | 0.105263 | 0.200000 | 0.253571 | 0.409091 |
| unrooted_displayed_trees_distance | 50.0 | 1.000000 | 0.000000 | 1.000000 | 1.000000 | 1.000000 | 1.000000 | 1.000000 | 1.000000 | 1.000000 |
| rooted_softwired_network_distance | 50.0 | 0.316842 | 0.151366 | 0.000000 | 0.094410 | 0.206818 | 0.320714 | 0.420756 | 0.488172 | 0.606061 |
| rooted_hardwired_network_distance | 50.0 | 0.294652 | 0.135998 | 0.000000 | 0.100000 | 0.190476 | 0.318182 | 0.391304 | 0.437138 | 0.560000 |
| rooted_displayed_trees_distance | 50.0 | 1.000000 | 0.000000 | 1.000000 | 1.000000 | 1.000000 | 1.000000 | 1.000000 | 1.000000 | 1.000000 |
| rooted_tripartition_distance | 50.0 | 0.308566 | 0.137040 | 0.000000 | 0.100000 | 0.202381 | 0.318182 | 0.391304 | 0.458333 | 0.615385 |
| rooted_path_multiplicity_distance | 50.0 | 0.172429 | 0.084171 | 0.000000 | 0.048780 | 0.106312 | 0.181818 | 0.222222 | 0.263043 | 0.420000 |
| rooted_nested_labels_distance | 50.0 | 0.409699 | 0.151512 | 0.095238 | 0.181818 | 0.333333 | 0.400000 | 0.518519 | 0.620690 | 0.774194 |
| runtime_raxml | 50.0 | 8.360280 | 2.374050 | 5.034000 | 6.182000 | 6.654250 | 7.533000 | 9.792000 | 12.513500 | 14.229000 |
| runtime_inference | 50.0 | 20.311020 | 9.062882 | 2.427000 | 14.252400 | 16.126500 | 20.659500 | 24.285000 | 28.329600 | 62.442000 |

| | count | mean | std | min | 10% | 25% | 50% | 75% | 90% | max |
|---|---|---|---|---|---|---|---|---|---|---|
| n_reticulations_inferred | 50.0 | 0.980000 | 0.141421 | 0.000000 | 1.000000 | 1.000000 | 1.000000 | 1.000000 | 1.000000 | 1.000000 |
| bic_diff | 50.0 | -1.268297 | 8.784703 | -35.580720 | -9.571771 | -2.070683 | 0.403440 | 3.579005 | 5.926375 | 13.215820 |
| bic_diff_relative | 50.0 | -0.000062 | 0.000353 | -0.001720 | -0.000271 | -0.000060 | 0.000013 | 0.000102 | 0.000169 | 0.000478 |
| aic_diff | 50.0 | -1.956028 | 10.229879 | -38.384210 | -10.719141 | -2.070675 | 0.403440 | 3.579000 | 5.926367 | 13.215820 |
| aic_diff_relative | 50.0 | -0.000089 | 0.000410 | -0.001764 | -0.000467 | -0.000061 | 0.000013 | 0.000103 | 0.000173 | 0.000487 |
| aicc_diff | 50.0 | -1.955558 | 10.228175 | -38.360750 | -10.719140 | -2.070675 | 0.403445 | 3.579000 | 5.926367 | 13.215820 |
| aicc_diff_relative | 50.0 | -0.000089 | 0.000410 | -0.001764 | -0.000467 | -0.000061 | 0.000013 | 0.000103 | 0.000173 | 0.000487 |
| lnL_diff | 50.0 | 1.058013 | 5.427370 | -6.607920 | -2.963183 | -1.789502 | -0.201720 | 1.035340 | 5.359570 | 23.192100 |
| lnL_diff_relative | 50.0 | -0.000096 | 0.000434 | -0.001775 | -0.000470 | -0.000061 | 0.000013 | 0.000104 | 0.000173 | 0.000489 |
| unrooted_softwired_network_distance | 50.0 | 0.033817 | 0.048739 | 0.000000 | 0.000000 | 0.000000 | 0.000000 | 0.051974 | 0.095238 | 0.227273 |
| unrooted_hardwired_network_distance | 50.0 | 0.124273 | 0.104715 | 0.000000 | 0.000000 | 0.052632 | 0.105263 | 0.200000 | 0.253571 | 0.409091 |
| unrooted_displayed_trees_distance | 50.0 | 1.000000 | 0.000000 | 1.000000 | 1.000000 | 1.000000 | 1.000000 | 1.000000 | 1.000000 | 1.000000 |
| rooted_softwired_network_distance | 50.0 | 0.317127 | 0.153666 | 0.000000 | 0.095238 | 0.192857 | 0.320714 | 0.421474 | 0.488172 | 0.647059 |
| rooted_hardwired_network_distance | 50.0 | 0.298276 | 0.136503 | 0.000000 | 0.104737 | 0.190476 | 0.318182 | 0.391304 | 0.462500 | 0.560000 |
| rooted_displayed_trees_distance | 50.0 | 1.000000 | 0.000000 | 1.000000 | 1.000000 | 1.000000 | 1.000000 | 1.000000 | 1.000000 | 1.000000 |
| rooted_tripartition_distance | 50.0 | 0.307290 | 0.138209 | 0.000000 | 0.100000 | 0.190476 | 0.318182 | 0.410326 | 0.464500 | 0.615385 |
| rooted_path_multiplicity_distance | 50.0 | 0.172482 | 0.085917 | 0.000000 | 0.048780 | 0.095238 | 0.181818 | 0.222222 | 0.297872 | 0.420000 |
| rooted_nested_labels_distance | 50.0 | 0.404623 | 0.149477 | 0.095238 | 0.181818 | 0.333333 | 0.400000 | 0.504274 | 0.620690 | 0.774194 |
| runtime_raxml | 50.0 | 8.360280 | 2.374050 | 5.034000 | 6.182000 | 6.654250 | 7.533000 | 9.792000 | 12.513500 | 14.229000 |
| runtime_inference | 50.0 | 17.662660 | 7.406696 | 1.997000 | 11.787100 | 12.835750 | 16.723000 | 20.585250 | 24.931700 | 55.455000 |

**Table 23.** Percentiles for experiment B, reticulation probability 0.3, starting from RAxML-NG best tree. Top: `LhType.AVERAGE`, bottom: `LhType.BEST`.

##### 6.11 B: Reticulation probability 0.4

We simulated 50 data sets with 20 taxa and 1 reticulation each. We varied the first-parent probability of the simulated reticulation to be in $\{0.1, 0.2, 0.3, 0.4, 0.5\}$. Here, we present the plots and tables for reticulation probability 0.4.

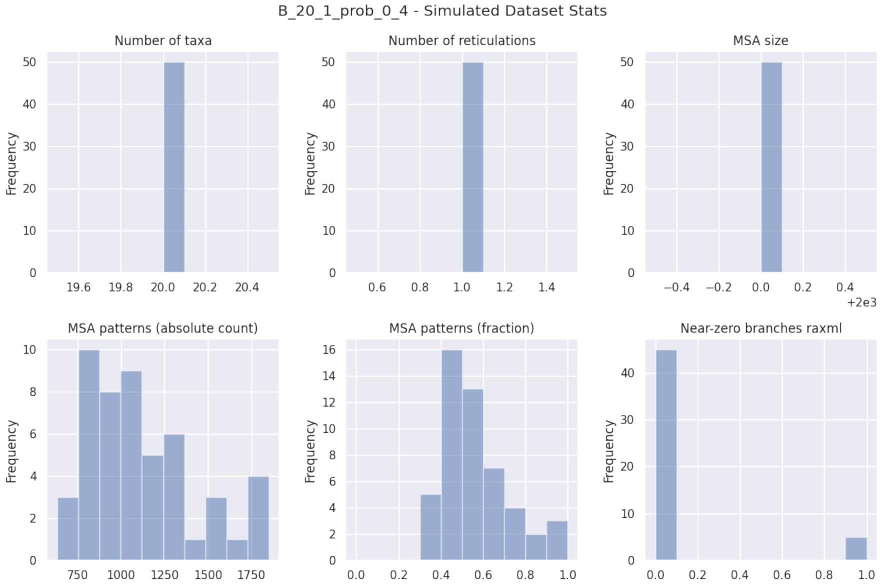


**Fig. 54.** Simulated dataset statistics for experiment B, reticulation probability 0.4.

| B_20_1_prob_0.4_norandom | LhType.AVERAGE | LhType.BEST |
|---|---|---|
| Inferred BIC better or equal | 20 (40.00 %) | 25 (50.00 %) |
| Inferred AIC better or equal | 19 (38.00 %) | 24 (48.00 %) |
| Inferred AICc better or equal | 19 (38.00 %) | 24 (48.00 %) |
| Inferred BIC worse | 30 (60.00 %) | 25 (50.00 %) |
| Inferred AIC worse | 31 (62.00 %) | 26 (52.00 %) |
| Inferred AICc worse | 31 (62.00 %) | 26 (52.00 %) |
| Inferred lnL better or equal | 19 (38.00 %) | 24 (48.00 %) |
| Inferred lnL worse | 31 (62.00 %) | 26 (52.00 %) |
| Inferred n_reticulations less | 7 (14.00 %) | 2 (4.00 %) |
| Inferred n_reticulations equal | 43 (86.00 %) | 48 (96.00 %) |
| Inferred n_reticulations more | 0 (0.00 %) | 0 (0.00 %) |
| Unrooted softwired distance zero | 23 (46.00 %) | 25 (50.00 %) |
| Good result | 34 (68.00 %) | 37 (74.00 %) |
| Passable result | 10 (20.00 %) | 12 (24.00 %) |
| Bad result | 6 (12.00 %) | 1 (2.00 %) |

**Table 24.** Summary statistics for experiment B, reticulation probability 0.4, starting from RAxML-NG best tree.

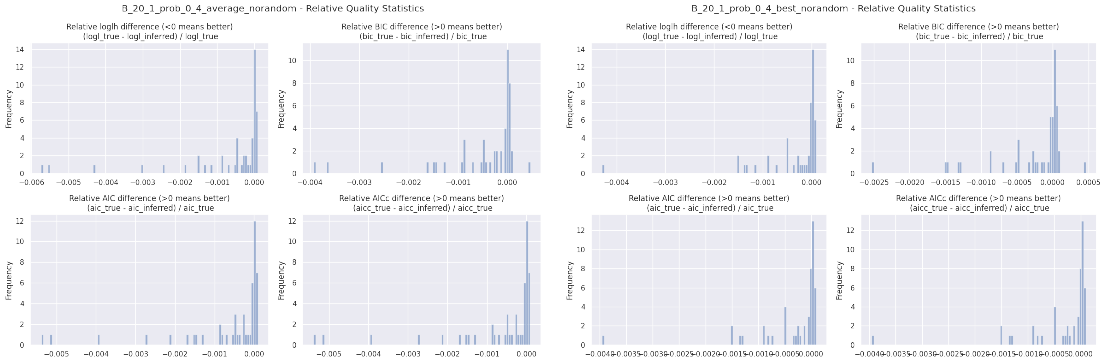


**Fig. 55.** Relative BIC, AIC, AICc, and loglikelihood differences for experiment B, reticulation probability 0.4, starting from RAxML-NG best tree. Left: `LhType.AVERAGE`, right: `LhType.BEST`.

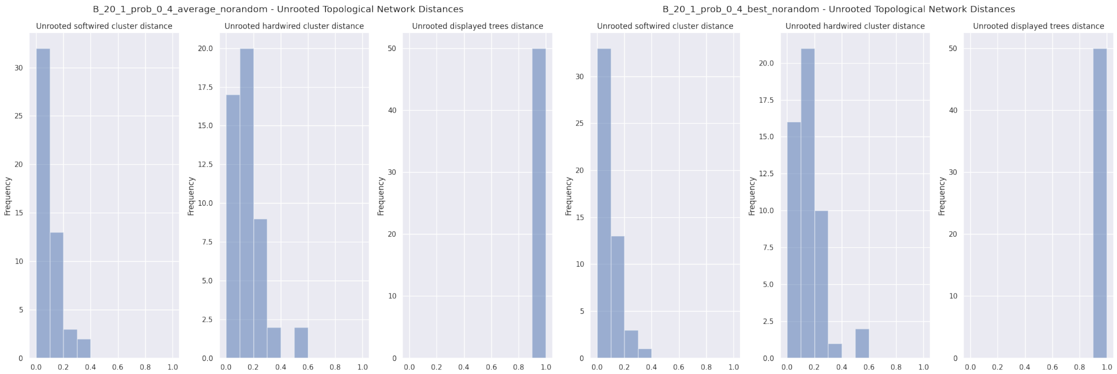


**Fig. 56.** Unrooted relative distances for experiment B, reticulation probability 0.4, starting from RAxML-NG best tree. Left: `LhType.AVERAGE`, right: `LhType.BEST`.

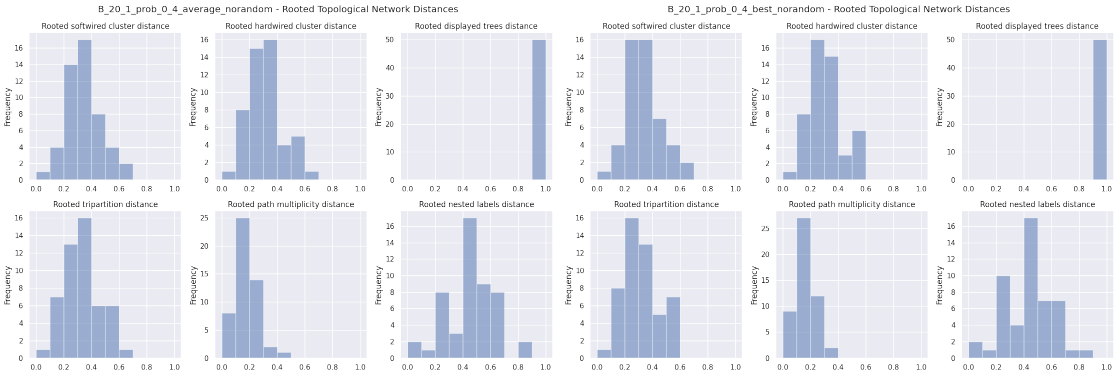


**Fig. 57.** Rooted relative distances for experiment B, reticulation probability 0.4, starting from RAxML-NG best tree. Left: `LhType.AVERAGE`, right: `LhType.BEST`.

| | count | mean | std | min | 10% | 25% | 50% | 75% | 90% | max |
|---|---|---|---|---|---|---|---|---|---|---|
| n_reticulations_inferred | 50.0 | 0.860000 | 0.350510 | 0.000000 | 0.000000 | 1.000000 | 1.000000 | 1.000000 | 1.000000 | 1.000000 |
| bic_diff | 50.0 | -13.170978 | 23.616038 | -99.987770 | -41.811406 | -18.391057 | -1.486615 | 0.512622 | 2.406045 | 12.412710 |
| bic_diff_relative | 50.0 | -0.000463 | 0.000883 | -0.003927 | -0.001460 | -0.000498 | -0.000046 | 0.000019 | 0.000060 | 0.000462 |
| aic_diff | 50.0 | -17.985094 | 32.798093 | -134.374300 | -61.247563 | -20.231837 | -2.880080 | 0.497407 | 1.461592 | 4.538230 |
| aic_diff_relative | 50.0 | -0.000644 | 0.001245 | -0.005387 | -0.001730 | -0.000665 | -0.000087 | 0.000018 | 0.000056 | 0.000094 |
| aicc_diff | 50.0 | -17.981811 | 32.791197 | -134.350860 | -61.224113 | -20.231837 | -2.880080 | 0.497408 | 1.461583 | 4.538230 |
| aicc_diff_relative | 50.0 | -0.000644 | 0.001244 | -0.005386 | -0.001729 | -0.000665 | -0.000087 | 0.000018 | 0.000056 | 0.000094 |
| lnL_diff | 50.0 | 9.552546 | 17.591805 | -2.269110 | -0.730796 | -0.248700 | 1.440040 | 10.115917 | 34.623781 | 71.187160 |
| lnL_diff_relative | 50.0 | -0.000687 | 0.001339 | -0.005735 | -0.001926 | -0.000668 | -0.000088 | 0.000018 | 0.000056 | 0.000095 |
| unrooted_softwired_network_distance | 50.0 | 0.072628 | 0.091584 | 0.000000 | 0.000000 | 0.000000 | 0.045549 | 0.105263 | 0.192262 | 0.363636 |
| unrooted_hardwired_network_distance | 50.0 | 0.136864 | 0.117976 | 0.000000 | 0.000000 | 0.055556 | 0.105263 | 0.189474 | 0.253571 | 0.520000 |
| unrooted_displayed_trees_distance | 50.0 | 1.000000 | 0.000000 | 1.000000 | 1.000000 | 1.000000 | 1.000000 | 1.000000 | 1.000000 | 1.000000 |
| rooted_softwired_network_distance | 50.0 | 0.338254 | 0.129914 | 0.050000 | 0.199231 | 0.232601 | 0.346990 | 0.407407 | 0.501724 | 0.657143 |
| rooted_hardwired_network_distance | 50.0 | 0.316696 | 0.125050 | 0.052632 | 0.150000 | 0.238095 | 0.318182 | 0.391304 | 0.502000 | 0.615385 |
| rooted_displayed_trees_distance | 50.0 | 1.000000 | 0.000000 | 1.000000 | 1.000000 | 1.000000 | 1.000000 | 1.000000 | 1.000000 | 1.000000 |
| rooted_tripartition_distance | 50.0 | 0.335528 | 0.129873 | 0.052632 | 0.190476 | 0.246753 | 0.347826 | 0.410326 | 0.520000 | 0.678571 |
| rooted_path_multiplicity_distance | 50.0 | 0.188309 | 0.079139 | 0.048780 | 0.095238 | 0.139535 | 0.181818 | 0.238889 | 0.297872 | 0.420000 |
| rooted_nested_labels_distance | 50.0 | 0.453898 | 0.173864 | 0.095238 | 0.260870 | 0.333333 | 0.461538 | 0.571429 | 0.666667 | 0.882353 |
| runtime_raxml | 50.0 | 8.266320 | 2.342121 | 5.007000 | 5.937400 | 6.569500 | 7.582500 | 9.378250 | 11.928000 | 14.984000 |
| runtime_inference | 50.0 | 19.466400 | 13.443946 | 2.505000 | 2.842900 | 15.182000 | 18.130500 | 22.896750 | 27.263200 | 93.665000 |

| | count | mean | std | min | 10% | 25% | 50% | 75% | 90% | max |
|---|---|---|---|---|---|---|---|---|---|---|
| n_reticulations_inferred | 50.0 | 0.960000 | 0.197949 | 0.000000 | 1.000000 | 1.000000 | 1.000000 | 1.000000 | 1.000000 | 1.000000 |
| bic_diff | 50.0 | -7.467701 | 15.375576 | -64.636350 | -26.826223 | -8.393400 | -0.300040 | 0.612675 | 2.406045 | 12.412710 |
| bic_diff_relative | 50.0 | -0.000259 | 0.000542 | -0.002531 | -0.000910 | -0.000323 | -0.000009 | 0.000024 | 0.000060 | 0.000462 |
| aic_diff | 50.0 | -8.843163 | 18.305075 | -99.022900 | -26.826223 | -13.249473 | -0.692325 | 0.594030 | 1.673182 | 4.538230 |
| aic_diff_relative | 50.0 | -0.000318 | 0.000683 | -0.003957 | -0.000926 | -0.000446 | -0.000019 | 0.000024 | 0.000056 | 0.000094 |
| aicc_diff | 50.0 | -8.842225 | 18.302373 | -98.999440 | -26.826224 | -13.249473 | -0.692325 | 0.594032 | 1.673182 | 4.538230 |
| aicc_diff_relative | 50.0 | -0.000318 | 0.000682 | -0.003956 | -0.000926 | -0.000446 | -0.000019 | 0.000024 | 0.000056 | 0.000094 |
| lnL_diff | 50.0 | 4.581581 | 9.634799 | -2.269110 | -0.836591 | -0.297020 | 0.346160 | 6.624737 | 15.173278 | 53.511440 |
| lnL_diff_relative | 50.0 | -0.000332 | 0.000727 | -0.004297 | -0.001159 | -0.000448 | -0.000019 | 0.000024 | 0.000056 | 0.000095 |
| unrooted_softwired_network_distance | 50.0 | 0.063774 | 0.081647 | 0.000000 | 0.000000 | 0.000000 | 0.020833 | 0.105263 | 0.161153 | 0.320000 |
| unrooted_hardwired_network_distance | 50.0 | 0.135450 | 0.113924 | 0.000000 | 0.000000 | 0.055556 | 0.105263 | 0.189474 | 0.253571 | 0.520000 |
| unrooted_displayed_trees_distance | 50.0 | 1.000000 | 0.000000 | 1.000000 | 1.000000 | 1.000000 | 1.000000 | 1.000000 | 1.000000 | 1.000000 |
| rooted_softwired_network_distance | 50.0 | 0.329381 | 0.124206 | 0.050000 | 0.199231 | 0.232601 | 0.346154 | 0.404040 | 0.501724 | 0.657143 |
| rooted_hardwired_network_distance | 50.0 | 0.312127 | 0.119076 | 0.052632 | 0.186429 | 0.238095 | 0.285714 | 0.384387 | 0.502000 | 0.560000 |
| rooted_displayed_trees_distance | 50.0 | 1.000000 | 0.000000 | 1.000000 | 1.000000 | 1.000000 | 1.000000 | 1.000000 | 1.000000 | 1.000000 |
| rooted_tripartition_distance | 50.0 | 0.322403 | 0.124334 | 0.052632 | 0.190476 | 0.238095 | 0.295455 | 0.391304 | 0.520000 | 0.592593 |
| rooted_path_multiplicity_distance | 50.0 | 0.175805 | 0.071805 | 0.048780 | 0.095238 | 0.139535 | 0.160677 | 0.222222 | 0.297872 | 0.367347 |
| rooted_nested_labels_distance | 50.0 | 0.431047 | 0.167845 | 0.095238 | 0.260870 | 0.278986 | 0.400000 | 0.518519 | 0.666667 | 0.857143 |
| runtime_raxml | 50.0 | 8.266320 | 2.342121 | 5.007000 | 5.937400 | 6.569500 | 7.582500 | 9.378250 | 11.928000 | 14.984000 |
| runtime_inference | 50.0 | 17.131180 | 10.853892 | 2.150000 | 11.426200 | 12.523000 | 15.732500 | 19.468500 | 23.028600 | 82.427000 |

**Table 25.** Percentiles for experiment B, reticulation probability 0.4, starting from RAxML-NG best tree. Top: `LhType.AVERAGE`, bottom: `LhType.BEST`.

##### 6.12 B: Reticulation probability 0.5

We simulated 50 data sets with 20 taxa and 1 reticulation each. We varied the first-parent probability of the simulated reticulation to be in $\{0.1, 0.2, 0.3, 0.4, 0.5\}$. Here, we present the plots and tables for reticulation probability 0.5.

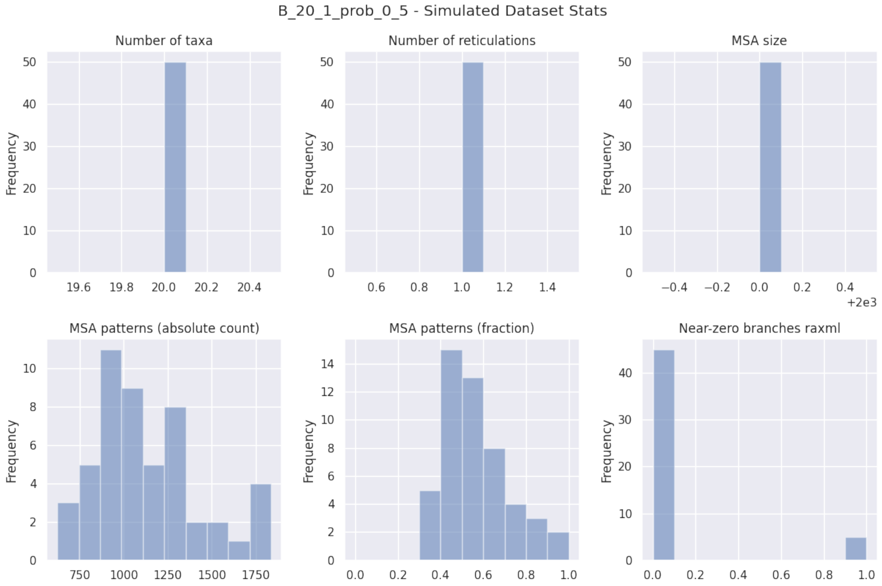


**Fig. 58.** Simulated dataset statistics for experiment B, reticulation probability 0.5.

| B_20_1_prob_0.5_norandom | LhType.AVERAGE | LhType.BEST |
|---|---|---|
| Inferred BIC better or equal | 21 (42.00 %) | 23 (46.00 %) |
| Inferred AIC better or equal | 21 (42.00 %) | 23 (46.00 %) |
| Inferred AICc better or equal | 21 (42.00 %) | 23 (46.00 %) |
| Inferred BIC worse | 29 (58.00 %) | 27 (54.00 %) |
| Inferred AIC worse | 29 (58.00 %) | 27 (54.00 %) |
| Inferred AICc worse | 29 (58.00 %) | 27 (54.00 %) |
| Inferred lnL better or equal | 21 (42.00 %) | 23 (46.00 %) |
| Inferred lnL worse | 29 (58.00 %) | 27 (54.00 %) |
| Inferred n_reticulations less | 3 (6.00 %) | 0 (0.00 %) |
| Inferred n_reticulations equal | 47 (94.00 %) | 50 (100.00 %) |
| Inferred n_reticulations more | 0 (0.00 %) | 0 (0.00 %) |
| Unrooted softwired distance zero | 19 (38.00 %) | 21 (42.00 %) |
| Good result | 34 (68.00 %) | 38 (76.00 %) |
| Passable result | 13 (26.00 %) | 12 (24.00 %) |
| Bad result | 3 (6.00 %) | 0 (0.00 %) |

**Table 26.** Summary statistics for experiment B, reticulation probability 0.5, starting from RAxML-NG best tree.

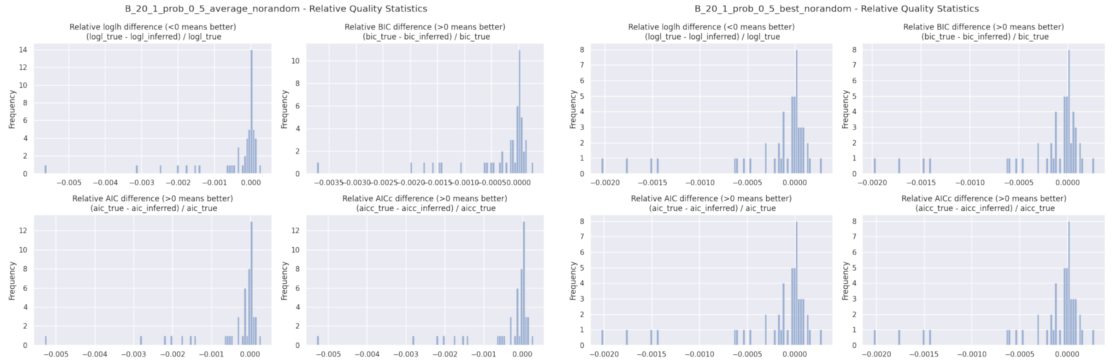


**Fig. 59.** Relative BIC, AIC, AICc, and loglikelihood differences for experiment B, reticulation probability 0.5, starting from RAxML-NG best tree. Left: `LhType.AVERAGE`, right: `LhType.BEST`.

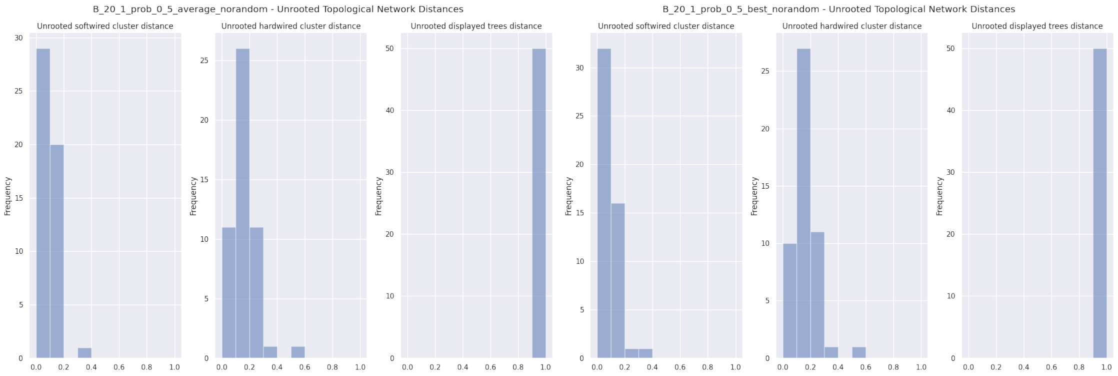


**Fig. 60.** Unrooted relative distances for experiment B, reticulation probability 0.5, starting from RAxML-NG best tree. Left: `LhType.AVERAGE`, right: `LhType.BEST`.

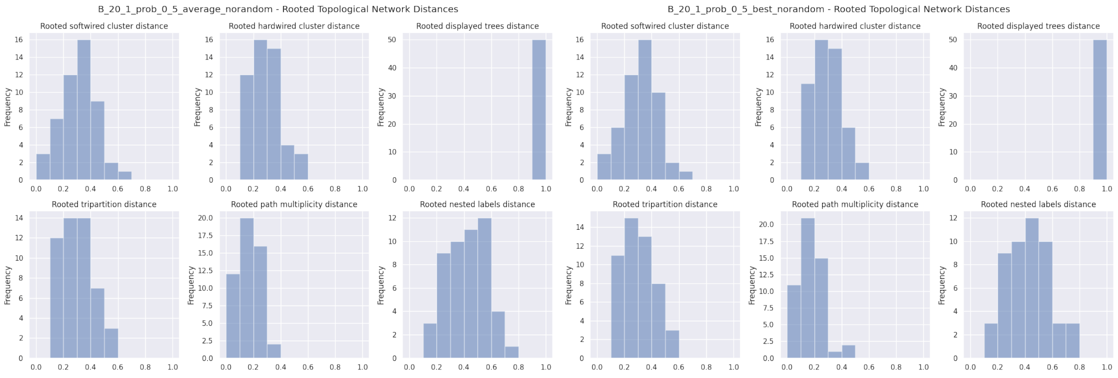


**Fig. 61.** Rooted relative distances for experiment B, reticulation probability 0.5, starting from RAxML-NG best tree. Left: `LhType.AVERAGE`, right: `LhType.BEST`.

| | count | mean | std | min | 10% | 25% | 50% | 75% | 90% | max |
|---|---|---|---|---|---|---|---|---|---|---|
| n_reticulations_inferred | 50.0 | 0.940000 | 0.239898 | 0.000000 | 1.000000 | 1.000000 | 1.000000 | 1.000000 | 1.000000 | 1.000000 |
| bic_diff | 50.0 | -9.213215 | 18.870128 | -88.017760 | -38.910278 | -9.618305 | -0.694400 | 0.727093 | 2.900803 | 6.453050 |
| bic_diff_relative | 50.0 | -0.000321 | 0.000718 | -0.003723 | -0.001415 | -0.000313 | -0.000027 | 0.000021 | 0.000086 | 0.000274 |
| aic_diff | 50.0 | -11.276409 | 24.890811 | -122.404290 | -42.161782 | -9.618305 | -0.694400 | 0.727082 | 2.900802 | 6.453050 |
| aic_diff_relative | 50.0 | -0.000404 | 0.000970 | -0.005293 | -0.001536 | -0.000319 | -0.000027 | 0.000021 | 0.000087 | 0.000280 |
| aicc_diff | 50.0 | -11.275002 | 24.886269 | -122.380840 | -42.161782 | -9.618312 | -0.694405 | 0.727090 | 2.900803 | 6.453040 |
| aicc_diff_relative | 50.0 | -0.000404 | 0.000970 | -0.005292 | -0.001536 | -0.000319 | -0.000027 | 0.000021 | 0.000087 | 0.000280 |
| lnL_diff | 50.0 | 5.878205 | 13.232194 | -3.226530 | -1.450401 | -0.363547 | 0.347195 | 4.809150 | 21.080886 | 65.202150 |
| lnL_diff_relative | 50.0 | -0.000423 | 0.001036 | -0.005668 | -0.001543 | -0.000320 | -0.000028 | 0.000021 | 0.000087 | 0.000282 |
| unrooted_softwired_network_distance | 50.0 | 0.074161 | 0.069974 | 0.000000 | 0.000000 | 0.000000 | 0.090909 | 0.105263 | 0.144361 | 0.320000 |
| unrooted_hardwired_network_distance | 50.0 | 0.138405 | 0.094587 | 0.000000 | 0.000000 | 0.105263 | 0.111111 | 0.197619 | 0.238095 | 0.500000 |
| unrooted_displayed_trees_distance | 50.0 | 1.000000 | 0.000000 | 1.000000 | 1.000000 | 1.000000 | 1.000000 | 1.000000 | 1.000000 | 1.000000 |
| rooted_softwired_network_distance | 50.0 | 0.312897 | 0.124577 | 0.086957 | 0.129891 | 0.228147 | 0.338542 | 0.393281 | 0.426603 | 0.666667 |
| rooted_hardwired_network_distance | 50.0 | 0.290958 | 0.118050 | 0.100000 | 0.145526 | 0.200000 | 0.272727 | 0.391304 | 0.420833 | 0.576923 |
| rooted_displayed_trees_distance | 50.0 | 1.000000 | 0.000000 | 1.000000 | 1.000000 | 1.000000 | 1.000000 | 1.000000 | 1.000000 | 1.000000 |
| rooted_tripartition_distance | 50.0 | 0.301525 | 0.124160 | 0.100000 | 0.145000 | 0.209524 | 0.272727 | 0.391304 | 0.458333 | 0.576923 |
| rooted_path_multiplicity_distance | 50.0 | 0.167923 | 0.072613 | 0.048780 | 0.090592 | 0.139535 | 0.139535 | 0.222222 | 0.260870 | 0.333333 |
| rooted_nested_labels_distance | 50.0 | 0.419454 | 0.144772 | 0.181818 | 0.260870 | 0.333333 | 0.400000 | 0.518519 | 0.600000 | 0.709677 |
| runtime_raxml | 50.0 | 8.328440 | 2.378372 | 4.951000 | 5.748700 | 6.560250 | 7.759000 | 9.534500 | 11.169300 | 14.527000 |
| runtime_inference | 50.0 | 20.191480 | 6.636571 | 2.507000 | 14.395400 | 16.307000 | 19.990000 | 23.777500 | 28.354400 | 35.420000 |

| | count | mean | std | min | 10% | 25% | 50% | 75% | 90% | max |
|---|---|---|---|---|---|---|---|---|---|---|
| n_reticulations_inferred | 50.0 | 1.000000 | 0.000000 | 1.000000 | 1.000000 | 1.000000 | 1.000000 | 1.000000 | 1.000000 | 1.000000 |
| bic_diff | 50.0 | -5.756791 | 13.574637 | -51.816450 | -21.620362 | -5.171863 | -0.169670 | 0.895948 | 2.623965 | 6.453050 |
| bic_diff_relative | 50.0 | -0.000190 | 0.000475 | -0.001986 | -0.000596 | -0.000154 | -0.000007 | 0.000026 | 0.000084 | 0.000274 |
| aic_diff | 50.0 | -5.756792 | 13.574637 | -51.816450 | -21.620362 | -5.171862 | -0.169675 | 0.895945 | 2.623964 | 6.453040 |
| aic_diff_relative | 50.0 | -0.000194 | 0.000484 | -0.002026 | -0.000605 | -0.000156 | -0.000007 | 0.000027 | 0.000085 | 0.000280 |
| aicc_diff | 50.0 | -5.756792 | 13.574637 | -51.816450 | -21.620353 | -5.171862 | -0.169675 | 0.895948 | 2.623964 | 6.453040 |
| aicc_diff_relative | 50.0 | -0.000194 | 0.000484 | -0.002026 | -0.000605 | -0.000156 | -0.000007 | 0.000027 | 0.000085 | 0.000280 |
| lnL_diff | 50.0 | 2.878396 | 6.787318 | -3.226530 | -1.311983 | -0.447977 | 0.084835 | 2.585930 | 10.810185 | 25.908230 |
| lnL_diff_relative | 50.0 | -0.000195 | 0.000486 | -0.002036 | -0.000607 | -0.000157 | -0.000007 | 0.000027 | 0.000085 | 0.000282 |
| unrooted_softwired_network_distance | 50.0 | 0.069414 | 0.072573 | 0.000000 | 0.000000 | 0.000000 | 0.086957 | 0.105263 | 0.145238 | 0.320000 |
| unrooted_hardwired_network_distance | 50.0 | 0.142223 | 0.095075 | 0.000000 | 0.000000 | 0.105263 | 0.150000 | 0.197619 | 0.238095 | 0.500000 |
| unrooted_displayed_trees_distance | 50.0 | 1.000000 | 0.000000 | 1.000000 | 1.000000 | 1.000000 | 1.000000 | 1.000000 | 1.000000 | 1.000000 |
| rooted_softwired_network_distance | 50.0 | 0.318253 | 0.123122 | 0.086957 | 0.141071 | 0.235577 | 0.333333 | 0.398485 | 0.427436 | 0.666667 |
| rooted_hardwired_network_distance | 50.0 | 0.297994 | 0.115428 | 0.100000 | 0.150000 | 0.209524 | 0.272727 | 0.391304 | 0.418478 | 0.576923 |
| rooted_displayed_trees_distance | 50.0 | 1.000000 | 0.000000 | 1.000000 | 1.000000 | 1.000000 | 1.000000 | 1.000000 | 1.000000 | 1.000000 |
| rooted_tripartition_distance | 50.0 | 0.306613 | 0.121518 | 0.100000 | 0.150000 | 0.238095 | 0.272727 | 0.391304 | 0.460500 | 0.576923 |
| rooted_path_multiplicity_distance | 50.0 | 0.173934 | 0.081488 | 0.048780 | 0.095238 | 0.139535 | 0.139535 | 0.222222 | 0.260870 | 0.431373 |
| rooted_nested_labels_distance | 50.0 | 0.420644 | 0.148776 | 0.181818 | 0.260870 | 0.333333 | 0.400000 | 0.518519 | 0.666667 | 0.750000 |
| runtime_raxml | 50.0 | 8.328440 | 2.378372 | 4.951000 | 5.748700 | 6.560250 | 7.759000 | 9.534500 | 11.169300 | 14.527000 |
| runtime_inference | 50.0 | 17.444160 | 5.078269 | 8.812000 | 11.698700 | 13.056000 | 16.509500 | 20.467500 | 24.225300 | 31.236000 |

**Table 27.** Percentiles for experiment B, reticulation probability 0.5, starting from RAxML-NG best tree. Top: `LhType.AVERAGE`, bottom: `LhType.BEST`.

#### 6.13 C: Unpartitioned Data

In this experiment, we simulated 50 data sets with 20 taxa and 1 reticulation. In addition to normal inference, we started a second inference where we merged all simulated partitions into a single partition before running the inference.

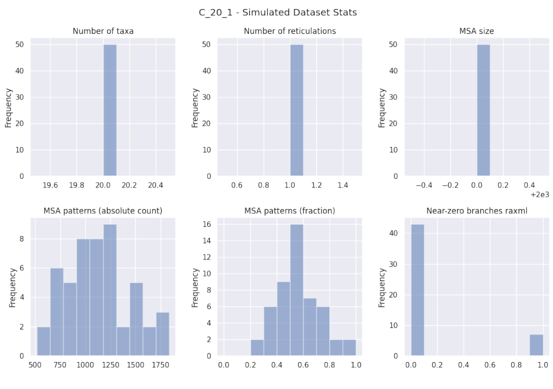


**Fig. 62.** Simulated dataset statistics for experiment C.

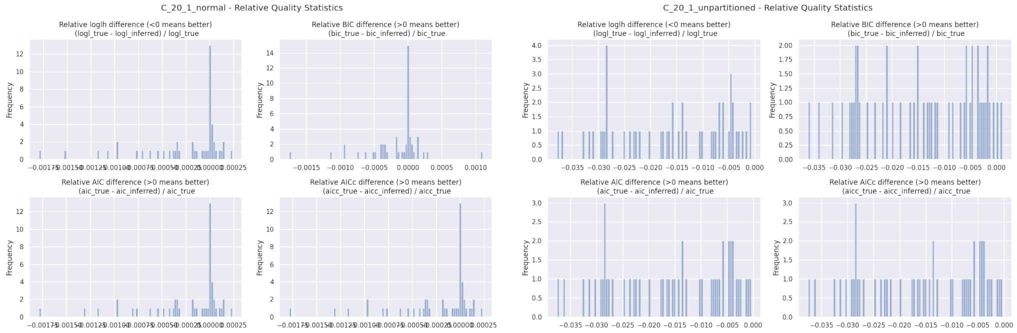


**Fig. 63.** Relative BIC, AIC, AICc, and loglikelihood differences for experiment C, `LhModel.AVERAGE`, starting from RAxML-NG best tree. Left: partitioned, right: unpartitioned.

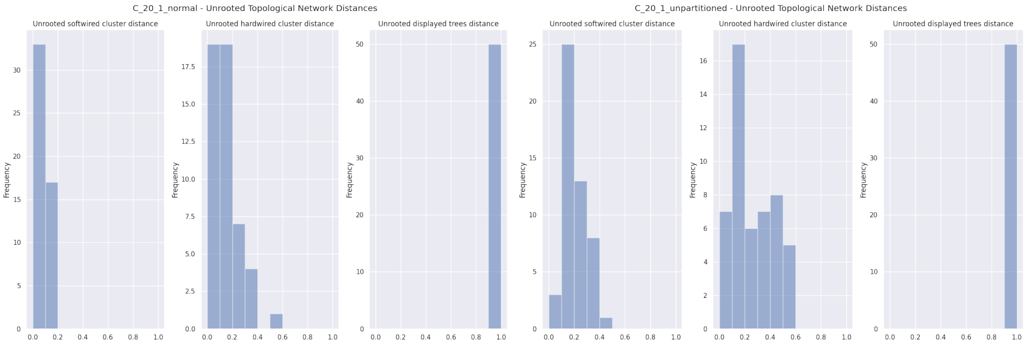


**Fig. 64.** Unrooted relative distances for experiment C, `LhModel.AVERAGE`, starting from RAxML-NG best tree. Left: partitioned, right: unpartitioned.

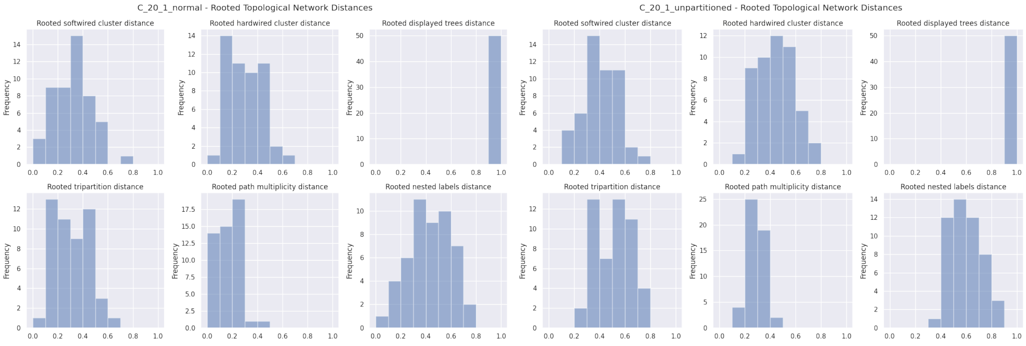


**Fig. 65.** Rooted relative distances for experiment C, `LhModel.AVERAGE`, starting from RAxML-NG best tree. Left: partitioned, right: unpartitioned.

| C_20_1_normal | LhType.AVERAGE |
|---|---|
| Inferred BIC better or equal | 23 (46.00 %) |
| Inferred AIC better or equal | 21 (42.00 %) |
| Inferred AICc better or equal | 21 (42.00 %) |
| Inferred BIC worse | 27 (54.00 %) |
| Inferred AIC worse | 29 (58.00 %) |
| Inferred AICc worse | 29 (58.00 %) |
| Inferred lnL better or equal | 21 (42.00 %) |
| Inferred lnL worse | 29 (58.00 %) |
| Inferred n_reticulations less | 3 (6.00 %) |
| Inferred n_reticulations equal | 47 (94.00 %) |
| Inferred n_reticulations more | 0 (0.00 %) |
| Unrooted softwired distance zero | 24 (48.00 %) |
| Good result | 39 (78.00 %) |
| Passable result | 10 (20.00 %) |
| Bad result | 1 (2.00 %) |

| C_20_1_unpartitioned | LhType.AVERAGE |
|---|---|
| Inferred BIC better or equal | 2 (4.00 %) |
| Inferred AIC better or equal | 0 (0.00 %) |
| Inferred AICc better or equal | 0 (0.00 %) |
| Inferred BIC worse | 48 (96.00 %) |
| Inferred AIC worse | 50 (100.00 %) |
| Inferred AICc worse | 50 (100.00 %) |
| Inferred lnL better or equal | 0 (0.00 %) |
| Inferred lnL worse | 50 (100.00 %) |
| Inferred n_reticulations less | 50 (100.00 %) |
| Inferred n_reticulations equal | 0 (0.00 %) |
| Inferred n_reticulations more | 0 (0.00 %) |
| Unrooted softwired distance zero | 0 (0.00 %) |
| Good result | 2 (4.00 %) |
| Passable result | 0 (0.00 %) |
| Bad result | 48 (96.00 %) |

**Table 28.** Summary statistics for experiment C, `LhModel.AVERAGE`, starting from RAxML-NG best tree. Left: partitioned data, right: unpartitioned data.

| | count | mean | std | min | 10% | 25% | 50% | 75% | 90% | max |
|---|---|---|---|---|---|---|---|---|---|---|
| n_reticulations_inferred | 50.0 | 0.940000 | 0.239898 | 0.000000 | 1.000000 | 1.000000 | 1.000000 | 1.000000 | 1.000000 | 1.000000 |
| bic_diff | 50.0 | -5.342807 | 12.323147 | -47.607100 | -24.437276 | -10.196735 | -0.057580 | 0.455475 | 3.563545 | 25.964170 |
| bic_diff_relative | 50.0 | -0.000164 | 0.000422 | -0.001752 | -0.000630 | -0.000334 | -0.000002 | 0.000014 | 0.000131 | 0.001098 |
| aic_diff | 50.0 | -7.406000 | 13.040210 | -49.946830 | -24.928723 | -10.540188 | -0.443360 | 0.348975 | 1.569980 | 5.342310 |
| aic_diff_relative | 50.0 | -0.000236 | 0.000420 | -0.001786 | -0.000822 | -0.000367 | -0.000013 | 0.000009 | 0.000059 | 0.000234 |
| aicc_diff | 50.0 | -7.404593 | 13.037956 | -49.923380 | -24.928724 | -10.540187 | -0.443360 | 0.348975 | 1.569980 | 5.342310 |
| aicc_diff_relative | 50.0 | -0.000236 | 0.000420 | -0.001786 | -0.000821 | -0.000367 | -0.000013 | 0.000009 | 0.000059 | 0.000234 |
| lnL_diff | 50.0 | 3.943001 | 6.960485 | -2.671160 | -0.784989 | -0.174483 | 0.221685 | 5.320187 | 12.464371 | 28.973410 |
| lnL_diff_relative | 50.0 | -0.000254 | 0.000446 | -0.001794 | -0.000969 | -0.000387 | -0.000013 | 0.000009 | 0.000060 | 0.000236 |
| unrooted_softwired_network_distance | 50.0 | 0.057302 | 0.065395 | 0.000000 | 0.000000 | 0.000000 | 0.043561 | 0.105263 | 0.160667 | 0.190476 |
| unrooted_hardwired_network_distance | 50.0 | 0.124390 | 0.116981 | 0.000000 | 0.000000 | 0.052632 | 0.105263 | 0.157895 | 0.288961 | 0.560000 |
| unrooted_displayed_trees_distance | 50.0 | 1.000000 | 0.000000 | 1.000000 | 1.000000 | 1.000000 | 1.000000 | 1.000000 | 1.000000 | 1.000000 |
| rooted_softwired_network_distance | 50.0 | 0.318455 | 0.148792 | 0.000000 | 0.135227 | 0.204348 | 0.312937 | 0.432143 | 0.501515 | 0.756098 |
| rooted_hardwired_network_distance | 50.0 | 0.295076 | 0.145960 | 0.000000 | 0.100000 | 0.190476 | 0.272727 | 0.416667 | 0.460500 | 0.678571 |
| rooted_displayed_trees_distance | 50.0 | 1.000000 | 0.000000 | 1.000000 | 1.000000 | 1.000000 | 1.000000 | 1.000000 | 1.000000 | 1.000000 |
| rooted_tripartition_distance | 50.0 | 0.309866 | 0.149409 | 0.000000 | 0.100000 | 0.190476 | 0.295455 | 0.416667 | 0.480000 | 0.689655 |
| rooted_path_multiplicity_distance | 50.0 | 0.170874 | 0.086571 | 0.000000 | 0.048780 | 0.095238 | 0.181818 | 0.222222 | 0.263043 | 0.400000 |
| rooted_nested_labels_distance | 50.0 | 0.432023 | 0.162355 | 0.095238 | 0.252964 | 0.333333 | 0.461538 | 0.571429 | 0.625287 | 0.750000 |
| runtime_raxml | 50.0 | 19.701100 | 6.737171 | 8.886000 | 12.700100 | 14.749250 | 18.828500 | 23.186500 | 27.760800 | 40.611000 |
| runtime_inference | 50.0 | 14.339380 | 5.102766 | 1.804000 | 8.965400 | 12.005500 | 14.658000 | 17.096750 | 19.133400 | 28.418000 |

| | count | mean | std | min | 10% | 25% | 50% | 75% | 90% | max |
|---|---|---|---|---|---|---|---|---|---|---|
| n_reticulations_inferred | 50.0 | 0.000000 | 0.000000 | 0.000000 | 0.000000 | 0.000000 | 0.000000 | 0.000000 | 0.000000 | 0.000000 |
| bic_diff | 50.0 | -461.512353 | 390.903511 | -1435.995710 | -917.150004 | -641.332433 | -374.055500 | -134.652185 | -45.274598 | 25.964150 |
| bic_diff_relative | 50.0 | -0.014295 | 0.010733 | -0.036166 | -0.027920 | -0.024484 | -0.013073 | -0.004618 | -0.001687 | 0.001098 |
| aic_diff | 50.0 | -495.898893 | 390.903509 | -1470.382240 | -951.536545 | -675.718965 | -408.442040 | -169.038725 | -79.661137 | -8.422400 |
| aic_diff_relative | 50.0 | -0.015707 | 0.010894 | -0.037550 | -0.029150 | -0.026378 | -0.014814 | -0.005757 | -0.003069 | -0.000364 |
| aicc_diff | 50.0 | -495.875441 | 390.903511 | -1470.358790 | -951.513104 | -675.695523 | -408.418590 | -169.015267 | -79.637687 | -8.398950 |
| aicc_diff_relative | 50.0 | -0.015706 | 0.010894 | -0.037549 | -0.029149 | -0.026377 | -0.014813 | -0.005756 | -0.003068 | -0.000363 |
| lnL_diff | 50.0 | 251.949446 | 195.451755 | 8.211200 | 43.830569 | 88.519363 | 208.221020 | 341.859488 | 479.768272 | 739.191130 |
| lnL_diff_relative | 50.0 | -0.016042 | 0.010933 | -0.037877 | -0.029555 | -0.026831 | -0.015230 | -0.006027 | -0.003401 | -0.000714 |
| unrooted_softwired_network_distance | 50.0 | 0.197691 | 0.097319 | 0.055556 | 0.105263 | 0.105263 | 0.190476 | 0.272727 | 0.333333 | 0.461538 |
| unrooted_hardwired_network_distance | 50.0 | 0.253025 | 0.156754 | 0.000000 | 0.055556 | 0.157895 | 0.205263 | 0.390152 | 0.480435 | 0.560000 |
| unrooted_displayed_trees_distance | 50.0 | 1.000000 | 0.000000 | 1.000000 | 1.000000 | 1.000000 | 1.000000 | 1.000000 | 1.000000 | 1.000000 |
| rooted_softwired_network_distance | 50.0 | 0.404708 | 0.130883 | 0.150000 | 0.269264 | 0.318182 | 0.392308 | 0.500000 | 0.552107 | 0.742857 |
| rooted_hardwired_network_distance | 50.0 | 0.435278 | 0.149328 | 0.105263 | 0.238095 | 0.318182 | 0.458333 | 0.572692 | 0.629630 | 0.724138 |
| rooted_displayed_trees_distance | 50.0 | 1.000000 | 0.000000 | 1.000000 | 1.000000 | 1.000000 | 1.000000 | 1.000000 | 1.000000 | 1.000000 |
| rooted_tripartition_distance | 50.0 | 0.507565 | 0.138597 | 0.238095 | 0.318182 | 0.391304 | 0.520000 | 0.629630 | 0.678571 | 0.766667 |
| rooted_path_multiplicity_distance | 50.0 | 0.289772 | 0.074778 | 0.162791 | 0.204545 | 0.244444 | 0.282609 | 0.354167 | 0.387755 | 0.450980 |
| rooted_nested_labels_distance | 50.0 | 0.599436 | 0.128388 | 0.347826 | 0.416667 | 0.494615 | 0.592593 | 0.689655 | 0.733333 | 0.882353 |
| runtime_raxml | 50.0 | 19.701100 | 6.737171 | 8.886000 | 12.700100 | 14.749250 | 18.828500 | 23.186500 | 27.760800 | 40.611000 |
| runtime_inference | 50.0 | 2.000220 | 0.322907 | 1.131000 | 1.624100 | 1.808250 | 2.017500 | 2.157000 | 2.405500 | 2.840000 |

**Table 29.** Percentiles for experiment C, starting from RAxML-NG best tree. Top: partitioned, bottom: unpartitioned.

##### 6.14 F: Parallel Scalability

**F1: Small MSA** We simulated 10 networks with 20 taxa and 3 reticulations each. We simulated $10,000$ MSA sites per displayed tree, resulting in a MSA with $80,000$ sites. We started the NetRAX inferences from RAxML-NG ML trees, using $\{1, 2, 4, 8, 16, 32, 64\}$ MPI processes.

The experiment ran on up to 4 in-house cluster compute nodes with Intel CPUs (E5-2630v3 with 20 MB cache, running at 2.40GHz). Each compute node has 2 CPUs with 8 physical cores each and 64 GB RAM.

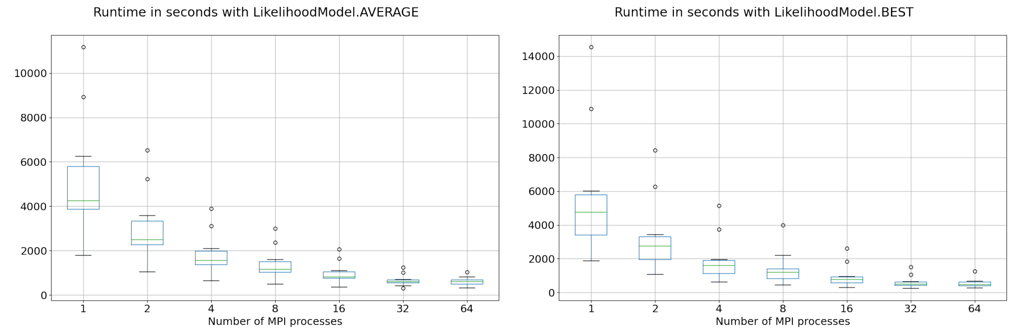


**Fig. 66.** Left: NetRAX inference runtime in seconds starting from a single RAxML-NG ML tree, over 10 different datasets. Left: `LhModel.AVERAGE`, right: `LhModel.BEST`.

#### 7 Detailed Experimental Results on Empirical Data

##### 7.1 Snake Genomes

| empirical_snakes | LhType.BEST single | LhType.AVERAGE single | LhType.BEST multi | LhType.AVERAGE multi |
|---|---|---|---|---|
| Inferred lnL | -727228.5853 | -727145.753 | -726893.6198 | -727209.282 |
| Inferred BIC | 1499442.358 | 1499276.693 | 1498837.061 | 1499403.751 |
| Relative BIC difference | 0.0001875409309 | 9.354290602e-05 | 0.000658713729 | 8.108553613e-05 |
| Inferred BIC LhType.BEST | 1499442.358 | 1499356.751 | 1498837.061 | 1499375.126 |
| Inferred BIC LhType.AVERAGE | 1499275.623 | 1499276.693 | 1498548.965 | 1499403.751 |
| Inferred n_reticulations | 1 | 1 | 2 | 1 |
| Unrooted softwired network distance | 0.04347826087 | 0.04347826087 | 0.25 | 0.04347826087 |
| Unrooted hardwired network distance | 0.09090909091 | 0.09090909091 | 0.3461538462 | 0.04545454545 |
| Unrooted displayed trees distance | 1 | 1 | 1 | 1 |
| Rooted softwired network distance | 0.3333333333 | 0.3333333333 | 0.40625 | 0.2592592593 |
| Rooted hardwired network distance | 0.3846153846 | 0.3846153846 | 0.3703703704 | 0.28 |
| Rooted displayed trees distance | 1 | 1 | 1 | 1 |
| Rooted tripartition distance | 0.3846153846 | 0.3846153846 | 0.4827586207 | 0.28 |
| Rooted path multiplicity distance | 0.2307692308 | 0.2307692308 | 0.2452830189 | 0.16 |
| Rooted nested labels distance | 0.5625 | 0.5625 | 0.5454545455 | 0.3571428571 |
| NetRAX inference runtime (seconds) | 193 | 844 | 4182 | 15615 |

**Table 30.** NetRAX results for the empirical snakes dataset, for `LhType.BEST` and `LhType.AVERAGE`, compared to the 1-reticulation network inferred by SNAQ. The term `single` refers to starting from the RAxML-NG best ML tree. The term `multi` refers to starting from 14 unique tree topologies out of 10 random and 10 RAxML-NG maximum parsimony trees. In all cases, NetRAX inferred a better BIC score (but we used our model for computing BIC).

#### References

1. Daniel H Huson, Regula Rupp, and Celine Scornavacca. *Phylogenetic networks: concepts, algorithms and applications*. Cambridge University Press, 2010.

2. Asanka Perera. Finding the optimal number of clusters for k-means through elbow method using a mathematical approach compared to graphical approach. https://www.linkedin.com/pulse/finding-optimal-number-clusters-k-means-through-elbow-asanka-perera/, 2017. Website. Accessed July 28, 2021.
3. Robert L Thorndike. Who belongs in the family? *Psychometrika*, 18(4):267–276, 1953.
